# White matter abnormalities across different epilepsy syndromes in adults: an ENIGMA Epilepsy study

**DOI:** 10.1101/2019.12.19.883405

**Authors:** Sean N Hatton, Khoa H Huynh, Leonardo Bonilha, Eugenio Abela, Saud Alhusaini, Andre Altmann, Marina KM Alvim, Akshara R Balachandra, Emanuele Bartolini, Benjamin Bender, Neda Bernasconi, Andrea Bernasconi, Boris Bernhardt, Núria Bargallo, Benoit Caldairou, Maria Eugenia Caligiuri, Sarah JA Carr, Gianpiero L Cavalleri, Fernando Cendes, Luis Concha, Esmaeil Davoodi-bojd, Patricia M Desmond, Orrin Devinsky, Colin P Doherty, Martin Domin, John S Duncan, Niels K Focke, Sonya F Foley, Antonio Gambardella, Ezequiel Gleichgerrcht, Renzo Guerrini, Khalid Hamandi, Akaria Ishikawa, Simon S Keller, Peter V Kochunov, Raviteja Kotikalapudi, Barbara AK Kreilkamp, Patrick Kwan, Angelo Labate, Soenke Langner, Matteo Lenge, Min Liu, Elaine Lui, Pascal Martin, Mario Mascalchi, José CV Moreira, Marcia E Morita-Sherman, Terence J O’Brien, Heath R Pardoe, José C Pariente, Letícia F Ribeiro, Mark P Richardson, Cristiane S Rocha, Raúl Rodríguez-Cruces, Felix Rosenow, Mariasavina Severino, Benjamin Sinclair, Hamid Soltanian-Zadeh, Pasquale Striano, Peter N Taylor, Rhys H Thomas, Domenico Tortora, Dennis Velakoulis, Annamaria Vezzani, Lucy Vivash, Felix von Podewils, Sjoerd B Vos, Bernd Weber, Gavin P Winston, Clarissa L Yasuda, Paul M Thompson, Neda Jahanshad, Sanjay M Sisodiya, Carrie R McDonald

## Abstract

The epilepsies are commonly accompanied by widespread abnormalities in cerebral white matter. ENIGMA-Epilepsy is a large quantitative brain imaging consortium, aggregating data to investigate patterns of neuroimaging abnormalities in common epilepsy syndromes, including temporal lobe epilepsy, extratemporal epilepsy, and genetic generalized epilepsy. Our goal was to rank the most robust white matter microstructural differences across and within syndromes in a multicentre sample of adult epilepsy patients. Diffusion-weighted MRI data were analyzed from 1,069 non-epileptic controls and 1,249 patients: temporal lobe epilepsy with hippocampal sclerosis (N=599), temporal lobe epilepsy with normal MRI (N=275), genetic generalized epilepsy (N=182) and nonlesional extratemporal epilepsy (N=193). A harmonized protocol using tract-based spatial statistics was used to derive skeletonized maps of fractional anisotropy and mean diffusivity for each participant, and fiber tracts were segmented using a diffusion MRI atlas. Data were harmonized to correct for scanner-specific variations in diffusion measures using a batch-effect correction tool (ComBat). Analyses of covariance, adjusting for age and sex, examined differences between each epilepsy syndrome and controls for each white matter tract (Bonferroni corrected at p<0.001). Across *“all epilepsies”* lower fractional anisotropy was observed in most fiber tracts with small to medium effect sizes, especially in the corpus callosum, cingulum and external capsule. Less robust effects were seen with mean diffusivity. Syndrome-specific fractional anisotropy and mean diffusivity differences were most pronounced in patients with hippocampal sclerosis in the ipsilateral parahippocampal cingulum and external capsule, with smaller effects across most other tracts. Those with temporal lobe epilepsy and normal MRI showed a similar pattern of greater ipsilateral than contralateral abnormalities, but less marked than those in patients with hippocampal sclerosis. Patients with generalized and extratemporal epilepsies had pronounced differences in fractional anisotropy in the corpus callosum, *corona radiata* and external capsule, and in mean diffusivity of the anterior *corona radiata*. Earlier age of seizure onset and longer disease duration were associated with a greater extent of microstructural abnormalities in patients with hippocampal sclerosis. We demonstrate microstructural abnormalities across major association, commissural, and projection fibers in a large multicentre study of epilepsy. Overall, epilepsy patients showed white matter abnormalities in the corpus callosum, cingulum and external capsule, with differing severity across epilepsy syndromes. These data further define the spectrum of white matter abnormalities in common epilepsy syndromes, yielding new insights into pathological substrates that may be used to guide future therapeutic and genetic studies.

## Introduction

Epilepsy affects over 50 million people worldwide (Bell *et al*., 2014). Focal epilepsies account for around 60% of all adult epilepsy cases, and temporal lobe epilepsy (TLE) is the most common (Téllez-Zenteno and Hernández-Ronquillo, 2012). TLE is associated with hippocampal sclerosis (HS) in 60-70% of cases (Coan and Cendes, 2013). Among adult epilepsy patients, up to 20% have genetic generalized epilepsy (GGE), with bilateral synchronous seizure onset and a presumed genetic background (Scheffer *et al*., 2017). These epilepsy syndromes are frequently studied in isolation and may have distinct pathophysiological substrates and mechanisms. Their unique *and* shared biological pathways are beginning to be unraveled using population genetics (International League Against Epilepsy Consortium on Complex Epilepsies, 2018) and transcriptomics (Altmann *et al*., 2017), paving the pathway for potential novel treatments.

Once considered primarily “gray matter” diseases, brain imaging studies with diffusion magnetic resonance imaging (dMRI) has helped reveal that both focal and generalized epilepsies represent network disorders with widespread white matter alterations even in the absence of visible MRI lesions (Engel *et al*., 2013). Patients with TLE, particularly those with HS, may exhibit white matter abnormalities both proximal to and distant from the seizure focus, often most pronounced in the ipsilateral hemisphere (Focke *et al*., 2008; Ahmadi *et al*., 2009; Labate *et al*., 2015; Caligiuri *et al*., 2016). Studies in patients with GGE have demonstrated microstructural compromises in frontal and parietal regions bilaterally, and in thalamocortical pathways (Keller *et al*., 2011; Lee *et al*., 2014; Szaflarski *et al*., 2016). White matter disruption in epilepsy is also linked to cognitive (McDonald *et al*., 2008; Yogarajah *et al*., 2008, 2010) and postsurgical seizure (Bonilha *et al*., 2015; Keller *et al*., 2015, 2017; Gleichgerrcht *et al*., 2018) outcomes, indicating the importance of white matter networks in the pathophysiology and co-morbidities of epilepsy.

Meta-analyses and single-site studies of dMRI suggest widespread microstructural abnormalities in patients with focal epilepsy affecting association, commissural, and projection fibers, whereas microstructural differences in GGE are reportedly less pronounced (Otte *et al*., 2012; Slinger *et al*., 2016). Unfortunately, the exact tracts, spatial pattern, and extent of damage reported varies across studies, making it hard to draw conclusions about syndrome-specific and generalized white matter pathology in epilepsy. Inconsistencies may be due, in part, to small sample sizes at individual centers, which may lack power to detect reliable differences across a large number of white matter tracts and multiple diffusion measures. Methods for image acquisition, processing, and tract selection also differ greatly across studies, adding other sources of variability. Few studies consider white matter abnormalities as a function of sex, age, and key clinical characteristics leading to multiple uncertainties in the findings. Although meta-analyses reduce some of these limitations, harmonizing the image processing and data analysis in a consortia effort, alleviates some of the known sources of variation and allows for the statistical modeling of other population differences. Furthermore, pooling raw data across a large number of centers in a *mega-analysis^1^* may offer greater power to detect group effects and enable cross-syndrome comparisons that have not previously been possible. Further, due to the collation of worldwide harmonized image analysis protocols, analysis, and reporting of results, white matter differences in epilepsy can now be directly compared to those of other neurological and psychiatric disorders, highlighting pathology that may be unique to epilepsy and/or its treatments.

Enhancing NeuroImaging Genetics through Meta-Analysis (ENIGMA) is a global initiative, combining individually collected samples from studies around the world into a single large-scale study, with coordinated image processing, and integrating imaging, phenotypic, and genomic data from hundreds of research centers worldwide (Thompson *et al*., n.d.). Standardized protocols for image processing, quality assurance, and statistical analyses were applied using the validated ENIGMA-dMRI protocols for multi-site diffusion tensor imaging (DTI) harmonization, http://enigma.usc.edu/ongoing/dti-working-group/ (Jahanshad *et al*., 2013; Kochunov *et al*., 2014, 2015)

Our primary goal was to identify and rank the most robust white matter microstructural alterations across and within common epilepsy syndromes in a sample of 1,249 adult epilepsy patients and 1,069 healthy controls across nine countries from North and South America, Europe and Australia. First, we studied all patients in aggregate (“all epilepsies”) compared to age and sex matched controls, followed by targeted analyses focusing on patients with right and left TLE-HS, right and left non-lesional TLE (TLE-NL), nonlesional extratemporal epilepsy (ExE), and GGE. We characterize effect size (ES) differences across *and* within these epilepsy syndromes in fractional anisotropy (FA) and mean diffusivity (MD), as well as axial (AD) and radial (RD) diffusivity. We also examine regional white matter associations with age of seizure onset and disease duration.

We hypothesized that, compared to controls, each patient group would show white matter alterations beyond the suspected epileptogenic region, with unique patterns in each group. Specifically, we hypothesized that patients with TLE would show the most pronounced alterations in ipsilateral temporo-limbic regions, most notably in TLE-HS. We hypothesized that patients with GGE would show bilateral fronto-thalamocortical alterations. We also hypothesized that common white matter alterations would emerge across the patient groups and that many of these regional alterations would correlate with disease duration and be similar to those seen in other brain disorders.

## Materials and methods

All study participants provided written informed consent for the local study, and the local institutional review boards and ethics committees approved each included cohort study.

### Study sample

This study from the ENIGMA-Epilepsy working group consists of 21 cohorts from nine different countries and includes dMRI scans from 1,069 healthy controls and 1,249 adult epilepsy patients. Demographic and clinical characteristics of the samples are presented in Table 1 (by site) and Table 2 (across site). An epilepsy specialist assessed seizure and syndrome classifications at each center, using the International League Against Epilepsy (ILAE) criteria. For the TLE subgroups, we included anyone with the typical electroclinical constellation of this syndrome (Berg *et al*., 2010). All TLE-HS patients had a neuroradiologically-confirmed diagnosis of unilateral atrophy and increased T2 signal on clinical MRI, whereas all of the TLE-NL patients had a normal MRI undertaken at the same time as the analyzed dMRI scan. Participants with a normal MRI and frontal, occipital, or parietal epilepsy were labelled as ExE. Participants with tonic-clonic, absence or myoclonic seizures with generalized spike-wave discharges on EEG were included in the GGE group. We excluded participants with a progressive disease (e.g., Rasmussen’s encephalitis), malformations of cortical development, tumors or prior neurosurgery. Participants were between 18 and 70 years of age.

**Table 1.** Characteristics of the patient and control samples by site.

| Center | Age controls<br>(Mean ± SD) | Age cases<br>(Mean ± SD) | Age of onset<br>(Mean ± SD) | Duration of<br>illness<br>(Mean ± SD) | Total<br>controls | Total<br>cases | L TLE-HS<br>cases | R TLE-HS<br>cases | L TLE-NL<br>cases | R TLE-NL<br>cases | GGE<br>cases | ExE cases | Total *n* |
| --- | --- | --- | --- | --- | --- | --- | --- | --- | --- | --- | --- | --- | --- |
| Bonn | 38.0 ± 14.0 | 41.8 ± 12.9 | 17.7 ± 11.6 | 23.8 ± 14.2 | 44 | 64 | 42 | 22 | 0 | 0 | 0 | 0 | 108 |
| CUBRIC | 28.3 ± 8.4 | 28.6 ± 7.7 | 14.7 ± 4.0 | 14.0 ± 9.5 | 35 | 34 | 0 | 0 | 0 | 0 | 34 | 0 | 69 |
| EKUT | 40.0 ± 14.9 | 33.9 ± 13.1 | 19.3 ± 14.2 | 14.6 ± 10.4 | 19 | 15 | 0 | 0 | 0 | 0 | 15 | 0 | 34 |
| EPICZ | 38.6 ± 11.2 | 38.3 ± 9.7 | 19.1 ± 12.4 | 19.2 ± 12.5 | 113 | 86 | 19 | 26 | 26 | 15 | 0 | 0 | 199 |
| EPiGEN_Ireland | 34.8 ± 9.5 | 35.9 ± 9.0 | 19.1 ± 10.9 | 18.1 ± 12.9 | 67 | 47 | 9 | 5 | 17 | 16 | 0 | 0 | 114 |
| Florence | 34.6 ± 11.6 | 37.6 ± 13.5 | 14.0 ± 7.4 | 19.0 ± 9.9 | 35 | 4 | 0 | 0 | 0 | 2 | 0 | 2 | 39 |
| Genova | 25.3 ± 8.2 | 25.3 ± 8.2 | 10.4 ± 4.6 | 14.7 ± 7.7 | 20 | 17 | 0 | 1 | 3 | 4 | 7 | 2 | 37 |
| Greifswald | n/a | 34.6 ± 14.0 | 19.2 ± 9.7 | 15.4 ± 9.2 | n/a | 48 | 0 | 0 | 0 | 0 | 35 | 13 | 48 |
| HFHS_2.6mm | 29.5 ± 7.0 | 38.3 ± 13.0 | 19.0 ± 16.9 | 19.0 ± 12.9 | 21 | 57 | 6 | 8 | 8 | 7 | 0 | 28 | 78 |
| HFHS_3mm | n/a | 40.1 ± 15.4 | - | - | 0 | 14 | 3 | 3 | 3 | 3 | 0 | 2 | 14 |
| IDIBAPS_31Dir | n/a | 39.3 ± 10.1 | 22.5 ± 13.4 | 16.4 ± 11.1 | 0 | 25 | 5 | 7 | 3 | 2 | 1 | 7 | 25 |
| IDIBAPS_39Dir | 32.4 ± 6.6 | 33.3 ± 9.4 | 16.9 ± 11.8 | 17.6 ± 11.9 | 30 | 19 | 3 | 8 | 0 | 1 | 1 | 6 | 49 |
| IDIBAPS_88Dir | 34.1 ± 5.1 | 37.6 ± 10.5 | 15.6 ± 11.0 | 22.0 ± 13.3 | 22 | 27 | 5 | 10 | 3 | 3 | 0 | 6 | 49 |
| KCL | 30.4 ± 7.5 | 35.0 ± 9.5 | - | - | 92 | 89 | 24 | 38 | 0 | 0 | 27 | 0 | 181 |
| Liverpool_Walton | 31.8 ± 7.7 | 31.1 ± 10.3 | 14.3 ± 8.8 | 16.7 ± 11.9 | 40 | 33 | 5 | 3 | 8 | 5 | 1 | 11 | 73 |
| Melbourne | n/a | 37.2 ± 9.9 | 22.4 ± 12.8 | 14.8 ± 11.2 | 0 | 25 | 13 | 5 | 6 | 1 | 0 | 0 | 25 |
| MNI | 30.7 ± 7.4 | 32.1 ± 9.5 | 16.4 ± 9.6 | 15.4 ± 10.6 | 46 | 149 | 20 | 22 | 23 | 15 | 0 | 69 | 195 |
| MUSC | 55.2 ± 6.6 | 36.6 ± 10.7 | 18.5 ± 13.5 | 17.8 ± 12.1 | 58 | 43 | 19 | 6 | 11 | 7 | 0 | 0 | 101 |
| NYU | 30.3 ± 10.0 | 32.6 ± 8.9 | 24.0 ± 12.7 | 8.6 ± 9.9 | 26 | 45 | 2 | 2 | 8 | 3 | 0 | 30 | 71 |
| UCL | 37.7 ± 12.4 | 38.7 ± 11.4 | 13.9 ± 10.7 | 24.7 ± 14.3 | 29 | 53 | 24 | 13 | 6 | 10 | 0 | 0 | 82 |
| UCSD | 37.3 ± 13.5 | 34.0 ± 11.9 | 19.6 ± 12.9 | 15.2 ± 13.3 | 47 | 55 | 17 | 12 | 13 | 13 | 0 | 0 | 102 |
| UMG | 36.0 ± 10.4 | 34.1 ± 10.8 | 16.6 ± 8.1 | 15.6 ± 11.7 | 20 | 36 | 2 | 5 | 0 | 0 | 24 | 5 | 56 |
| UNAM | 33.7 ± 12.2 | 31.4 ± 11.7 | 15.8 ± 11.1 | 15.5 ± 12.8 | 34 | 30 | 9 | 10 | 9 | 2 | 0 | 0 | 64 |
| UNICAMP | 35.5 ± 10.8 | 40.6 ± 9.9 | 12.3 ± 9.6 | 28.3 ± 12.2 | 271 | 234 | 92 | 74 | 15 | 4 | 37 | 12 | 505 |
| Combined | 35.5 ± 11.7 | 36.1 ± 11.2 | 16.3 ± 11.3 | 19.9 ± 13.0 | 1069 | 1249 | 319 | 280 | 162 | 113 | 182 | 193 | 2318 |

**Table 2.** Demographic and clinical characteristics of the total sample. Post-hoc comparisons revealed that controls were younger than both TLE-HS groups and older than patients with GGE and ExE (all p-values<0.05). Both TLE-HS groups and the left TLE-NL group were older than the GGE and ExE groups (*p*<0.05). The right TLE-HS group was older than both TLE-NL groups (*p*<0.05).

|  | **N** | **Age (SD)** | **Sex**<br>**(% Male)** | **Age of onset**<br>**(SD)** | **Duration of**<br>**illness (SD)** |
| --- | --- | --- | --- | --- | --- |
| **Controls** | 1,069 | 35.5 (11.7) | 41.7 | ----- | ----- |
| **All patients** | 1,249 | 36.1 (11.2) | 42.9 | 16.3 (11.3) | 19.9 (13.0) |
| **L TLE-HS** | 319 | 38.3 (10.6) | 43.3 | 14.5 (11.5) | 24.2 (13.3) |
| **R TLE-HS** | 280 | 39.3 (10.8) | 42.5 | 15.3 (11.6) | 24.1 (13.9) |
| **L TLE-NL** | 162 | 36.1 (10.9) | 38.3 | 18.8 (11.6) | 17.1 (12.1) |
| **R TLE-NL** | 113 | 35.1 (11.4) | 37.2 | 19.5 (12.2) | 15.1 (11.2) |
| **GGE** | 182 | 31.8 (9.9) | 44.5 | 14.8 (7.3) | 16.8 (10.5) |
| **ExE** | 193 | 32.7 (11.5) | 43.5 | 17.6 (11.1) | 15.1 (11.5) |

### Image processing and analysis

Scanner descriptions and acquisition protocols for all sites are provided in Supplementary Table 1. Individual scanners that used different acquisition protocols are listed as separate scanner instances. Each site conducted the preprocessing of diffusion-weighted images, including eddy current correction, echo-planar imaging (EPI)-induced distortion correction, and tensor estimation. Next, diffusion-tensor imaging (DTI) images were processed using the ENIGMA-DTI protocols. These image processing and quality control protocols are freely available at the ENIGMA-DTI (http://enigma.ini.usc.edu/ongoing/dti-working-group/) and NITRC (https://www.nitrc.org/projects/enigma_dti/) webpages. Measures of FA, MD, AD and RD were obtained for 38 regions of interest using the Johns Hopkins University (JHU) atlas (Figure 1).

**Figure 1.**
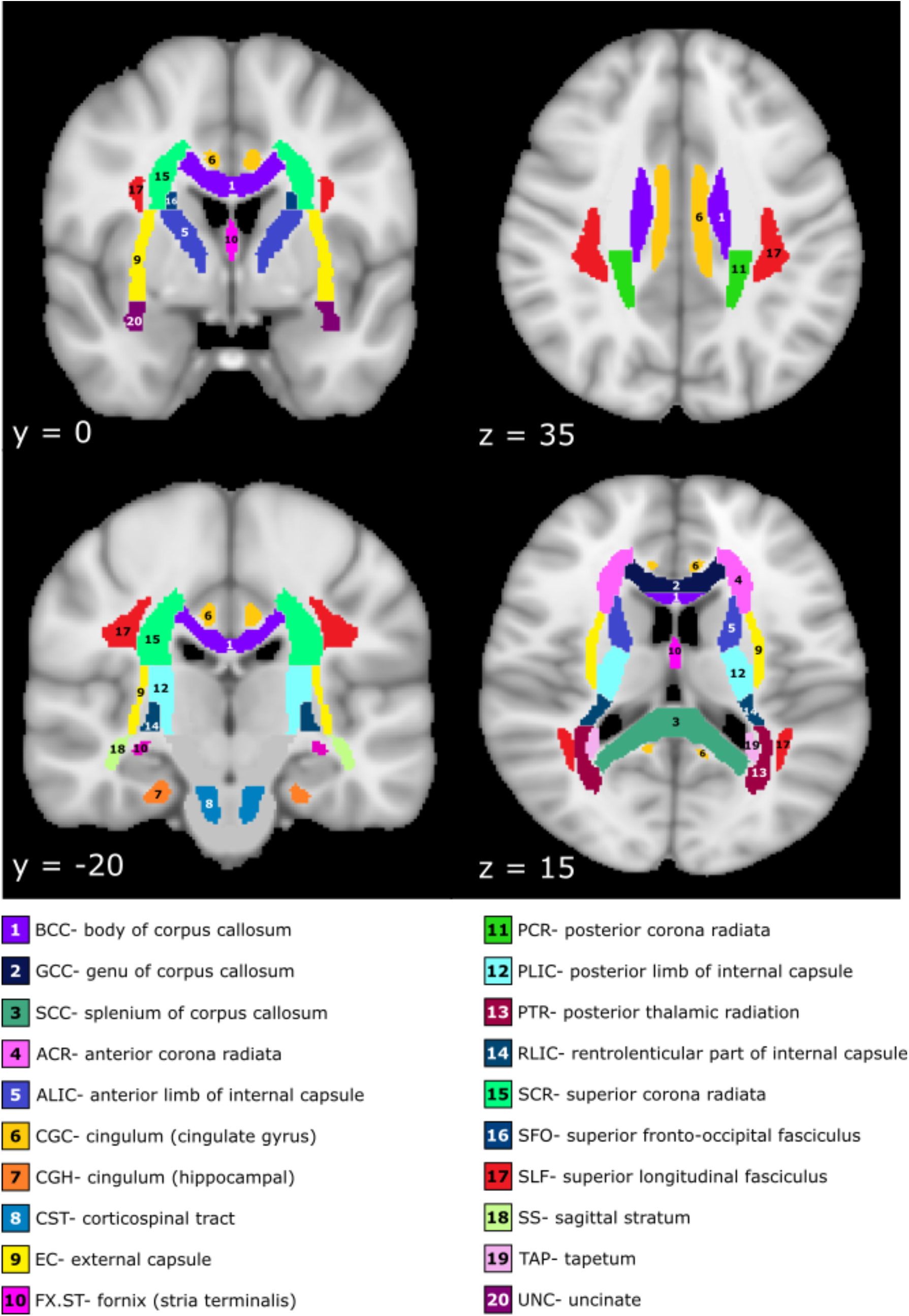
Fiber atlas.

For analyses of all patients and for each syndrome, we used the left and right tracts, the midline structures of the body (BCC), genu (GCC), and splenium (SCC) of the corpus callosum and the average diffusion metric (FA/MD/AD/RD) across the whole brain (38 ROIs). Corrections for multiple comparisons were carried out for each epilepsy syndrome, based on the number of ROIs: 34 bilateral white matter regions + BCC + GCC + SCC + Average FA=38 ROIs: Bonferroni-corrected threshold for significance p=0.05/38=0.001.

### Data harmonization

The batch-effect correction tool, ComBat, was used to harmonize between-site and between-protocol variations in diffusion metrics as previously demonstrated (Fortin *et al*., 2017). The method globally rescales the data (all ROIs for FA, MD, RD or AD separately) for each scanner instance using a z-score transformation map common to all features. ComBat uses an empirical Bayes framework (Johnson *et al*., 2007) to improve the variance of the parameter estimates, assuming that all ROIs share the same common distribution. Thus, all ROIs are used to inform the statistical properties of the scanner effects. We set each scanner instance as each individual scanner used in the collection of MR exams, and where there were different scanning protocols used on the same scanner, each protocol was set as a different scanning instance. Scanner type was used as the batch effect and diagnosis (patients versus controls) and syndrome (GGE, TLE-HS, TLE-NL, and ExE) were used as the biological phenotypes of interest. This technique has been recently applied in other ENIGMA DTI investigations of brain disorders (Villalón-Reina *et al*., 2019; Zavaliangos-Petropulu *et al*., 2019).

### Statistical analysis

Statistical analysis was performed in the Statistical Package for the Social Sciences (SPSS v26.0). Analysis of variance (ANOVA) was used to test for differences in demographic and clinical characteristics among the epilepsy syndromes. To test for differences between syndromes and controls and to test for global effects of age at scan and sex on white matter, multivariate analysis of covariance (MANCOVA) was performed per diffusion metric, adjusting for age, age^2^ and sex. Age^2^ was included in all analyses to model the non-linear effects of age on diffusivity measures (Lebel *et al*., 2012). ANCOVAs were then performed of each patient syndrome compared to controls controlling for age, age^2^ and sex and Bonferroni corrected at *p*<0.001. Cohen’s *d* effect sizes (ES) were calculated for each right and left fiber tract between controls and each patient syndrome based on the estimated marginal means (adjusted for age, age^2^, and sex) and interpreted according to the following criteria: *small d=0.20-0.49; medium d=0.50-0.79; large d>=0.80* (Sawilowsky, 2009). Throughout the text and figures, positive ES values correspond to patients having higher values than controls, whereas negative ES values correspond to patients having lower values relative to controls. Partial correlations controlling for the same covariates were performed to evaluate the relationship between each fiber tract FA/MD and age of seizure onset and disease duration (corrected *p*<0.001). To demonstrate the most robust group differences, only medium and large effects are discussed in the text (see Supplementary materials for all results: Tables 5-32 and Figures 5–6).

## Results

### Demographics

Demographic and clinical characteristics of each sample are presented in Table 2. ANOVA revealed differences across the cohort in age [*F*(6, 2083) = 13.3, *p*<0.001)] and across the patient syndromes in age of seizure onset [*F*(5, 1095) = 6.3, *p*<0.001] and disease duration [*F*(5,1039) = 22.4, *p*<0.001]. Post-hoc comparisons revealed that controls were younger than both TLE-HS groups and older than patients with GGE and ExE (all p-values<0.05). Both TLE-HS groups and the left TLE-NL group were older than the GGE and ExE groups (*p*<0.05). The right TLE-HS group was older than both TLE-NL groups (*p*<0.05).

The GGE and TLE-HS groups had an earlier age of seizure onset than the TLE-NL groups. Duration of illness in TLE-HS groups was longer than in all other groups.

### Data harmonization with ComBat

Initial frequency plots revealed high variability in the distribution of diffusivity measures (e.g., mean FA, mean MD) among scanner instances (Fig. 2A). After batch correction with ComBat, the distributions were centered around a common mean (Fig. 2B), but maintained their expected association with age (Fig. 2C). Following this process, extreme ROI outliers beyond 3 SD were removed from the subsequent analysis (i.e., per ROI for a given subject, not per subject).

**Figure 2.**
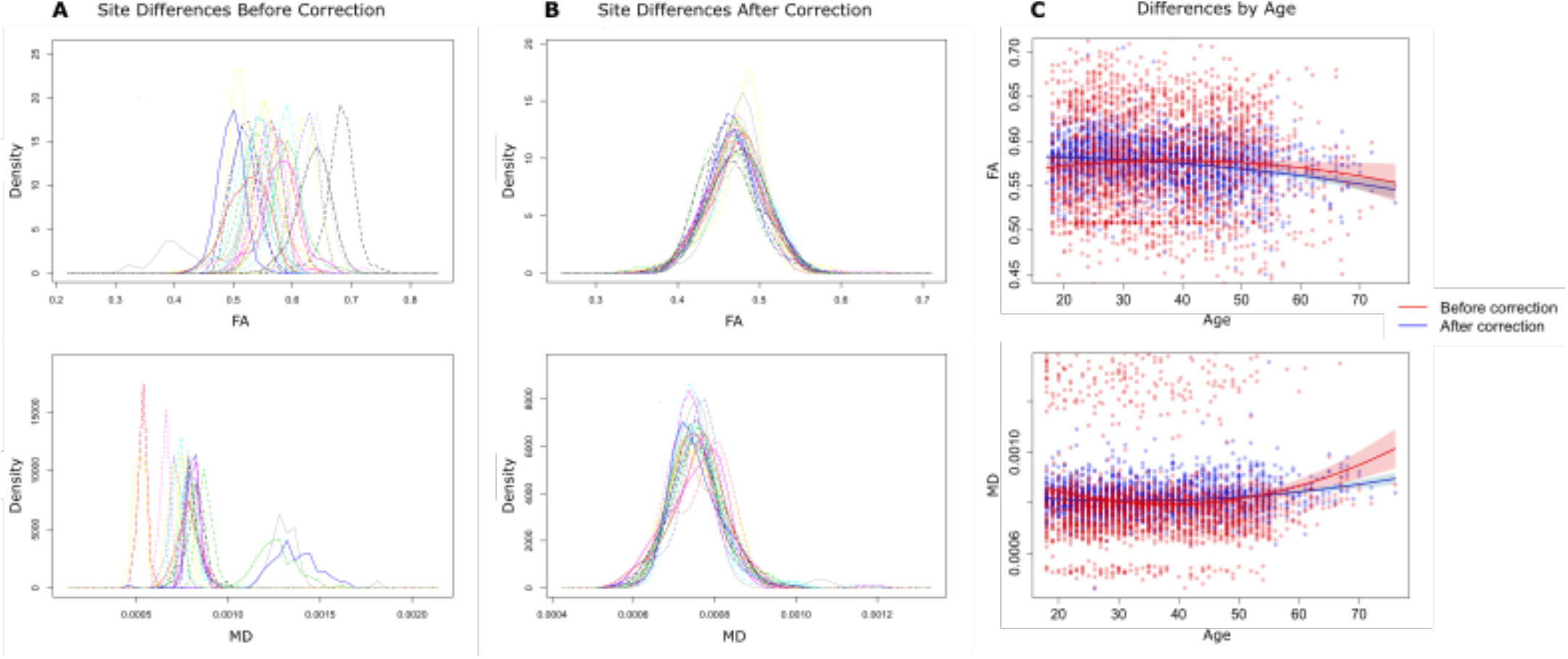
dMRI harmonization using ComBat. Average FA (top) and MD (bottom) measures across 24 scanners showing differences in mean FA measures per scanner (left) which are harmonized using ComBat (middle). The process corrects the variance in scanner without altering the biological variance expected with age (right). Red= *before* correction; Blue= *after* correction.

### All epilepsies group

#### Multivariate tests of within-subject effects

Comparing the whole epilepsy group with healthy controls, significant differences were observed in FA (*F*(228,13452)=4.7, *p*<0.001, Pillai’s Trace=0.44, partial η^2^=0.07), MD (*F*(228,11688)=2.8, *p*<0.001, Pillai’s Trace=0.31, partial η^2^=0.05), RD (*F*(228,11790)=3.28, *p*<0.001, Pillai’s Trace=0.36, partial η^2^=0.06), and AD (*F*(228,11946)=1.96, *p*<0.001, Pillai’s Trace=0.22, partial η^2^=0.04). Sex, age, and age^2^ all significantly contributed to the model (see Supplementary Table 4). Compared to females, males generally had higher FA, slightly higher RD and no difference in MD or AD.

#### All epilepsies vs healthy controls

The “all epilepsies” group showed lower FA than controls globally in 36 of 38 ROIs (*p*<0.001; Fig. 3, Supplementary Table 5), with medium ES observed for the average FA (*d*=−0.71), followed by external capsule (EC; left *d*=−0.64, right *d*=−0.63), body (*d*=−0.59) and genu (*d*=−0.59) of the corpus callosum (BCC and GCC), cingulate gyrus of the cingulum bundle (CGC, left *d*=−0.57, right *d*=−0.50), sagittal stratum (SS, left *d*=0.55, right *d*=0.52), anterior *corona radiata* (ACR, left *d*=−0.50, right *d*=−0.52), and left parahippocampal cingulum (CGH, *d*=−0.52).

**Figure 3.**
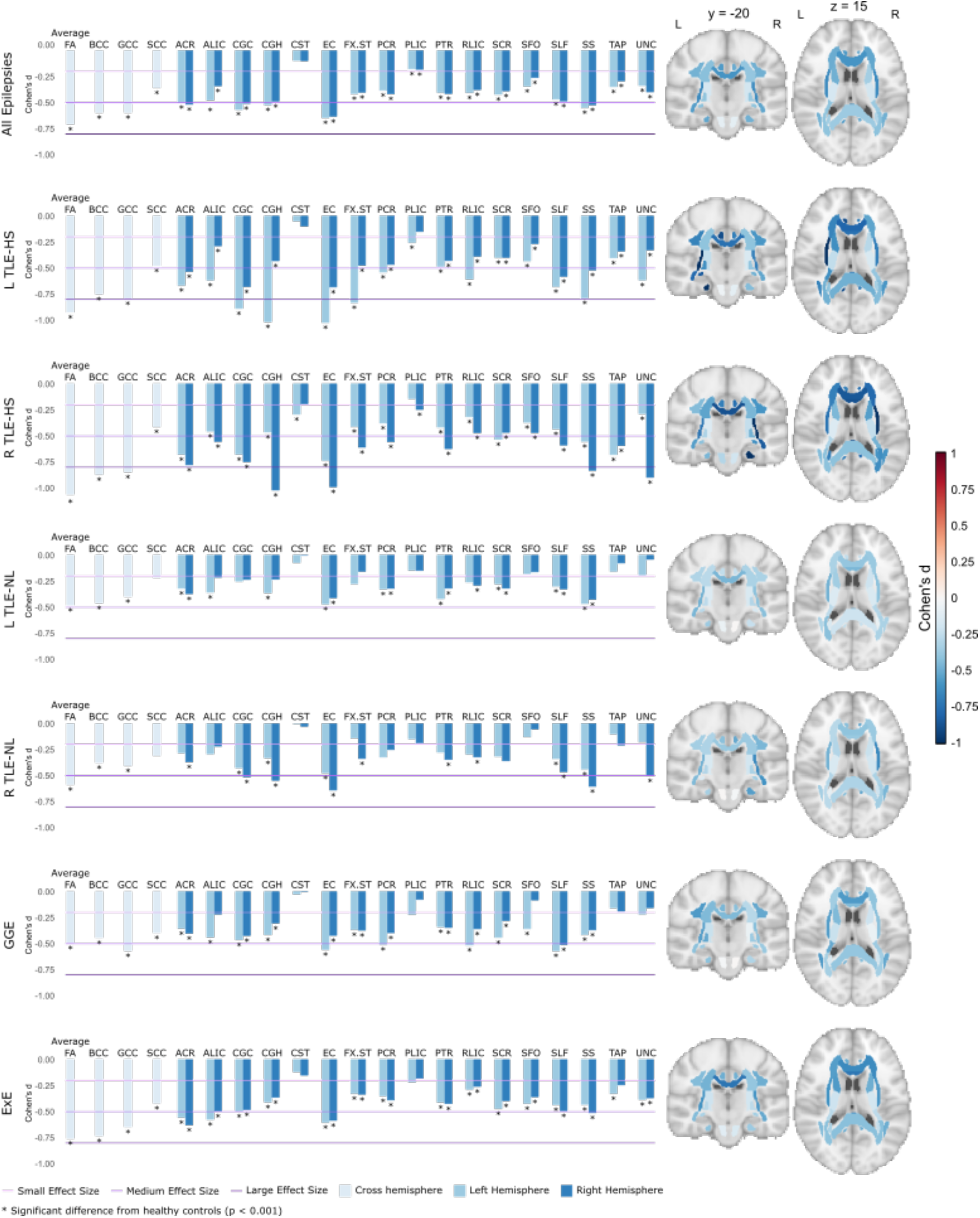
FA Effect Size Bar Graphs.

The “all epilepsies” group showed higher MD than the controls with small effect size in 27 ROIs (Fig. 4, Supplementary Table 12). Similar to the MD, patients showed higher RD with small effect size in 34 ROIs (Supplementary Fig. 1, Supplementary Table 19) with a medium-sized effect seen in the right EC (*d*=0.52). Small-sized effects were observed for AD in 8 ROIs (Supplementary Fig. 2, Supplementary Table 26).

**Figure 4.**
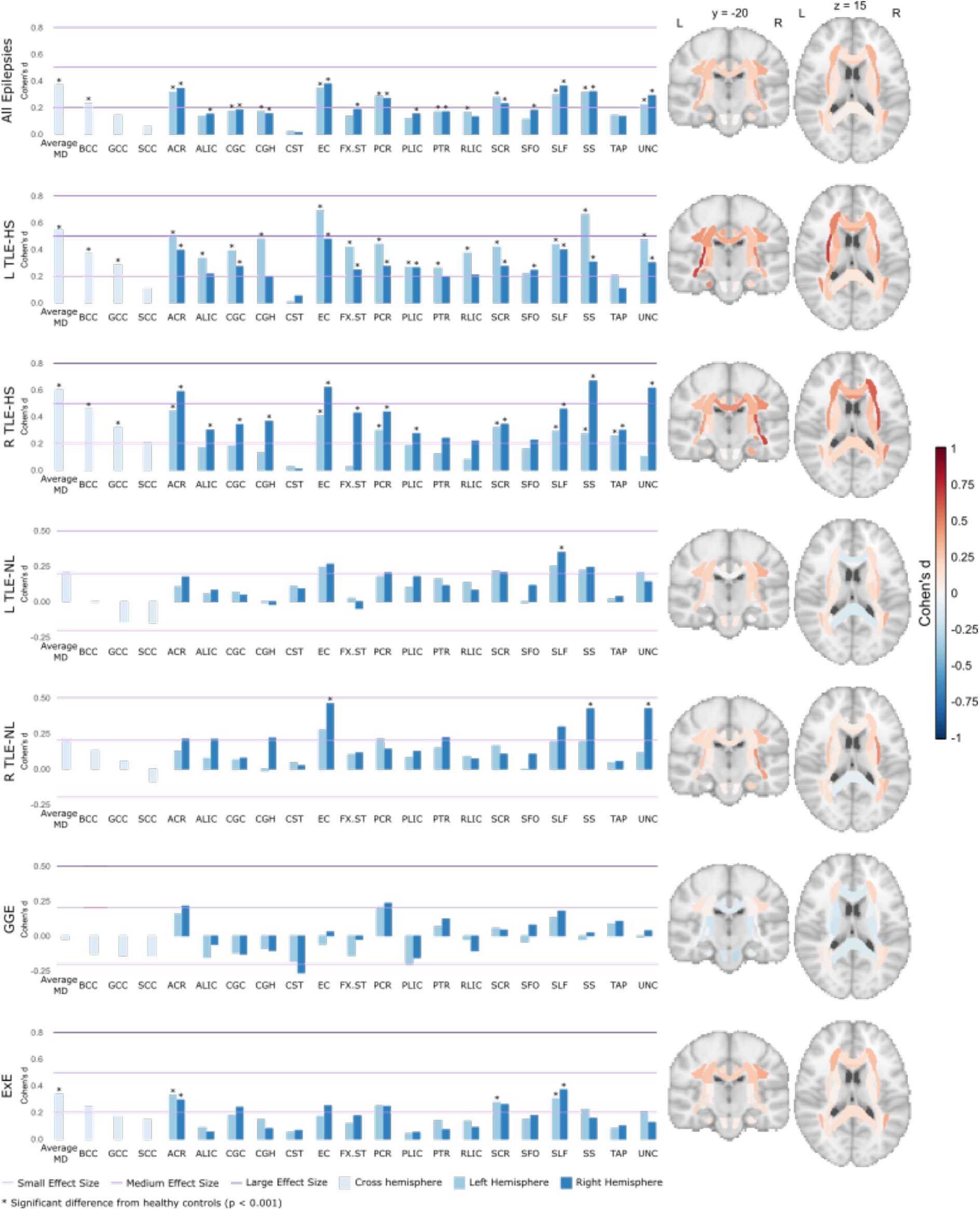
MD Effect Size Bar Graphs.

#### Age of onset and disease duration

Across all epilepsy patients, earlier age of onset was associated with lower FA across 28 ROIs (*r*=0.1 to 0.3, *p*<0.001), higher MD in 16 ROIs, and higher RD in 28 ROIs (Supplementary Tables 33,35,37). The most robust associations were observed between an earlier age of seizure onset and lower FA in the Average FA, GCC, and bilateral EC, and CGC (r’s between 0.16 and 0.19). There were no significant relationships between age of onset and AD in the “all epilepsies” group (Supplementary Table 39).

Across all epilepsy patients, disease duration showed significant but small effects on diffusivity measures (Supplementary Tables 34, 36, 38, 40). A longer disease duration was associated with lower FA in 14 ROIs, increased MD in 9 ROIs, and increased RD in 27 ROIs. Of note, longer disease duration was associated with lower FA in BCC, CGC, and EC, and with lower Average FA (r’s = −0.15 to −0.17). There were no significant relationships between disease duration and AD in all patients.

### TLE-HS group

#### TLE-HS patients vs healthy controls

Left TLE-HS patients (*n*=319) showed significantly lower FA than controls in 35 of 38 ROIs (Fig. 3, Supplementary Table 6), with large ES differences in the left EC (*d*=−1.02), left CGH (*d*=−1.01), Average FA (*d*=−0.92), left CGC (*d*=−0.89), and left fornix/*stria terminalis* (FXST, *d*=−0.83). Medium-sized effects were observed in GCC (*d*=−0.79), left SS (*d*=−0.78), BCC (*d*=−0.75), right CGC (*d*=−0.68), right EC (*d*=−0.68), left superior longitudinal fasciculus (SLF, *d*=−0.67), left ACR (*d*=−0.66), left anterior limb of the internal capsule (ALIC, *d*=−0.61), left retrolenticular portion of the internal capsule (RLIC, *d*=−0.61), left uncinate (UNC, *d*=−0.61), right SLF (*d*=−0.58), left PCR (*d*=−0.54), right ACR (*d*=−0.53), and right SS (*d*=−0.52).

Significantly higher MD was observed in 28 ROIs (Fig. 4, Supplementary Table 13), with medium-sized effects in the left EC (*d*=0.69), left SS (*d*=0.66), and average MD (*d*=0.55). Left TLE-HS patients showed significantly higher RD in 33 of 38 ROIs (Supplementary Fig. 1, Supplementary Table 20). A large effect of higher RD was observed for the left EC (*d*=0.83), and medium-sized effects were seen for the left SS (*d*=0.79), left CGC (*d*=0.69), left CGH (*d*=0.67), average RD (*d*=0.66), right EC (*d*=0.61), left FXST (*d*=0.59), left ACR (*d*=0.57), SLF (left *d*=0.56, right *d*=0.53), right ACR (*d*=0.52), and left RLIC (*d*=0.51). Small-sized effects were observed for AD in 5 ROIs (Supplementary Figure 2, Supplementary Table 27).

Right TLE-HS patients (*n*=280) showed lower FA than controls in 26 of 38 ROIs (*p*<0.001; Fig. 3, Supplementary Table 7), with large effects observed for the average FA (*d*=−1.06), right CGH (*d*=−1.02), right EC (*d*=−0.99), right UNC (*d*=−0.90), BCC (*d*=−0.87), GCC (*d*=−0.85) and right SS (*d*=−0.83). Medium-sized effects were observed in the right ACR (*d*=−0.78), right CGC (*d*=−0.73), left EC (*d*=−0.68), left ACR (*d*=−0.68), left CGC (*d*=−0.68), left tapetum (TAP, *d*=−0.68), right PTR (*d*=−0.62), right FX/ST (*d*=−0.61), right SLF (*d*=−0.59), right TAP (*d*=−0.59), right ALIC (*d*=−0.55), right PCR (*d*=−0.55), left SS (*d*=−0.55), and left SCR (*d*=−0.53).

Higher MD in the patient group were observed in 23 ROIs (Fig. 4, Supplementary Table 14), with medium-sized effects shown in the right SS (*d*=0.67), right EC (*d*=0.62), right UNC (*d*=0.62), average MD (*d*=0.60) and right ACR (*d*=0.59). Significantly higher RD in the patient group were observed in 31 ROIs (Supplementary Figure 1, Supplementary Table 21), with a large effect observed for the right EC (*d*=0.84), and medium effects seen for the right SS (*d*=0.76), right UNC (*d*=0.72), average RD (*d*=0.70), right ACR (*d*=0.69), left EC (*d*=0.63), right SLF (*d*=0.62), right CGH (*d*=0.61), right CGC (*d*=0.57), BCC (*d*=0.56), right FX/ST (*d*=0.52), and left ACR (*d*=0.51). Small effects were observed for AD in 6 ROIs (Supplementary Figure 2, Supplementary Table 28).

#### Age of onset and disease duration

For left TLE-HS, earlier age of onset was associated with lowerFA in 4 ROIs, including the Average FA, BCC, GCC, and left EC. Earlier age of seizure onset was associated with higher RD in 2 ROIs (Supplementary Tables 33,37). There was no detected relationship between age of onset and either MD or AD in the left TLE-HS group (Supplementary Tables 35, 39).

For right TLE-HS, earlier age of onset was related to lower FA across 9 ROIs, including the Average FA, BCC, SCC, right EC, left and right CGH, right PCR, and right SLF, increased MD in 7 ROIs, and increased RD in 7 ROIs (Supplementary Tables 33, 35, 37). There was no detected relationship between age of onset and AD in the TLE-HS groups (Supplementary Table 39).

For left TLE-HS patients, a longer disease duration was associated with lower FA in 4 ROIs (BCC, bilateral CGC, and left EC) and higher RD in one ROI (left SS). There were no significant relationships between disease duration and MD or AD (Supplementary Tables 34, 36, 38, 40). For right TLE-HS, disease duration showed significant small effects on diffusivity measures (Supplementary Tables 34, 36, 38, 40). A longer disease duration was associated with lower FA in 8 ROIs (Average FA and SCC, bilateral TAP, and right CGH, EC, PCR, and UNC), higher MD in 6 ROIs, and higher RD in 10 ROIs.

### TLE-NL group

#### TLE-NL vs healthy controls

The left TLE-NL patients (*n*=162) showed significantly lower FA than controls in 20 ROIs (*p*<0.001; Fig. 3, Supplementary Table 8) of small ES. MD was higher in one ROI, the right SLF (*d*=0.35, *p*<0.001, Fig. 4, Supplementary Table 15). Significantly higher RD was observed in 6 ROIs (Supplementary Figure 1, Supplementary Table 22) of small ES. No significant effects of AD were detected (Supplementary Figure 2, Supplementary Table 29).

Right TLE-NL patients (*n* = 113) showed significantly lower FA than controls in 19 ROIs (*p*<0.001; Fig. 3, Supplementary Table 9), with medium-sized effects observed in the right EC (*d*=−0.64), right SS (*d*=−0.60), Average FA (*d*=−0.58), right CGH (*d*=−0.55), right CGC (*d*=−0.51) and right UNC (*d*=−0.50). MD was increased in three ROIs (*p*<0.001, Fig. 4, Supplementary Table 16), specifically the right EC (*d*=0.46), right UNC (*d*=0.42) and right SS (*d*=0.42). Significantly higher RD was observed in 4 ROIs (Supplementary Figure 1, Supplementary Table 23), with a medium-sized effect shown in the right EC (*d*=0.50). No significant effects of AD were detected (Supplementary Figure 2, Supplementary Table 30).

#### Age of onset and disease of illness

For left TLE-NL, no diffusivity measure was associated with earlier age of onset or duration of illness (Supplementary Tables 33-40).

For right TLE-NL patients, younger age of onset and was related to lower FA in the right UNC (*r*=0.30, *p*<0.001), but not MD, AD, or RD (Supplementary Tables 33, 35, 37, 39). Disease duration was also associated with decreased FA in the right UNC (*r*= =−0.37, *p*<0.001), but not MD, AD, or RD (Supplementary Tables 34, 36, 38, 40).

### GGE group

#### GGE patients vs healthy controls

GGE patients (*n*=113) showed significantly lower FA than controls in 28 ROIs (*p*<0.001; Fig. 3, Supplementary Table 10), with medium-sized effects observed in the GCC (*d*=−0.57), left SLF (*d*=−0.57), left EC (*d*=−0.55), left RLIC (*d*=−0.51), right SLF (*d*=−0.51) and left PCR (*d*=−0.50). No significant effects were seen for MD or AD (Fig. 4, Supplementary Fig. 2, Supplementary Tables 17, 31), but small ESs were seen for RD (4 ROIs, *p*<0.001, Supplementary Fig. 1, Supplementary Table 24).

#### Age of onset and disease duration

For GGE patients, there was no significant association between diffusivity measures and either age of onset of epilepsy, or its duration.

### ExE group

#### ExE patients vs healthy controls

ExE patients (*n*=193) showed significantly lower FA than controls in 33 ROIs (*p*<0.001; Fig. 1, Supplementary Table 11), with medium-sized effects observed for Average FA (*d*=−0.75), BCC (*d*=−0.65), GCC (*d*=−0.64), right ACR (*d*=−0.63), bilateral EC (left *d*=−0.60, right *d*=−0.58), left ALIC (*d*=−0.57), left ACR (*d*=−0.55), and right SS (*d*=−0.51). Small-sized effects were seen in MD (6 ROIs, *p*<0.001, Fig. 4, Supplementary Table 18) and RD (22 ROIs, *p*<0.001, Supplementary Figure 1, Supplementary Table 25). No significant effects were seen in AD (Supplementary Figure 2, Supplementary Table 32).

#### Age of onset and disease duration

For ExE patients, there were no significant associations between diffusivity measures and either age of onset of epilepsy, or its duration.

### Cross-syndrome comparisons

Post-hoc comparisons were conducted across the syndromes in the five ROIs that showed the largest ES in the “all epilepsies” analysis, namely Average FA/MD, ACR, BCC, CGC, and EC averaged across hemispheres (Figure 5). ANCOVAs, adjusting for age, age^2^, sex, age of seizure onset, and disease duration revealed significant group differences in Average FA [*F* (5, 874) = 3.8, *p*<0.05], as well as FA of the CGC [*F*(5, 874) = 5.0, *p*<0.05], and EC [*F*(5, 874) = 5.8, *p*<0.05].

**Figure 5.**
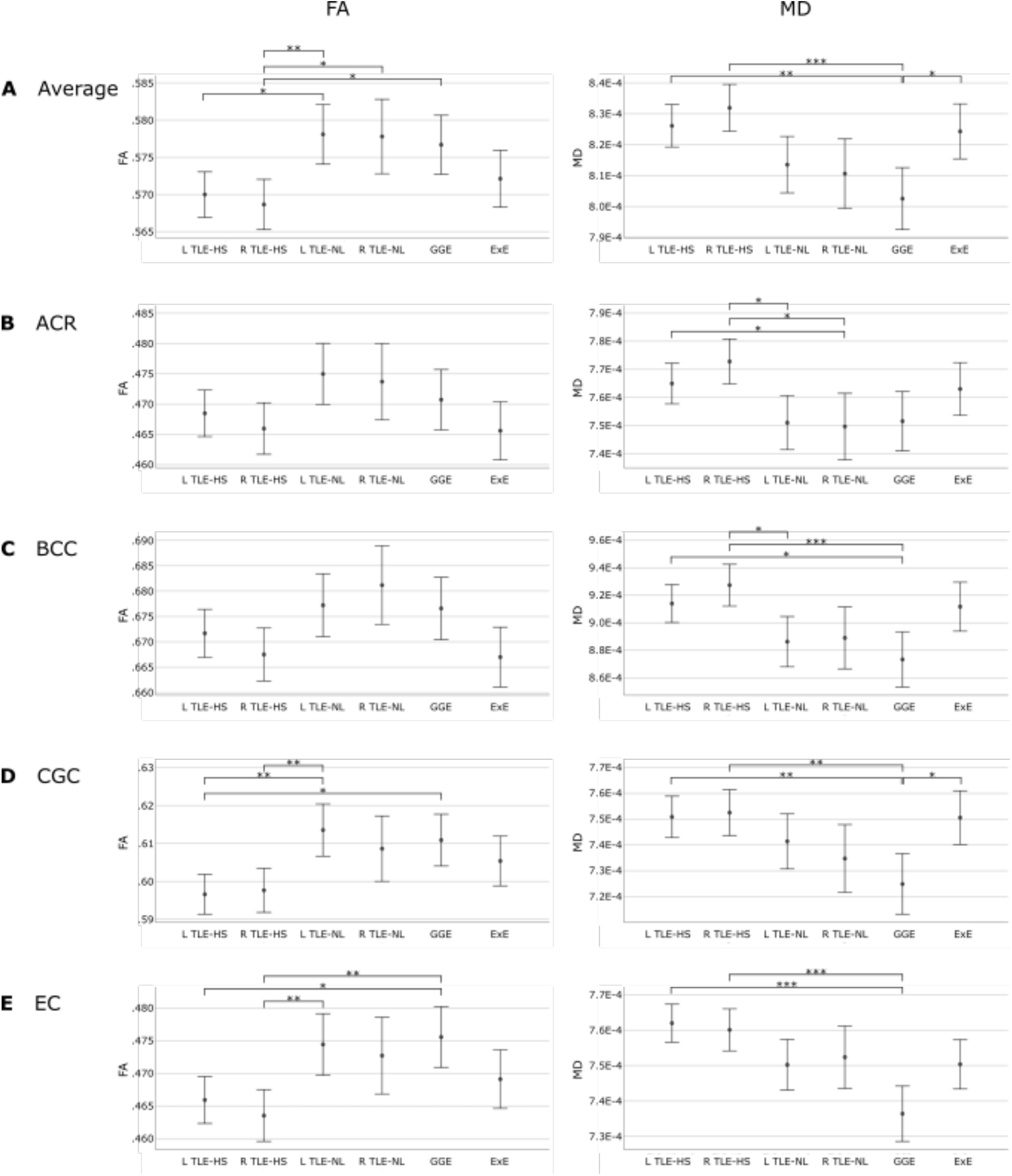
Syndromic difference in average FA and MD in five ROIs. Mean FA (left) and MD (right) for each patient syndrome, controlling for age, age^2^, sex, age of onset, and disease duration. Error bars reflect 95% confidence intervals. Significant differences are marked with asterisks (* for p<0.05, ** for p<0.01, *** for p<0.001).

There were differences across syndromes or the Average MD [*F*(5, 874) = 5.3, *p*<.05], ACR [*F*(5, 874) = 3.8, *p*<0.05], BCC [*F*(5, 874) = 4.9, *p*<0.05], CGC [*F*(5, 874) = 4.2, *p*<0.05], and the EC [*F*(5, 874) = 6.2, *p*<0.05]. Patients with left and right TLE-HS generally showed lower FA and higher MD than those with TLE and normal MRI, GGE in Average FA/MD, CGC, and the EC. The nonlesional groups (GGE, TLE-NL, and ExE) did not differ from one another.

### Comparisons with other disorders

Given the many common white matter FA differences observed across epilepsy syndromes, the question arises as to whether these effects are specific to epilepsy or also seen with other brain disorders. Figure 6 displays ES differences observed in the “all epilepsies” group (n=1,249) relative to findings from four other ENIGMA working groups: schizophrenia (SCZ; n=1,984; mean age, 36.2; 67% men), 22q11 syndrome (n=334; mean age, 16.9; 54% men), bipolar disorder (BP; n=1,482; mean age, 39.6; 39.3% men), and major depressive disorder (MDD; n=921; mean age, 40.7; 39% men). The magnitude of the ESs were typically larger in epilepsy, compared to the other disorders for most white matter regions. Across the white matter regions, ESs in patients with epilepsy were significantly correlated with those in patients with BP (Spearman’s rho *r*=0.53, *p*<0.05), SCZ (*r*=0.53, *p*<0.05), and MDD (*r*=0.44, *p*<0.05).

**Figure 6.**
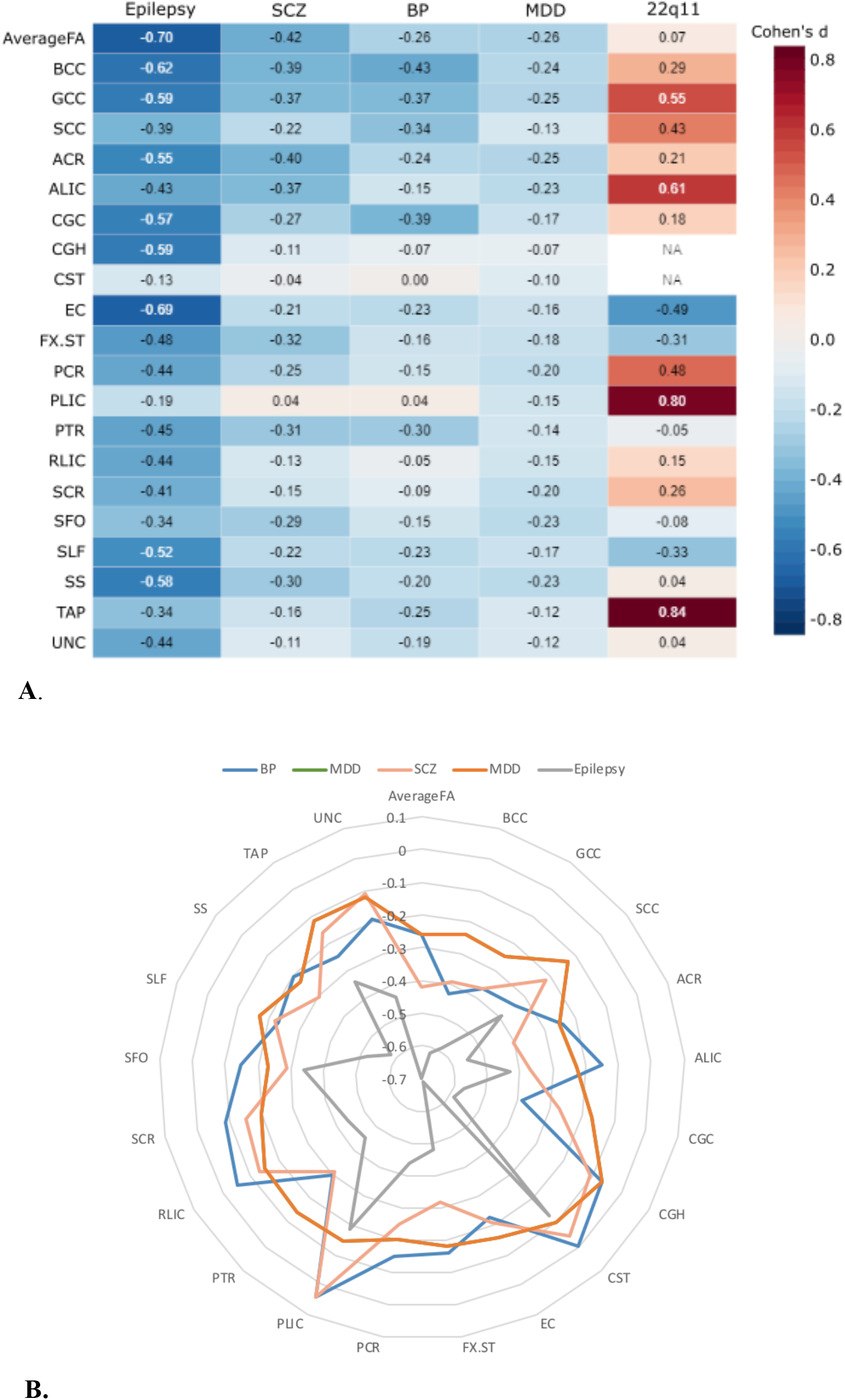
**A). Heat map of FA effect sizes for the “all epilepsies” group compared to those in four other ENIGMA disorders**: SCZ = schizophrenia; BP = bipolar disorder; MDD = major depressive disorder. **B). Radar plot of the four disorders that showed significant correlations across white matter tracts.** Positive values reflect patient group values were on average higher than controls, whereas negative values reflect cases where patient group values were on average lower than that of controls.

### Approach for multiple comparisons correction

Regional differences in diffusion parameters between syndromes and healthy controls were expressed as ESs (Cohen’s d). In order to identify significant differences we adopted a Bonferroni correction that adjusted for testing 38 ROIs for each syndrome (*p*<0.001). However, an even more conservative approach would correct for each of the four diffusion metrics across the seven epilepsy syndromes. Implementing this very conservative Bonferroni cutoff would result in a threshold of *p*<0.05/(38*7*4)=4.7e-05. The combination of the large sample size and the observed medium to large effect sizes in the study results in most regional differences remaining significant even at the very conservative p-value threshold. However, in contrasts involving a smaller number of patients (controls compared to, e.g., GGE or TLE-NL) more regions would lose the “significant” label due to the reduced statistical power. Despite these few exceptions, the exact *p*-value cutoff does not alter the main finding that there were widespread white matter abnormalities across epilepsy syndromes. We report all effect sizes, confidence intervals, and *p*-values in supplementary material for researchers interested to examine these results with alternate definitions of statistical significance.

## Discussion

This multi-site DTI study in epilepsy compared data from 1,249 patients with common epilepsy syndromes to 1,069 healthy controls. Data were acquired at 21 sites across North America, South American, Europe, and Australia and harmonized using the same postprocessing pipeline and batch effect harmonization tool.

There were marked white matter alterations across epilepsy syndromes compared to controls, with the most pronounced effects in patients with TLE-HS and modest effects in GGE. The anterior-midline fibers, including the CC, CGC, ACR and EC, were among the most affected fiber bundles *across* epilepsy syndromes--a finding not previously recognized in the literature, and one that could suggest vulnerability of these fibers to the consequences of epilepsy (e.g., treatments, recurrent seizures).

Previous studies have investigated white matter alterations *within* a specific epilepsy syndrome, and our first analysis included a diverse aggregation of epilepsy syndromes to address the question of shared pathological changes *across* syndromes. This analysis revealed widespread reductions in FA across most association, commissural, and projection fibers bilaterally, with smaller effects of increased MD. The most robust alterations were observed in frontocentral regions, including the *genu* and body of the CC, ACR, CGC, and EC. These regional changes mirror results from our structural MRI findings (Whelan *et al*., 2018), which revealed subcortical atrophy and neocortical thinning in fronto-central, midline structures, including the thalamus, pallidum, pre- and post central gyri, and superior frontal regions bilaterally. These regions showed the strongest association with both age of seizure onset and disease duration. Therefore, white matter abnormalities in these regions may be a result of both shared developmental (i.e., disruptions of late-myelinating pathways due to seizures) and degenerative (i.e., demyelination and/or axonal loss due to years of epilepsy, recurrent seizures, exposure to anti-epileptic drugs (AEDs), and other factors) processes. Although lower FA and higher MD have been interpreted as reflecting a combination of pathological processes, these differences appear to be primarily driven by higher RD, supporting the concept of disrupted or altered myelin rather than significant axonal loss (Arfanakis *et al*., 2002; Concha *et al*., 2009)

### Temporal Lobe Epilepsy

A recent imaging-based meta-analysis (Slinger *et al*., 2016) found that patients with TLE had pronounced and widespread white matter injury relative to other patient syndromes. We found that this pattern was much more robust and ipsilateral in patients with HS, particularly in hippocampal afferent and efferent tracts, including the CGH, FX/ST, and UNC. The proximity of these latter white matter regions to the epileptogenic zone and being greater ipsilaterally than contralaterally implies that these alterations are driven by intrinsic factors specific to the (lateralized) TLE-HS syndrome rather than long-term effects of AEDs. Contrary to prior work (Ahmadi *et al*., 2009; Whelan *et al*., 2018), we did not find greater abnormalities in patients with left TLE relative to right TLE in either the TLE-HS or TLE-NL groups, nor did we find greater injury in males with TLE relative to females. Rather, men with TLE-HS and TLE-NL showed higher global FA values relative to women. This contrasts with a previous meta-analysis which found men with focal epilepsy to be more vulnerable to white matter injury relative to women (Slinger *et al*., 2016). These findings do not appear to reflect differences in age, age of seizure onset, or disease duration, as these characteristics did not differ between men and women in our TLE cohorts. It has been reported that compared to age-matched women, men have greater white matter volume and neuronal number with fewer neuronal processes (Rabinowicz *et al*., 1999), which is hypothesized to represent fewer, thicker and more organized fibers in men compared to more crossing fiber tracts in women (Schmithorst *et al*., 2008). This hypothesis is supported by the finding that higher FA in males compared to females was also observed in our healthy control sample.

Patients with TLE-NL showed a very mild pattern of white matter disruption compared to TLE-HS (Campos *et al*., 2015) (Fig. 3 and 4). Although this may, in part, reflect the greater likelihood for patients with TLE-NL to have a benign form of epilepsy, previous studies have shown that even among drug-resistant TLE, patients with TLE-NL harbor less severe cortical (Bernhardt *et al*., 2016, 2019) and white matter (Liu *et al*., 2012) abnormalities compared to TLE-HS. White matter disruptions in TLE-NL were notable in the EC and SS, both of which contain long-range association pathways. In particular, the SS contains fibers of the inferior fronto-occipital fasciculus and the inferior longitudinal fasciculus (Cristina *et al*., n.d.). These fibers course through the temporal lobe lateral to the CGH and FX/ST supporting data suggesting that TLE-HS and TLE-NL involve different epileptogenic networks (Zaveri *et al*., 2001; Mueller *et al*., 2009). However, many FA/MD alterations did not differ between TLE-HS and TLE-NL patients, once age of seizure onset and disease duration were taken into account (Fig. 5). Thus, the magnitude of these differences appears to be influenced by differences in clinical characteristics. This is supported by the association of an earlier age of seizure onset and longer disease duration with regional white matter disruption in TLE-HS, but not in TLE-NL.

### Genetic generalized epilepsy

GGE includes several related syndromes, defined electrographically by generalized, bisynchronous, and symmetric activity with spike-wave or polyspike-wave discharges (Weir, 1965; Seneviratne *et al*., 2012). These syndromes have traditionally been associated with thalamocortical dysfunction, with some studies reporting atrophy in the thalamus (Whelan *et al*., 2018) or thalamocortical networks (Bernhardt *et al*., 2009), and other studies reporting no structural changes relative to controls (McGill *et al*., 2014). Although patients with GGE showed modest alterations relative to patients with focal epilepsy across most fibers, these differences were broader than those previously observed (Slinger *et al*., 2016) and include commissural (GCC), projection (ACR) and corticocortical association pathways (EC, SLF). In addition, the magnitude of these changes in the ACR was similar to those observed in other epilepsy syndromes after adjusting for clinical and demographic characteristics (Fig. 5). The ACR is part of the limbic-thalamo-cortical circuitry and includes thalamic projections from the internal capsule to the cortex, including prominent connections to the frontal lobe bilaterally (Catani *et al*., 2002; Wakana *et al*., 2004). Thus, these projection fibers may be particularly affected in GGE, supporting their electrographic signature and previous work in subsets of patients with GGE, including juvenile myoclonic epilepsy suggestive of frontothalamic pathology (Woermann *et al*., 1999; Keller *et al*., 2011)

### Extratemporal epilepsy

The group of patients with non-lesional focal ExE showed the most marked alterations in the BCC, GCC, ACR, CGC, and EC, and showed a similar pattern to the GGE group of bilateral fronto-central alterations. As frontal lobe epilepsy is the second most common type of focal epilepsy (Manford *et al*., 1992), it is likely that this group dominated the ExE group, explaining the predominance of fronto-midline pathology. Unlike the TLE-HS, neither age of seizure onset nor disease duration were associated with regional white matter compromise. This may be related to the heterogeneity of this patient group, their latter age of seizure onset and/or the fact that this non-lesional group may represent a “mild” form of ExE.

### Summary

In summary, we found shared white matter compromise across epilepsy syndromes, dominated by regional alterations in bilateral midline, fiber bundles. The question arises as to whether these shared alterations are specific to epilepsy or are non-specific effects of a chronic brain disorder. Comparison of our ES to those of five other ENIGMA populations revealed that the pattern of microstructural compromise in epilepsy was similar to the patterns observed in BP, SCZ, and MDD. In particular, the CC body and genu were commonly affected across disorders, suggesting that microstructural compromise could reflect a shared patho-physiological mechanism. However, patients with epilepsy tend to have high rates of comorbid mood disorder (Kanner, 2007) and these were not characterized in our epilepsy sample. Therefore, overlap in neuropathology between our epilepsy cohort and major neuropsychiatric disorders could, in part, underlie the high rates of co-morbidity observed among these disorders. Increasing evidence suggests that neuropsychiatric disorders themselves are not separated by sharp neurobiological boundaries (Baker *et al*., 2019), but have overlapping of genetic influences and brain dysfunction (Brainstorm Consortium et al., 2018; Radonjić et al., 2019). Although genetic overlap between these neuropsychiatric disorders and epilepsy is low, overlap in dysfunctional brain networks may be partly due to comorbidities, and warrants further investigation.

### Limitations

First, although much effort was taken to apply post-acquisition harmonization, each scanner used varies in either image acquisition protocol or scanner hardware, or both, which increased methodological heterogeneity. Conversely, our results can be considered independent of any specific acquisition scheme, head coil or scanner model. Accordingly, while the absence of a single, standardized MR protocol incorporates scanner variance into the data, it also provides breadth that enhances the generalizability of findings. The statistical batch normalization process ComBat corrected differences between scanner instances, but may not adequately accommodate the heterogeneous neuropathology of epilepsy, resulting in a smoothing out of differences between syndromes..

A second limitation is the challenge of directly ascribing lower FA and higher MD to demyelination and/or axonal injury. Specifically, lower FA can reflect the effects of crossing fibers, increases in extracellular diffusion (e.g., inflammation, edema) or other technical or biological factors. Therefore, advanced diffusion sequences such as high angular resolution diffusion imaging (HARDI) and multishell dMRI acquisitions, together with analysis of quantitative contrasts sensitive to tissue microstructural features and neuropathological investigations, would help better unravel the biological underpinnings of our findings.

Finally, detailed clinical information was not available for all patients. Therefore, we could not directly assess how these diffusional changes were associated specific AED regimens, comorbid disorders, cognitive performances, or how they relate to clinical outcomes (i.e., drug resistance or post-operative seizure outcome). These data are now being collected across the consortium to better characterize patients and evaluate the clinical utility of identifying syndrome-specific and shared microstructural injury in epilepsy.

### Conclusions

Overall, in the largest DTI mega-analysis of epilepsy, we demonstrate a pattern of robust white matter alterations within and across patient syndromes, revealing shared and unique features for each syndrome. These microstructural alterations may reflect a number of pathological processes, including disrupted myelination, myelin or axonal loss due to chronic seizures, or increases in extracellular fluid due to inflammation. Such cross-syndrome and cross-disease comparisons could help to inform gene expression studies and provide novel insights into shared cognitive and psychiatric co-morbidities.

## Acknowledgments

Luis Concha: We extend our gratitude to the many people who have at some point participated in this study: Juan Ortíz-Retana, Erick Pasaye, Leopoldo González-Santos, Leticia Velázquez-Pérez, David Trejo, Héctor Barragán, Arturo Domínguez, Ildefonso Rodríguez-Leyva, Ana Luisa Velasco, Luis Octavio Jiménez, Daniel Atilano, Elizabeth González Olvera, Rafael Moreno, Vicente Camacho, Ana Elena Rosas, and Alfonso Fajardo.

## Funding

Sean N Hatton: NIH R01NS065838; NIH R21NS107739

Khoa H Huynh: None.

Leonardo Bonilha: R01 NS110347-01A1

Eugenio Abela: European Union Horizon 2020 research and innovation programme under the Marie Sklodowska-Curie grant agreement no.75088

Saud Alhusaini: This work was part funded by Science Foundation Ireland (16/RC/3948) and was cofunded under the European Regional Development Fund and by FutureNeuro industry partners

Andre Altmann: Dr Altmann holds an MRC eMedLab Medical Bioinformatics Career Development Fellowship. This work was partly supported by the Medical Research Council [grant number MR/L016311/ 1].

Marina KM Alvim: FAPESP 15/17066-0

Akshara R Balachandra: NIH R01NS065838

Emanuele Bartolini: None.

Benjamin Bender: None

Neda Bernasconi: None

Andrea Bernasconi: None

Boris Bernhardt: B.C.B. acknowledges support from CIHR (FDN-154298), SickKids Foundation (NI17-039), Natural Sciences and Engineering Research Council (NSERC; Discovery-1304413), Azrieli Center for Autism Research of the Montreal Neurological Institute (ACAR), and the Canada Research Chairs program.

Benoit Caldairou: None.

Maria Eugenia Caligiuri: None

Sarah JA Carr: None.

Gianpiero L Cavalleri: This work was part funded by Science Foundation Ireland (16/RC/3948) and was cofunded under the European Regional Development Fund and by FutureNeuro industry partners

Fernando Cendes: The UNICAMP research centre was funded by FAPESP (São Paulo Research Foundation); Contract grant number: 2013/07559-3.

Luis Concha: This work was supported by a grant from the Mexican Council of Science and Technology (CONACYT 181508); and from UNAM-DGAPA (IB201712). Imaging was performed at the National Laboratory for magnetic resonance imaging, which has received funding from CONACYT (grant numbers 232676, 251216, and 280283).

Esmaeil Davoodi-bojd: None.

Patricia M Desmond: None.

Orrin Devinsky: Finding A Cure for Epilepsy and Seizures

Colin P Doherty: None.

Martin Domin: None.

John S Duncan: National Institute of Health Research

Niels K Focke: DFG FO750/5-1

Sonya F Foley: CUBRIC, Cardiff University

Antonio Gambardella: None.

Ezequiel Gleichgerrcht: R01 NS110347-01A1

Khalid Hamandi: Health and Care Research Wales

Akaria Ishikawa: Fapesp-Brainn (2013/07559-3), Fapesp (2015/17335-0), Fapesp (2016/16355-0)

Simon S Keller: Medical Research Council (MR/S00355X/1 and MR/K023152/1) and Epilepsy Research UK (1085) grants

Peter V Kochunov: R01EB015611

Raviteja Kotikalapudi: None.

Barbara AK Kreilkamp: Epilepsy Action Postgraduate Research Bursary (Research Grants Programme 2014-2015)

Patrick Kwan: Patrick Kwan is supported by the MRFF Practitioner Fellowship

Angelo Labate: None.

Soenke Langner: None.

Matteo Lenge: None

Min Liu: None.

Elaine Lui: None.

Pascal Martin: P.M. was supported by the PATE program (F1315030) of the University of Tübingen.

Mario Mascalchi: None.

José CV Moreira: FAPESP-BRAINN (2013/07559-3)

Marcia E Morita-Sherman: NIH Grant

Terence J O’Brien: NHMRC Program Grant (#APP1091593)

Heath R Pardoe: None.

Jose C Pariente: None.

Letícia F Ribeiro: FAPESP-BRAINN (07559-3)

Mark P Richardson: Medical Research Council programme grant (MR/K013998/1); Medical Research Council Centre for Neurodevelopmental Disorders (MR/N026063/1); NIHR Biomedical Research Centre at South London and Maudsely NHS Foundation Trust

Cristiane S Rocha: FAPESP-BRAINN (07559-3)

Felix Rosenow: CePTER-Grant, LOEWE Programme, Minitster of Research and Arts, Hesse, Germany

Mariasavina Severino: None.

Benjamin Sinclair: Benjamin Sinclair is supported by Alfred Health, Monash University and Australian Commonwealth grant ICG000723.

Hamid Soltanian-Zadeh: None.

Pasquale Striano: None.

Peter N Taylor: None.

Rhys H Thomas: Epilepsy Research UK

Domenico Tortora: None

Dennis Velakoulis: None.

Annamaria Vezzani: Epitarget

Lucy Vivash: None.

Felix von Podewils: None.

Sjoerd B Vos: National Institute for Health Research University College London Hospitals Biomedical Research Centre (NIHR BRC UCLH/UCL High Impact Initiative BW.mn.BRC10269)

Bernd Weber: None.

Gavin P Winston: Medical Research Council (G0802012), National Institute for Health Research University College London Hospitals Biomedical Research Centre, Epilepsy Society

Clarissa LIN Yasuda: FAPESP-BRAINN (2013/07599-3), CNPQ (403726/2016-6)

Paul M Thompson: PT was funded in part by the United States NIH Big Data to Knowledge (BD2K) program under consortium grant U54 EB020403, the ENIGMA World Aging Center (R56 AG058854), and the ENIGMA Sex Differences Initiative (R01 MH116147).

Neda Jahanshad: R01MH117601, R01AG059874, R01MH121246, U54EB020403, P41EB015922

Sanjay M Sisodiya: The work was partly undertaken at UCLH/UCL, which received a proportion of funding from the Department of Health’s NIHR Biomedical Research Centres funding scheme. We are grateful to the Wolfson Trust and the Epilepsy Society for supporting the Epilepsy Society MRI scanner.

Carrie R McDonald: NIH R01NS065838; NIH R21NS107739

## Footnotes

1 A *meta-analysis* aggregate of summary results (e.g. effect size estimates, standard errors, and confidence intervals) across studies, but a *mega-analysis* aggregates individual participant data across studies, and may allow additional data harmonization. For an empirical comparison between the two techniques using structural MRI data, please refer to (Boedhoe *et al*., 2018).

