## Supplemental Material for "White matter abnormalities across different epilepsy syndromes in adults: an ENIGMA Epilepsy study"

Supplementary Information

**Supplementary Table 1.** Scanner protocols by each research center.

**Supplementary Table 2.** Results of t-test for age differences, and chi-squared test for sex differences at each research center.

**Supplementary Table 3.** Average age of onset and duration of illness per research center.

**Supplementary Table 4.** Multivariate tests.

**Supplementary Figure 1.** RD effect size graphs.

**Supplementary Figure 2.** AD effect size graphs.

**Supplementary Table 5.** Effect sizes for FA differences between healthy controls and the ‘All Epilepsies’ syndrome.

**Supplementary Table 6.** Effect sizes for FA differences between healthy controls and the ‘L TLE-HS’ syndrome.

**Supplementary Table 7.** Effect sizes for FA differences between healthy controls and the ‘R TLE-HS’ syndrome.

**Supplementary Table 8.** Effect sizes for FA differences between healthy controls and the ‘L TLE-NL’ syndrome.

**Supplementary Table 9.** Effect sizes for FA differences between healthy controls and the ‘R TLE-NL’ syndrome.

**Supplementary Table 10.** Effect sizes for FA differences between healthy controls and the ‘GGE’ syndrome.

**Supplementary Table 11.** Effect sizes for FA differences between healthy controls and the ‘ExE’ syndrome.

**Supplementary Table 12.** Effect sizes for MD differences between healthy controls and the ‘All Epilepsies’ syndrome.

**Supplementary Table 13.** Effect sizes for MD differences between healthy controls and the ‘L TLE-HS’ syndrome.

**Supplementary Table 14.** Effect sizes for MD differences between healthy controls and the ‘R TLE-HS’ syndrome.

**Supplementary Table 15.** Effect sizes for MD differences between healthy controls and the ‘L TLE-NL’ syndrome.

**Supplementary Table 16.** Effect sizes for MD differences between healthy controls and the ‘R TLE-NL’ syndrome.

**Supplementary Table 17.** Effect sizes for MD differences between healthy controls and the ‘GGE’ syndrome.

**Supplementary Table 19.** Effect sizes for MD differences between healthy controls and the ‘ExE’ syndrome.

**Supplementary Table 20.** Effect sizes for RD differences between healthy controls and the 'All Epilepsies' syndrome.

**Supplementary Table 21.** Effect sizes for RD differences between healthy controls and the 'L TLE-HS' syndrome.

**Supplementary Table 22.** Effect sizes for RD differences between healthy controls and the 'R TLE-HS' syndrome.

**Supplementary Table 23.** Effect sizes for RD differences between healthy controls and the 'L TLE-NL' syndrome.

**Supplementary Table 24.** Effect sizes for RD differences between healthy controls and the 'R TLE-NL' syndrome.

**Supplementary Table 25.** Effect sizes for RD differences between healthy controls and the 'GGE' syndrome.

**Supplementary Table 26.** Effect sizes for RD differences between healthy controls and the 'ExE' syndrome.

**Supplementary Table 27.** Effect sizes for AD differences between healthy controls and the 'All Epilepsies' syndrome.

**Supplementary Table 28.** Effect sizes for AD differences between healthy controls and the 'L TLE-HS' syndrome.

**Supplementary Table 29.** Effect sizes for AD differences between healthy controls and the 'R TLE-HS' syndrome.

**Supplementary Table 30.** Effect sizes for AD differences between healthy controls and the 'L TLE-NL' syndrome.

**Supplementary Table 31.** Effect sizes for AD differences between healthy controls and the 'R TLE-NL' syndrome.

**Supplementary Table 32.** Effect sizes for AD differences between healthy controls and the 'GGE' syndrome.

**Supplementary Table 33.** Effect sizes for AD differences between healthy controls and the 'ExE' syndrome.

**Supplementary Table 34.** Relationship between FA and age of disease onset by syndrome.

**Supplementary Table 35.** Relationship between FA and disease duration by syndrome.

**Supplementary Table 36.** Relationship between MD and age of disease onset by syndrome.

**Supplementary Table 37.** Relationship between MD and disease duration by syndrome.

**Supplementary Table 38.** Relationship between RD and age of disease onset by syndrome.

**Supplementary Table 39.** Relationship between RD and disease duration by syndrome.

**Supplementary Table 40.** Relationship between AD and age of disease onset by syndrome.

**Supplementary Table 41.** Relationship between AD and disease duration by syndrome.

**Supplementary Table 1.** Scanner protocols by each research center. An asterisk (*) indicates a variable TR due to cardiac gating.

| Center | Scanner | Orientation | # of Slices | Voxel Size (mm3) | Gradient Directions | b-value (mm/s2) | # b=0 scans | TE (ms) | TR (ms) | Relevant Citation |
|---|---|---|---|---|---|---|---|---|---|---|
| **Bonn** | Siemens Trio | Axial | 160 | 1 x 1 x 1 | 60 | 1000 | 7 | 3.97 | 1300 | Kreilkamp, B. A., Weber, B., Richardson, M. P., & Keller, S. S. (2017). Automated tractography in patients with temporal lobe epilepsy using TRActs Constrained by UnderLying Anatomy (TRACULA). *NeuroImage: Clinical,14*, 67-76. doi:10.1016/j.nicl.2017.01.003 |
| **CUBRIC** | GE Signa HDx | - | 60 | 2.4 mm slice thickness | 30 | 1200 | 3 | 87 | * | Caeyenberghs, K., Powell, H., Thomas, R., Brindley, L., Church, C., Evans, J., . . . Hamandi, K. (2015). Hyperconnectivity in juvenile myoclonic epilepsy: A network analysis. *NeuroImage: Clinical,7*, 98-104. doi:10.1016/j.nicl.2014.11.018 |
| **EKUT** | Siemens Trio | - | 52 | 1.81 x 1.81 x 1.79 | 48 | 1200 (2x) | 6 (2x) | 93 | 9400 | - |
| **EPICZ** | GE Discovery MR750 | Axial | 80 | 2 x 2 x 2 | 27 | 1000 | 4 | 81.4 | 10000 | Caligiuri, M. E., Labate, A., Cherubini, A., Mumoli, L., Ferlazzo, E., Aguglia, U., . . . Gambardella, A. (2016). Integrity of the corpus callosum in patients with benign temporal lobe epilepsy. *Epilepsia,57*(4), 590-596. doi:10.1111/epi.13339 |
| **EPIGEN-Ireland** | Philips Achieva | Axial | 70 | 1.75 x 1.75 x 2 | 32 | 1000 | - | 52 | 12786 | Whelan, C. D., Alhusaini, S., Ohanlon, E., Cheung, M., Iyer, P. M., Meaney, J. F., . . . Cavalleri, G. L. (2015). White matter alterations in patients with MRI-negative temporal lobe epilepsy and their asymptomatic siblings. *Epilepsia,56*(10), 1551-1561. doi:10.1111/epi.13103 |
| **Florence** | Philips Achieva | - | 69 | 2 x 2 x 2 | 32 | 1000 | 1 | 80 | 4000 | - |
| **Genova** | Philips Ingenia | Axial | 65 | 2 x 2 x 2 | 64 | 1000 | 1 | 90 | 7000 | - |
| **Greifswald** | Siemens Verio | - | 80 | 1.8 x 1.8 x 1.8 | 64 | 1000 | 1 | 107 | 15300 | Domin, M., Bartels, S., Geithner, J., Wang, Z. I., Runge, U., Grothe, M., . . . Podewils, F. V. (2018). Juvenile Myoclonic Epilepsy Shows Potential Structural White Matter Abnormalities: A TBSS |

| | | | | | | | | | | |
|---|---|---|---|---|---|---|---|---|---|---|
| | | | | | | | | | | Study. *Frontiers in Neurology,9*. doi:10.3389/fneur.2018.00509 |
| **Henry Ford** | GE Signa | Axial | 60 | 1.96 × 1.96 × 2.6 | 25 | 1000 | 1 | 76 | 7500 | Nazem-Zadeh, M., Bowyer, S. M., Moran, J. E., Davoodi-Bojd, E., Zillgitt, A., Weiland, B. J., . . . Soltanian-Zadeh, H. (2016). MEG Coherence and DTI Connectivity in mTLE. *Brain Topography,29*(4), 598-622. doi:10.1007/s10548-016-0488-0 |
| **IDIBAPS_31DIR** | Siemens Trio | Axial | 55 | 2.4 x 2.4 x 2.4 | 30 | 1000 | 1 | 90 | 6900 | |
| **IDIBAPS_39DIR** | Siemens Trio | Axial | 64 | 1.97 x 1.97 x 2 | 36 | 1000 | 3 | 88 | 8138 | Córdova-Palomera, A., Reus, M. A., Fatjó-Vilas, M., Falcón, C., Bargalló, N., Heuvel, M. P., & Fañanás, L. (2016). FKBP5 modulates the hippocampal connectivity deficits in depression: A study in twins. *Brain Imaging and Behavior,11*(1), 62-75. doi:10.1007/s11682-015-9503-4 |
| **IDIBAPS_88DIR** | Siemens Trio | Axial | 55 | 1.25 x 1.25 x 2.5 | 82 | 1000 | 6 | 98 | 7600 | Aparicio, J., Carreño, M., Bargalló, N., Setoain, X., Rubí, S., Rumià, J., . . . Donaire, A. (2016). Combined 18 F-FDG-PET and diffusion tensor imaging in mesial temporal lobe epilepsy with hippocampal sclerosis. *NeuroImage: Clinical,12*, 976-989. doi:10.1016/j.nicl.2016.05.002 |
| **KCL** | GE Discovery MR750 | Axial | 66 | 2.4 x 2.4 x 2.4 | 32 | 1000 | 6 | 75 | * | - |
| **Liverpool_Walton** | GE Discovery MR750 | Axial | 66 | 1 x 1 x 2 | 60 | 1000 | 6 | 82 | 8000 | Kreilkamp, B. A., Weber, B., Richardson, M. P., & Keller, S. S. (2017). Automated tractography in patients with temporal lobe epilepsy using TRActs Constrained by UnderLying Anatomy (TRACULA). *NeuroImage: Clinical,14*, 67-76. doi:10.1016/j.nicl.2017.01.003 |
| **MNI** | Siemens Trio | Axial | 63 | 2 x 2 x 2 | 64 | 1000 | 1 | 90 | 8400 | Liu, M., Bernhardt, B. C., Hong, S., Caldairou, B., Bernasconi, A., & Bernasconi, N. (2016). The superficial white matter in temporal lobe epilepsy: A key link between structural and functional network disruptions. *Brain,139*(9), 2431-2440. doi:10.1093/brain/aww167 |
| **NYU** | Siemens Allegra | Axial | 60 | 2.5 x 2.5 x 2.5 | 64 | 3000 | 8 | 99 | 7900 | - |

| | | | | | | | | | | |
|---|---|---|---|---|---|---|---|---|---|---|
| **Melbourne** | Siemens Trio | Axial | 55 | 2.5 x 2.5 x 2.5 | 64 | 3000 | 1 | 122 | 8700 | - |
| **UCL** | GE Signa HDx | Axial | 60 | 1.875×1.875×2.4 | 52 | 1200 | 6 | 73 | * | Taylor, P. N., Sinha, N., Wang, Y., Vos, S. B., Tisi, J. D., Miserocchi, A., . . . Duncan, J. S. (2018). The impact of epilepsy surgery on the structural connectome and its relation to outcome. *NeuroImage: Clinical*,*18*, 202-214. doi:10.1016/j.nicl.2018.01.028 |
| **UCSD** | GE Discovery MR750 | Axial | 53 | 1.86 x 1.86 x 2.5 | 30 | 1000 | 2 | 82.9 | 8000 | Reyes, A., Paul, B. M., Marshall, A., Chang, Y. A., Bahrami, N., Kansal, L., . . . Mcdonald, C. R. (2018). Does bilingualism increase brain or cognitive reserve in patients with temporal lobe epilepsy? *Epilepsia*,*59*(5), 1037-1047. doi:10.1111/epi.14072 |
| **UMG** | Siemens Trio | - | 31 | 1.89 x 1.89 x 1.89 | 30 | 1000 | 1 | 93 | 10000 | Bonilha, L., Gleichgerrcht, E., Fridriksson, J., Rorden, C., Breedlove, J. L., Nesland, T., . . . Focke, N. K. (2015). Reproducibility of the Structural Brain Connectome Derived from Diffusion Tensor Imaging. *Plos One*,*10*(9). doi:10.1371/journal.pone.0135247 |
| **UNAM** | Philips Achieva | - | - | 2 x 2 x 2 | 60 | 2000 | 1 | 64.3 | 11860 | Rodríguez-Cruces, R., Velázquez-Pérez, L., Rodríguez-Leyva, I., Velasco, A. L., Trejo-Martínez, D., Barragán-Campos, H. M., . . . Concha, L. (2018). Association of white matter diffusion characteristics and cognitive deficits in temporal lobe epilepsy. *Epilepsy & Behavior*,*79*, 138-145. doi:10.1016/j.yebeh.2017.11.040 |
| **UNICAMP** | Philips Achieva | Axial | 70 | 2 x 2 x 2 | 32 | 1000 | 1 | 61 | 8500 | Campos, B. M., Coan, A. C., Beltramini, G. C., Liu, M., Yassuda, C. L., Ghizoni, E., . . . Cendes, F. (2014). White matter abnormalities associate with type and localization of focal epileptogenic lesions. *Epilepsia*,*56*(1), 125-132. doi:10.1111/epi.12871 |

**Supplementary Table 2.** Results of t-test for age differences, and chi-squared test for sex differences at each research center.

| | T-test for age differences | | | | Chi squared test for sex differences | |
|---|---|---|---|---|---|---|
| **Site** | **Difference of means** | **Standard error** | **T-statistic** | **P-value** | **Pearson chi-squared value** | **P-value** |
| **Bonn** | 3.81 | 2.65 | -1.44 | 0.15 | 0.59 | 0.44 |
| **CUBRIC** | 0.39 | 1.94 | -0.20 | 0.84 | 0.01 | 0.94 |
| **EKUT** | -6.08 | 4.80 | 1.27 | 0.22 | 0.68 | 0.41 |
| **EPICZ** | -0.31 | 1.40 | 0.22 | 0.83 | 3.92 | 0.05 |
| **EPIGEN_Ireland** | 1.15 | 1.67 | -0.69 | 0.49 | 5.24 | 0.02 |
| **Florence** | 3.02 | 5.14 | -0.59 | 0.57 | 0.32 | 0.57 |
| **Genova** | 0.00 | 2.60 | 0.00 | 1.00 | 0.00 | 1.00 |
| **Greifswald** | - | - | - | - | - | - |
| **HFHS_2.6mm** | 8.75 | 2.09 | -4.19 | 9.3E-05 | 2.97 | 0.08 |
| **HFHS_3mm** | - | - | - | - | - | - |
| **IDIBAPS_31DIR** | - | - | - | - | - | - |
| **IDIBAPS_39DIR** | 0.96 | 1.79 | -0.48 | 0.63 | 2.00 | 0.16 |
| **IDIBAPS_88DIR** | 3.47 | 1.73 | -2.01 | 0.05 | 0.14 | 0.70 |
| **KCL** | 4.64 | 1.27 | -3.64 | 0.00 | 0.92 | 0.34 |
| **LiverpoolWalton** | -0.74 | 2.10 | 0.35 | 0.72 | 0.10 | 0.75 |
| **Melbourne** | - | - | - | - | - | - |
| **MNI** | 1.32 | 1.32 | -0.99 | 0.32 | 0.14 | 0.71 |
| **MUSC** | -18.50 | 2.00 | 9.25 | 3.4E-14 | 1.92 | 0.17 |
| **NYU** | 2.27 | 2.36 | -0.96 | 0.34 | 0.32 | 0.57 |
| **UCL** | 1.00 | 2.78 | -0.36 | 0.72 | 0.10 | 0.75 |
| **UCSD** | -3.26 | 2.53 | 1.29 | 0.20 | 0.37 | 0.54 |
| **UMG** | -1.94 | 2.77 | 0.70 | 0.49 | 0.82 | 0.36 |
| **UNAM** | -2.26 | 2.89 | 0.78 | 0.44 | 0.28 | 0.59 |
| **UNICAMP** | 5.04 | 0.89 | -5.66 | 2.5E-08 | 0.27 | 0.60 |

**Supplementary Table 3.** Average age of onset and duration of illness per research center.

| | Age of Onset | | | | | | | Duration of Illness | | | | | | |
|---|---|---|---|---|---|---|---|---|---|---|---|---|---|---|
| **Site** | **All Epilepsies** | **GGE** | **TLE-HS L** | **TLE-HS R** | **TLE-NL L** | **TLE-NL R** | **ExE** | **All Epilepsies** | **GGE** | **TLE-HS L** | **TLE-HS R** | **TLE-NL L** | **TLE-NL R** | **ExE** |
| **Bonn** | 17.7 | - | 18.3 | 16.5 | - | - | - | 23.8 | - | 23.5 | 24.6 | - | - | - |
| **CUBRIC** | 14.7 | 14.7 | - | - | - | - | - | 14.0 | 14.0 | - | - | - | - | - |
| **EKUT** | 19.3 | 19.3 | - | - | - | - | - | 14.6 | 14.6 | - | - | - | - | - |
| **EPICZ** | 19.1 | - | 19.4 | 17.5 | 17.5 | 25.7 | - | 19.2 | - | 18.3 | 23.9 | 19.7 | 13.4 | - |
| **EPIGEN_Ireland** | 19.1 | - | 20.2 | 13.3 | 20.2 | 16.4 | - | 18.1 | - | 20.7 | 25.7 | 15.7 | 18.5 | - |
| **Florence** | 14.0 | - | - | - | - | 14.0 | 14.0 | 19.0 | - | - | - | - | 22.5 | 15.5 |
| **Genova** | 10.4 | 12.1 | - | 5.0 | 11.0 | 7.3 | 10.0 | 14.7 | 16.4 | - | 13.0 | 12.0 | 11.3 | 9.5 |
| **Greifswald** | 19.2 | 14.5 | - | - | - | - | 30.2 | 15.4 | 14.8 | - | - | - | - | 18.0 |
| **HFHS_2.6mm** | 19.0 | - | 13.2 | 8.3 | 18.8 | 22.7 | 17.4 | 19.0 | - | 21.3 | 28.4 | 20.9 | 19.3 | 19.6 |
| **HFHS_3mm** | - | - | - | - | - | - | - | - | - | - | - | - | - | - |
| **IDIBAPS_31Dir** | 22.5 | 8.0 | 22.3 | 24.1 | 17.7 | 16.0 | 24.3 | 16.4 | 13.0 | 17.3 | 17.4 | 20.7 | 27.0 | 13.6 |
| **IDIBAPS_39Dir** | 16.9 | - | 18.5 | 18.0 | - | 17.0 | 14.0 | 17.6 | - | 27.0 | 12.2 | - | 28.0 | 12.5 |
| **IDIBAPS_88Dir** | 15.6 | - | 15.5 | 16.3 | 19.3 | 14.0 | 16.8 | 22.0 | - | 25.5 | 17.5 | 20.0 | 24.7 | 11.7 |
| **KCL** | - | - | - | - | - | - | - | - | - | - | - | - | - | - |
| **LiverpoolWalton** | 14.3 | 19.0 | 9.0 | 15.7 | 14.0 | 19.6 | 11.4 | 16.7 | 9.0 | 24.8 | 23.2 | 14.6 | 8.0 | 17.6 |
| **Melbourne** | 22.4 | - | 15.2 | 31.4 | 26.8 | 28.0 | - | 14.8 | - | 20.8 | 8.4 | 7.0 | 9.0 | - |
| **MNI** | 16.4 | - | 14.6 | 14.3 | 21.8 | 18.8 | 15.4 | 15.4 | - | 17.1 | 24.2 | 14.0 | 13.2 | 12.8 |
| **MUSC** | 18.8 | - | 14.9 | 21.6 | 20.8 | 24.0 | - | 18.2 | - | 20.8 | 13.6 | 18.3 | 14.8 | - |
| **NYU** | 24.0 | - | 19.5 | 10.0 | 30.6 | 28.7 | 22.8 | 8.6 | - | 13.0 | 13.5 | 7.3 | 2.7 | 9.1 |
| **UCL** | 13.9 | - | 12.0 | 11.3 | 22.5 | 16.9 | - | 24.7 | - | 26.4 | 30.1 | 13.3 | 20.5 | - |
| **UCSD** | 19.6 | - | 20.9 | 19.8 | 17.3 | 19.9 | - | 15.2 | - | 15.1 | 16.0 | 17.4 | 13.5 | - |
| **UMG** | 16.6 | 15.3 | 31.0 | 9.3 | - | - | 20.4 | 15.6 | 16.3 | 11.0 | 10.0 | - | - | 15.6 |
| **UNAM** | 15.8 | - | 16.4 | 12.5 | 17.9 | 16.0 | - | 15.5 | - | 17.6 | 21.0 | 13.4 | 2.5 | - |
| **UNICAMP** | 12.3 | 13.5 | 10.0 | 13.4 | 10.3 | 17.5 | 10.1 | 28.3 | 22.8 | 31.6 | 30.5 | 26.9 | 25.0 | 23.5 |
| **Weighted Average** | 16.6 | 14.8 | 14.5 | 15.3 | 18.8 | 19.5 | 17.6 | 19.9 | 16.8 | 24.4 | 24.4 | 17.0 | 15.6 | 15.1 |

**Supplementary Table 4. Multivariate tests.** Increasing values of Pillai's trace indicate effects that contribute more to the model.

| Measure | Effect | Pillai's Trace | F | Hypothesis df | Error df | Sig. | Partial $\eta^2$ | Observed Power |
|---|---|---|---|---|---|---|---|---|
| FA | Diagnosis | 0.44 | 4.70 | 228 | 13452 | <0.001 | 0.074 | 1.0 |
| | Age | 0.05 | 2.90 | 38 | 2237 | <0.001 | 0.047 | 1.0 |
| | $Age^2$ | 0.04 | 2.33 | 38 | 2237 | <0.001 | 0.038 | 1.0 |
| | Sex | 0.14 | 9.27 | 38 | 2237 | <0.001 | 0.136 | 1.0 |
| | Sex*Diagnosis | 0.13 | 1.33 | 228 | 13416 | 0.001 | 0.022 | 1.0 |
| MD | Diagnosis | 0.31 | 2.80 | 228 | 11688 | <0.001 | 0.052 | 1.0 |
| | Age | 0.07 | 3.67 | 38 | 1943 | <0.001 | 0.067 | 1.0 |
| | $Age^2$ | 0.05 | 2.94 | 38 | 1943 | <0.001 | 0.054 | 1.0 |
| | Sex | 0.12 | 6.77 | 38 | 1943 | <0.001 | 0.117 | 1.0 |
| | Sex*Diagnosis | 0.13 | 1.08 | 228 | 11652 | 0.870 | 0.021 | 1.0 |
| AD | Diagnosis | 0.22 | 1.96 | 228 | 11946 | <0.001 | 0.036 | 1.0 |
| | Age | 0.07 | 4.05 | 38 | 1986 | <0.001 | 0.072 | 1.0 |
| | $Age^2$ | 0.05 | 2.98 | 38 | 1986 | <0.001 | 0.054 | 1.0 |
| | Sex | 0.10 | 5.77 | 38 | 1986 | <0.001 | 0.099 | 1.0 |
| | Sex*Diagnosis | 0.12 | 1.07 | 228 | 11910 | 0.215 | 0.020 | 1.0 |
| RD | Diagnosis | 0.36 | 3.28 | 228 | 11790 | <0.001 | 0.060 | 1.0 |
| | Age | 0.07 | 4.16 | 38 | 1960 | <.001 | 0.075 | 1.0 |
| | $Age^2$ | 0.06 | 3.58 | 38 | 1960 | <.001 | 0.065 | 1.0 |
| | Sex | 0.11 | 6.65 | 38 | 1960 | <.001 | 0.114 | 1.0 |
| | Sex*Diagnosis | 0.13 | 1.18 | 228 | 11754 | 0.033 | 0.022 | 1.0 |

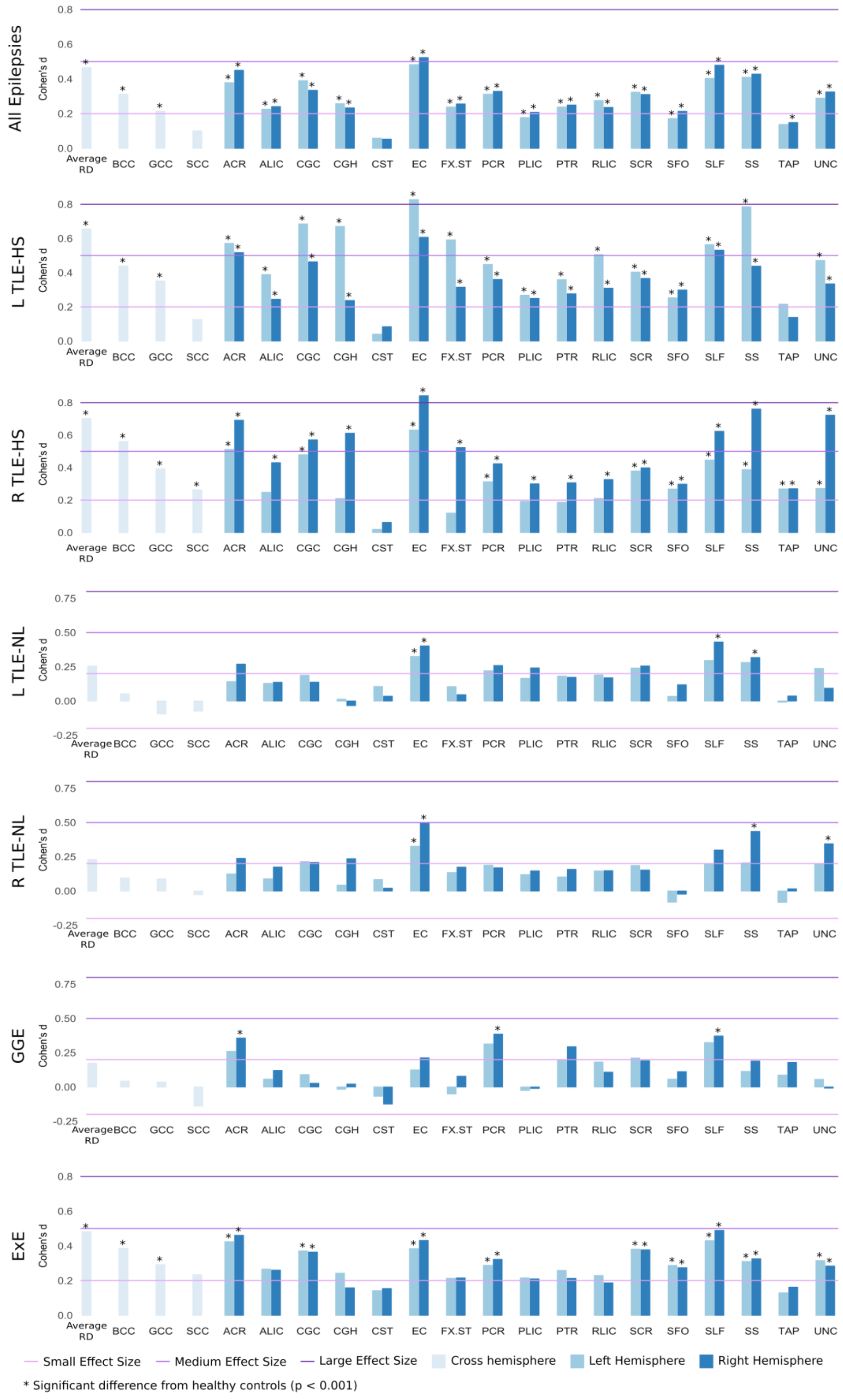


**Supplementary Figure 1.** RD effect size graphs.

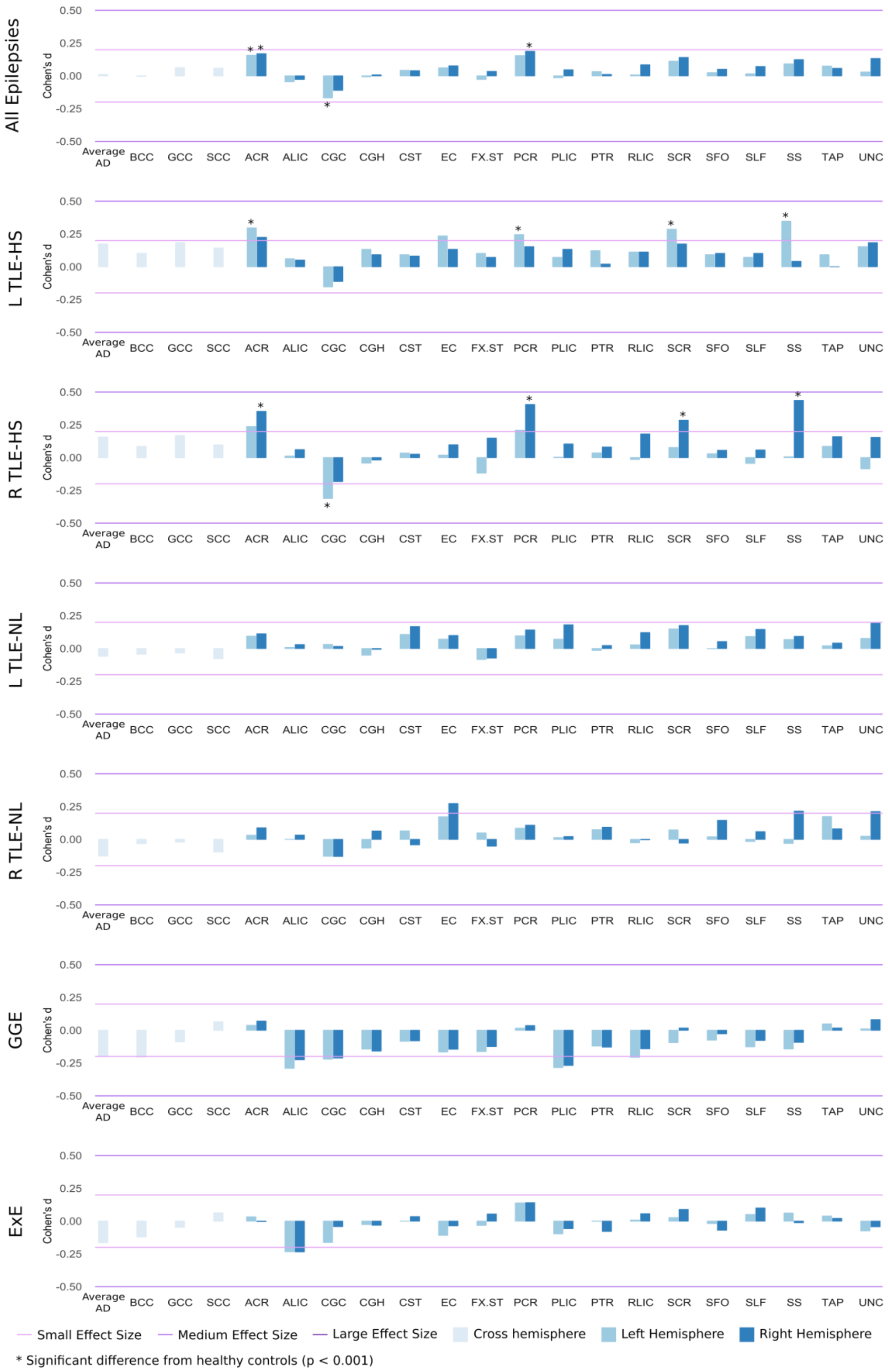


**Supplementary Figure 2.** AD effect size graphs.

**Supplementary Table 5.** Effect sizes for FA differences between healthy controls and the 'All Epilepsies' syndrome.

| ROI | Mean FA controls | Mean FA patients | Cohen's d | 95% CI lower | 95% CI upper | Standard error | Z score | P value |
|---|---|---|---|---|---|---|---|---|
| **AverageFA** | 0.585 | 0.569 | -0.71 | -0.79 | -0.62 | 0.04 | -24.62 | 2.3E-59 |
| **BCC** | 0.690 | 0.668 | -0.59 | -0.68 | -0.51 | 0.04 | -20.69 | 1.3E-45 |
| **GCC** | 0.695 | 0.674 | -0.59 | -0.68 | -0.51 | 0.04 | -20.66 | 1.5E-43 |
| **SCC** | 0.762 | 0.752 | -0.36 | -0.44 | -0.27 | 0.04 | -12.40 | 4.9E-17 |
| **ACR.L** | 0.479 | 0.463 | -0.50 | -0.58 | -0.41 | 0.04 | -17.31 | 3.0E-32 |
| **ACR.R** | 0.484 | 0.467 | -0.52 | -0.60 | -0.43 | 0.04 | -17.99 | 1.2E-33 |
| **ALIC.L** | 0.600 | 0.587 | -0.48 | -0.56 | -0.39 | 0.04 | -16.61 | 7.0E-26 |
| **ALIC.R** | 0.607 | 0.597 | -0.34 | -0.42 | -0.25 | 0.04 | -11.76 | 1.8E-15 |
| **CGC.L** | 0.648 | 0.624 | -0.57 | -0.65 | -0.48 | 0.04 | -19.73 | 2.7E-38 |
| **CGC.R** | 0.604 | 0.583 | -0.50 | -0.59 | -0.42 | 0.04 | -17.62 | 8.3E-34 |
| **CGH.L** | 0.553 | 0.527 | -0.52 | -0.60 | -0.43 | 0.04 | -18.10 | 3.3E-35 |
| **CGH.R** | 0.565 | 0.540 | -0.48 | -0.57 | -0.40 | 0.04 | -16.93 | 6.0E-30 |
| **CST.L** | 0.624 | 0.620 | -0.09 | -0.17 | -0.01 | 0.04 | -3.22 | 0.009 |
| **CST.R** | 0.614 | 0.610 | -0.10 | -0.18 | -0.01 | 0.04 | -3.35 | 0.021 |
| **EC.L** | 0.484 | 0.465 | -0.64 | -0.73 | -0.56 | 0.04 | -22.43 | 5.3E-53 |
| **EC.R** | 0.483 | 0.465 | -0.63 | -0.72 | -0.54 | 0.04 | -22.01 | 2.7E-47 |
| **FX.ST.L** | 0.580 | 0.564 | -0.41 | -0.49 | -0.33 | 0.04 | -14.33 | 1.6E-23 |
| **FX.ST.R** | 0.579 | 0.563 | -0.40 | -0.48 | -0.31 | 0.04 | -13.93 | 8.5E-21 |
| **PCR.L** | 0.506 | 0.494 | -0.38 | -0.46 | -0.30 | 0.04 | -13.25 | 1.9E-21 |
| **PCR.R** | 0.512 | 0.500 | -0.41 | -0.50 | -0.33 | 0.04 | -14.43 | 4.9E-23 |
| **PLIC.L** | 0.697 | 0.692 | -0.17 | -0.26 | -0.09 | 0.04 | -6.07 | 7.0E-06 |
| **PLIC.R** | 0.704 | 0.699 | -0.18 | -0.26 | -0.10 | 0.04 | -6.23 | 1.4E-05 |
| **PTR.L** | 0.625 | 0.611 | -0.40 | -0.48 | -0.32 | 0.04 | -14.00 | 5.3E-20 |
| **PTR.R** | 0.629 | 0.614 | -0.41 | -0.50 | -0.33 | 0.04 | -14.35 | 2.0E-21 |
| **RLIC.L** | 0.616 | 0.603 | -0.40 | -0.49 | -0.32 | 0.04 | -14.08 | 1.4E-20 |
| **RLIC.R** | 0.600 | 0.588 | -0.37 | -0.46 | -0.29 | 0.04 | -13.03 | 1.8E-17 |
| **SCR.L** | 0.516 | 0.505 | -0.42 | -0.50 | -0.33 | 0.04 | -14.60 | 3.0E-22 |
| **SCR.R** | 0.511 | 0.501 | -0.39 | -0.47 | -0.30 | 0.04 | -13.45 | 3.1E-18 |
| **SFO.L** | 0.549 | 0.535 | -0.34 | -0.42 | -0.26 | 0.04 | -11.88 | 1.1E-15 |
| **SFO.R** | 0.552 | 0.542 | -0.26 | -0.34 | -0.18 | 0.04 | -9.02 | 1.6E-09 |
| **SLF.L** | 0.529 | 0.516 | -0.47 | -0.55 | -0.38 | 0.04 | -16.24 | 1.5E-28 |
| **SLF.R** | 0.527 | 0.513 | -0.48 | -0.57 | -0.40 | 0.04 | -16.84 | 2.0E-32 |
| **SS.L** | 0.578 | 0.560 | -0.55 | -0.63 | -0.46 | 0.04 | -19.04 | 6.4E-36 |
| **SS.R** | 0.579 | 0.560 | -0.52 | -0.60 | -0.43 | 0.04 | -18.13 | 6.3E-35 |
| **TAP.L** | 0.571 | 0.552 | -0.35 | -0.43 | -0.26 | 0.04 | -12.06 | 2.8E-15 |
| **TAP.R** | 0.603 | 0.587 | -0.29 | -0.37 | -0.21 | 0.04 | -10.09 | 1.4E-12 |
| **UNC.L** | 0.543 | 0.525 | -0.33 | -0.42 | -0.25 | 0.04 | -11.64 | 4.1E-16 |
| **UNC.R** | 0.563 | 0.541 | -0.40 | -0.48 | -0.31 | 0.04 | -13.80 | 2.1E-20 |

**Supplementary Table 6.** Effect sizes for FA differences between healthy controls and the 'L TLE-HS' syndrome.

| ROI | Mean FA controls | Mean FA patients | Cohen's d | 95% CI lower | 95% CI upper | Standard error | Z score | P value |
|---|---|---|---|---|---|---|---|---|
| **AverageFA** | 0.585 | 0.565 | -0.92 | -1.05 | -0.79 | 0.07 | -16.49 | 6.6E-40 |
| **BCC** | 0.690 | 0.666 | -0.75 | -0.88 | -0.62 | 0.07 | -13.35 | 1.0E-26 |
| **GCC** | 0.695 | 0.670 | -0.79 | -0.92 | -0.66 | 0.07 | -14.11 | 2.6E-28 |
| **SCC** | 0.762 | 0.751 | -0.48 | -0.60 | -0.35 | 0.07 | -8.48 | 1.7E-10 |
| **ACR.L** | 0.479 | 0.459 | -0.66 | -0.79 | -0.53 | 0.07 | -11.83 | 1.1E-21 |
| **ACR.R** | 0.484 | 0.467 | -0.53 | -0.66 | -0.40 | 0.07 | -9.41 | 3.7E-14 |
| **ALIC.L** | 0.600 | 0.583 | -0.61 | -0.74 | -0.48 | 0.07 | -10.92 | 5.6E-20 |
| **ALIC.R** | 0.607 | 0.599 | -0.28 | -0.41 | -0.16 | 0.06 | -5.04 | 9.8E-06 |
| **CGC.L** | 0.648 | 0.613 | -0.89 | -1.02 | -0.76 | 0.07 | -15.87 | 1.6E-36 |
| **CGC.R** | 0.604 | 0.578 | -0.68 | -0.81 | -0.55 | 0.07 | -12.06 | 2.6E-24 |
| **CGH.L** | 0.553 | 0.507 | -1.01 | -1.15 | -0.88 | 0.07 | -18.07 | 1.4E-50 |
| **CGH.R** | 0.565 | 0.546 | -0.43 | -0.56 | -0.30 | 0.07 | -7.63 | 6.0E-11 |
| **CST.L** | 0.624 | 0.623 | -0.05 | -0.17 | 0.08 | 0.06 | -0.80 | 0.319 |
| **CST.R** | 0.614 | 0.611 | -0.10 | -0.22 | 0.03 | 0.06 | -1.71 | 0.099 |
| **EC.L** | 0.484 | 0.456 | -1.02 | -1.15 | -0.89 | 0.07 | -18.26 | 4.6E-49 |
| **EC.R** | 0.483 | 0.465 | -0.68 | -0.81 | -0.55 | 0.07 | -12.11 | 4.7E-25 |
| **FX.ST.L** | 0.580 | 0.550 | -0.83 | -0.96 | -0.70 | 0.07 | -14.81 | 4.8E-33 |
| **FX.ST.R** | 0.579 | 0.561 | -0.47 | -0.60 | -0.35 | 0.07 | -8.39 | 1.1E-12 |
| **PCR.L** | 0.506 | 0.490 | -0.54 | -0.67 | -0.41 | 0.07 | -9.54 | 1.9E-14 |
| **PCR.R** | 0.512 | 0.498 | -0.46 | -0.59 | -0.34 | 0.07 | -8.22 | 1.1E-11 |
| **PLIC.L** | 0.697 | 0.690 | -0.25 | -0.38 | -0.13 | 0.06 | -4.49 | 9.9E-05 |
| **PLIC.R** | 0.704 | 0.699 | -0.15 | -0.27 | -0.02 | 0.06 | -2.62 | 0.009 |
| **PTR.L** | 0.625 | 0.608 | -0.48 | -0.61 | -0.35 | 0.07 | -8.61 | 2.4E-11 |
| **PTR.R** | 0.629 | 0.613 | -0.43 | -0.55 | -0.30 | 0.07 | -7.59 | 6.8E-09 |
| **RLIC.L** | 0.616 | 0.596 | -0.61 | -0.74 | -0.48 | 0.07 | -10.75 | 2.1E-18 |
| **RLIC.R** | 0.600 | 0.587 | -0.39 | -0.51 | -0.26 | 0.06 | -6.86 | 3.0E-09 |
| **SCR.L** | 0.516 | 0.505 | -0.40 | -0.52 | -0.27 | 0.06 | -7.06 | 4.4E-09 |
| **SCR.R** | 0.511 | 0.500 | -0.40 | -0.52 | -0.27 | 0.06 | -7.02 | 1.3E-08 |
| **SFO.L** | 0.549 | 0.531 | -0.43 | -0.56 | -0.30 | 0.07 | -7.61 | 4.1E-11 |
| **SFO.R** | 0.552 | 0.542 | -0.26 | -0.39 | -0.14 | 0.06 | -4.68 | 4.3E-05 |
| **SLF.L** | 0.529 | 0.510 | -0.67 | -0.80 | -0.55 | 0.07 | -12.01 | 2.3E-23 |
| **SLF.R** | 0.527 | 0.511 | -0.58 | -0.71 | -0.45 | 0.07 | -10.27 | 7.7E-18 |
| **SS.L** | 0.578 | 0.552 | -0.78 | -0.91 | -0.65 | 0.07 | -13.96 | 1.2E-29 |
| **SS.R** | 0.579 | 0.561 | -0.52 | -0.64 | -0.39 | 0.07 | -9.19 | 4.3E-14 |
| **TAP.L** | 0.571 | 0.550 | -0.40 | -0.53 | -0.27 | 0.06 | -7.13 | 2.1E-09 |
| **TAP.R** | 0.603 | 0.586 | -0.33 | -0.46 | -0.21 | 0.06 | -5.94 | 2.0E-06 |
| **UNC.L** | 0.543 | 0.512 | -0.61 | -0.74 | -0.48 | 0.07 | -10.95 | 1.9E-19 |
| **UNC.R** | 0.563 | 0.547 | -0.32 | -0.45 | -0.20 | 0.06 | -5.74 | 8.3E-07 |

**Supplementary Table 7.** Effect sizes for FA differences between healthy controls and the 'R TLE-HS' syndrome.

| ROI | Mean FA controls | Mean FA patients | Cohen's d | 95% CI lower | 95% CI upper | Standard error | Z score | P value |
|---|---|---|---|---|---|---|---|---|
| **AverageFA** | 0.585 | 0.562 | -1.06 | -1.20 | -0.92 | 0.07 | -17.33 | 5.4E-46 |
| **BCC** | 0.690 | 0.659 | -0.87 | -1.01 | -0.73 | 0.07 | -14.22 | 2.9E-34 |
| **GCC** | 0.695 | 0.664 | -0.85 | -0.98 | -0.71 | 0.07 | -13.84 | 2.4E-33 |
| **SCC** | 0.762 | 0.750 | -0.41 | -0.54 | -0.27 | 0.07 | -6.70 | 6.6E-10 |
| **ACR.L** | 0.479 | 0.456 | -0.68 | -0.81 | -0.54 | 0.07 | -11.06 | 3.4E-22 |
| **ACR.R** | 0.484 | 0.458 | -0.78 | -0.91 | -0.64 | 0.07 | -12.71 | 2.0E-27 |
| **ALIC.L** | 0.600 | 0.587 | -0.45 | -0.59 | -0.32 | 0.07 | -7.40 | 1.1E-09 |
| **ALIC.R** | 0.607 | 0.591 | -0.55 | -0.69 | -0.42 | 0.07 | -9.03 | 2.8E-15 |
| **CGC.L** | 0.648 | 0.620 | -0.68 | -0.81 | -0.54 | 0.07 | -11.06 | 3.2E-21 |
| **CGC.R** | 0.604 | 0.574 | -0.75 | -0.89 | -0.61 | 0.07 | -12.23 | 1.2E-25 |
| **CGH.L** | 0.553 | 0.534 | -0.46 | -0.60 | -0.33 | 0.07 | -7.55 | 9.6E-11 |
| **CGH.R** | 0.565 | 0.515 | -1.02 | -1.16 | -0.88 | 0.07 | -16.69 | 8.4E-46 |
| **CST.L** | 0.624 | 0.613 | -0.29 | -0.43 | -0.16 | 0.07 | -4.76 | 9.0E-05 |
| **CST.R** | 0.614 | 0.606 | -0.19 | -0.33 | -0.06 | 0.07 | -3.12 | 0.002 |
| **EC.L** | 0.484 | 0.463 | -0.73 | -0.87 | -0.60 | 0.07 | -11.99 | 1.8E-25 |
| **EC.R** | 0.483 | 0.455 | -0.99 | -1.13 | -0.85 | 0.07 | -16.16 | 2.3E-42 |
| **FX.ST.L** | 0.580 | 0.564 | -0.41 | -0.54 | -0.27 | 0.07 | -6.65 | 4.0E-09 |
| **FX.ST.R** | 0.579 | 0.554 | -0.61 | -0.74 | -0.47 | 0.07 | -9.95 | 1.7E-18 |
| **PCR.L** | 0.506 | 0.493 | -0.37 | -0.51 | -0.24 | 0.07 | -6.13 | 2.2E-08 |
| **PCR.R** | 0.512 | 0.494 | -0.55 | -0.69 | -0.42 | 0.07 | -9.08 | 1.3E-15 |
| **PLIC.L** | 0.697 | 0.693 | -0.14 | -0.27 | 0.00 | 0.07 | -2.27 | 0.066 |
| **PLIC.R** | 0.704 | 0.696 | -0.24 | -0.38 | -0.11 | 0.07 | -4.00 | 2.1E-04 |
| **PTR.L** | 0.625 | 0.608 | -0.41 | -0.55 | -0.28 | 0.07 | -6.76 | 6.5E-10 |
| **PTR.R** | 0.629 | 0.605 | -0.62 | -0.76 | -0.48 | 0.07 | -10.17 | 1.2E-18 |
| **RLIC.L** | 0.616 | 0.604 | -0.31 | -0.45 | -0.18 | 0.07 | -5.15 | 1.9E-06 |
| **RLIC.R** | 0.600 | 0.584 | -0.47 | -0.60 | -0.33 | 0.07 | -7.66 | 2.9E-11 |
| **SCR.L** | 0.516 | 0.501 | -0.53 | -0.67 | -0.40 | 0.07 | -8.71 | 3.2E-13 |
| **SCR.R** | 0.511 | 0.497 | -0.46 | -0.60 | -0.33 | 0.07 | -7.60 | 4.8E-12 |
| **SFO.L** | 0.549 | 0.533 | -0.37 | -0.50 | -0.23 | 0.07 | -6.00 | 6.3E-08 |
| **SFO.R** | 0.552 | 0.534 | -0.47 | -0.60 | -0.33 | 0.07 | -7.64 | 5.0E-11 |
| **SLF.L** | 0.529 | 0.516 | -0.43 | -0.56 | -0.29 | 0.07 | -7.03 | 2.6E-09 |
| **SLF.R** | 0.527 | 0.509 | -0.59 | -0.72 | -0.45 | 0.07 | -9.60 | 2.3E-17 |
| **SS.L** | 0.578 | 0.557 | -0.55 | -0.69 | -0.41 | 0.07 | -8.99 | 1.4E-16 |
| **SS.R** | 0.579 | 0.549 | -0.83 | -0.97 | -0.69 | 0.07 | -13.57 | 2.0E-31 |
| **TAP.L** | 0.571 | 0.535 | -0.68 | -0.82 | -0.54 | 0.07 | -11.10 | 6.7E-22 |
| **TAP.R** | 0.603 | 0.570 | -0.59 | -0.73 | -0.46 | 0.07 | -9.70 | 9.1E-18 |
| **UNC.L** | 0.543 | 0.528 | -0.28 | -0.42 | -0.15 | 0.07 | -4.63 | 3.2E-05 |
| **UNC.R** | 0.563 | 0.515 | -0.90 | -1.04 | -0.76 | 0.07 | -14.69 | 9.3E-37 |

**Supplementary Table 8.** Effect sizes for FA differences between healthy controls and the 'L TLE-NL' syndrome.

| ROI | Mean FA controls | Mean FA patients | Cohen's d | 95% CI lower | 95% CI upper | Standard error | Z score | P value |
|---|---|---|---|---|---|---|---|---|
| **AverageFA** | 0.585 | 0.575 | -0.48 | -0.64 | -0.31 | 0.09 | -6.02 | 1.8E-07 |
| **BCC** | 0.690 | 0.675 | -0.46 | -0.63 | -0.29 | 0.09 | -5.77 | 1.1E-07 |
| **GCC** | 0.695 | 0.683 | -0.39 | -0.56 | -0.23 | 0.09 | -4.96 | 9.8E-06 |
| **SCC** | 0.762 | 0.757 | -0.21 | -0.38 | -0.04 | 0.09 | -2.66 | 0.029 |
| **ACR.L** | 0.479 | 0.470 | -0.32 | -0.48 | -0.15 | 0.09 | -3.98 | 2.0E-04 |
| **ACR.R** | 0.484 | 0.473 | -0.37 | -0.54 | -0.20 | 0.09 | -4.64 | 6.1E-05 |
| **ALIC.L** | 0.600 | 0.590 | -0.35 | -0.52 | -0.19 | 0.09 | -4.46 | 4.4E-05 |
| **ALIC.R** | 0.607 | 0.601 | -0.21 | -0.38 | -0.04 | 0.09 | -2.65 | 0.016 |
| **CGC.L** | 0.648 | 0.638 | -0.25 | -0.41 | -0.08 | 0.09 | -3.12 | 0.004 |
| **CGC.R** | 0.604 | 0.595 | -0.23 | -0.40 | -0.06 | 0.09 | -2.92 | 0.008 |
| **CGH.L** | 0.553 | 0.536 | -0.36 | -0.53 | -0.19 | 0.09 | -4.55 | 1.5E-05 |
| **CGH.R** | 0.565 | 0.555 | -0.23 | -0.40 | -0.06 | 0.09 | -2.91 | 0.012 |
| **CST.L** | 0.624 | 0.621 | -0.07 | -0.24 | 0.10 | 0.09 | -0.87 | 0.378 |
| **CST.R** | 0.614 | 0.614 | 0.00 | -0.17 | 0.17 | 0.09 | 0.00 | 0.910 |
| **EC.L** | 0.484 | 0.471 | -0.47 | -0.63 | -0.30 | 0.09 | -5.88 | 1.1E-07 |
| **EC.R** | 0.483 | 0.472 | -0.41 | -0.58 | -0.24 | 0.09 | -5.14 | 1.5E-06 |
| **FX.ST.L** | 0.580 | 0.571 | -0.27 | -0.44 | -0.10 | 0.09 | -3.43 | 0.002 |
| **FX.ST.R** | 0.579 | 0.573 | -0.15 | -0.32 | 0.01 | 0.09 | -1.94 | 0.050 |
| **PCR.L** | 0.506 | 0.497 | -0.32 | -0.49 | -0.16 | 0.09 | -4.07 | 4.3E-04 |
| **PCR.R** | 0.512 | 0.503 | -0.32 | -0.48 | -0.15 | 0.09 | -4.00 | 1.3E-04 |
| **PLIC.L** | 0.697 | 0.693 | -0.14 | -0.31 | 0.02 | 0.09 | -1.81 | 0.119 |
| **PLIC.R** | 0.704 | 0.700 | -0.15 | -0.31 | 0.02 | 0.09 | -1.84 | 0.079 |
| **PTR.L** | 0.625 | 0.611 | -0.42 | -0.58 | -0.25 | 0.09 | -5.24 | 3.9E-06 |
| **PTR.R** | 0.629 | 0.618 | -0.32 | -0.48 | -0.15 | 0.09 | -3.98 | 2.9E-04 |
| **RLIC.L** | 0.616 | 0.607 | -0.26 | -0.42 | -0.09 | 0.09 | -3.23 | 0.002 |
| **RLIC.R** | 0.600 | 0.590 | -0.29 | -0.45 | -0.12 | 0.09 | -3.63 | 6.1E-04 |
| **SCR.L** | 0.516 | 0.508 | -0.27 | -0.44 | -0.11 | 0.09 | -3.45 | 8.1E-04 |
| **SCR.R** | 0.511 | 0.503 | -0.32 | -0.48 | -0.15 | 0.09 | -3.99 | 7.9E-04 |
| **SFO.L** | 0.549 | 0.542 | -0.18 | -0.34 | -0.01 | 0.09 | -2.22 | 0.045 |
| **SFO.R** | 0.552 | 0.546 | -0.16 | -0.33 | 0.01 | 0.09 | -2.00 | 0.077 |
| **SLF.L** | 0.529 | 0.521 | -0.30 | -0.46 | -0.13 | 0.09 | -3.72 | 4.6E-04 |
| **SLF.R** | 0.527 | 0.518 | -0.32 | -0.49 | -0.16 | 0.09 | -4.07 | 1.5E-04 |
| **SS.L** | 0.578 | 0.563 | -0.46 | -0.63 | -0.30 | 0.09 | -5.84 | 1.0E-07 |
| **SS.R** | 0.579 | 0.565 | -0.42 | -0.59 | -0.26 | 0.09 | -5.35 | 1.1E-06 |
| **TAP.L** | 0.571 | 0.563 | -0.15 | -0.32 | 0.02 | 0.09 | -1.91 | 0.091 |
| **TAP.R** | 0.603 | 0.599 | -0.07 | -0.24 | 0.09 | 0.09 | -0.93 | 0.308 |
| **UNC.L** | 0.543 | 0.533 | -0.19 | -0.35 | -0.02 | 0.09 | -2.37 | 0.019 |
| **UNC.R** | 0.563 | 0.561 | -0.04 | -0.20 | 0.13 | 0.09 | -0.48 | 0.698 |

**Supplementary Table 9.** Effect sizes for FA differences between healthy controls and the 'R TLE-NL' syndrome.

| ROI | Mean FA controls | Mean FA patients | Cohen's d | 95% CI lower | 95% CI upper | Standard error | Z score | P value |
|---|---|---|---|---|---|---|---|---|
| **AverageFA** | 0.585 | 0.573 | -0.58 | -0.78 | -0.39 | 0.10 | -6.14 | 4.8E-08 |
| **BCC** | 0.690 | 0.678 | -0.37 | -0.57 | -0.18 | 0.10 | -3.93 | 1.1E-04 |
| **GCC** | 0.695 | 0.683 | -0.40 | -0.60 | -0.21 | 0.10 | -4.26 | 9.9E-05 |
| **SCC** | 0.762 | 0.755 | -0.31 | -0.50 | -0.11 | 0.10 | -3.22 | 0.005 |
| **ACR.L** | 0.479 | 0.471 | -0.28 | -0.47 | -0.08 | 0.10 | -2.94 | 0.006 |
| **ACR.R** | 0.484 | 0.473 | -0.37 | -0.56 | -0.17 | 0.10 | -3.88 | 4.0E-04 |
| **ALIC.L** | 0.600 | 0.591 | -0.29 | -0.49 | -0.09 | 0.10 | -3.06 | 0.002 |
| **ALIC.R** | 0.607 | 0.600 | -0.22 | -0.41 | -0.02 | 0.10 | -2.28 | 0.024 |
| **CGC.L** | 0.648 | 0.630 | -0.42 | -0.62 | -0.23 | 0.10 | -4.46 | 2.4E-05 |
| **CGC.R** | 0.604 | 0.584 | -0.51 | -0.71 | -0.32 | 0.10 | -5.39 | 5.9E-07 |
| **CGH.L** | 0.553 | 0.537 | -0.33 | -0.52 | -0.13 | 0.10 | -3.45 | 7.1E-04 |
| **CGH.R** | 0.565 | 0.538 | -0.55 | -0.74 | -0.35 | 0.10 | -5.75 | 5.2E-08 |
| **CST.L** | 0.624 | 0.623 | 0.00 | -0.20 | 0.20 | 0.10 | 0.00 | 0.921 |
| **CST.R** | 0.614 | 0.612 | -0.02 | -0.22 | 0.17 | 0.10 | -0.26 | 0.722 |
| **EC.L** | 0.484 | 0.471 | -0.47 | -0.67 | -0.28 | 0.10 | -4.98 | 2.1E-06 |
| **EC.R** | 0.483 | 0.465 | -0.64 | -0.84 | -0.44 | 0.10 | -6.73 | 2.6E-10 |
| **FX.ST.L** | 0.580 | 0.575 | -0.14 | -0.33 | 0.06 | 0.10 | -1.45 | 0.175 |
| **FX.ST.R** | 0.579 | 0.566 | -0.34 | -0.53 | -0.14 | 0.10 | -3.53 | 9.3E-04 |
| **PCR.L** | 0.506 | 0.497 | -0.32 | -0.52 | -0.13 | 0.10 | -3.39 | 0.003 |
| **PCR.R** | 0.512 | 0.505 | -0.25 | -0.45 | -0.05 | 0.10 | -2.63 | 0.012 |
| **PLIC.L** | 0.697 | 0.693 | -0.15 | -0.34 | 0.05 | 0.10 | -1.53 | 0.122 |
| **PLIC.R** | 0.704 | 0.699 | -0.18 | -0.38 | 0.01 | 0.10 | -1.95 | 0.065 |
| **PTR.L** | 0.625 | 0.616 | -0.27 | -0.47 | -0.08 | 0.10 | -2.89 | 0.006 |
| **PTR.R** | 0.629 | 0.618 | -0.35 | -0.54 | -0.15 | 0.10 | -3.64 | 5.5E-04 |
| **RLIC.L** | 0.616 | 0.608 | -0.29 | -0.49 | -0.10 | 0.10 | -3.08 | 0.006 |
| **RLIC.R** | 0.600 | 0.590 | -0.32 | -0.52 | -0.13 | 0.10 | -3.39 | 9.5E-04 |
| **SCR.L** | 0.516 | 0.508 | -0.31 | -0.51 | -0.12 | 0.10 | -3.29 | 0.002 |
| **SCR.R** | 0.511 | 0.503 | -0.36 | -0.55 | -0.16 | 0.10 | -3.77 | 0.001 |
| **SFO.L** | 0.549 | 0.543 | -0.13 | -0.32 | 0.07 | 0.10 | -1.33 | 0.169 |
| **SFO.R** | 0.552 | 0.550 | -0.05 | -0.25 | 0.14 | 0.10 | -0.56 | 0.702 |
| **SLF.L** | 0.529 | 0.520 | -0.34 | -0.53 | -0.14 | 0.10 | -3.54 | 4.5E-04 |
| **SLF.R** | 0.527 | 0.514 | -0.46 | -0.66 | -0.27 | 0.10 | -4.87 | 3.4E-06 |
| **SS.L** | 0.578 | 0.564 | -0.44 | -0.63 | -0.24 | 0.10 | -4.59 | 2.2E-05 |
| **SS.R** | 0.579 | 0.560 | -0.60 | -0.80 | -0.41 | 0.10 | -6.35 | 9.7E-09 |
| **TAP.L** | 0.571 | 0.567 | -0.10 | -0.29 | 0.10 | 0.10 | -1.02 | 0.398 |
| **TAP.R** | 0.603 | 0.593 | -0.21 | -0.40 | -0.01 | 0.10 | -2.17 | 0.035 |
| **UNC.L** | 0.543 | 0.534 | -0.17 | -0.37 | 0.02 | 0.10 | -1.83 | 0.071 |
| **UNC.R** | 0.563 | 0.536 | -0.50 | -0.70 | -0.30 | 0.10 | -5.28 | 8.5E-07 |

**Supplementary Table 10.** Effect sizes for FA differences between healthy controls and the 'GGE' syndrome.

| ROI | Mean FA controls | Mean FA patients | Cohen's d | 95% CI lower | 95% CI upper | Standard error | Z score | P value |
|---|---|---|---|---|---|---|---|---|
| **AverageFA** | 0.585 | 0.575 | -0.49 | -0.65 | -0.33 | 0.08 | -6.59 | 4.5E-10 |
| **BCC** | 0.690 | 0.677 | -0.44 | -0.60 | -0.28 | 0.08 | -5.91 | 1.8E-08 |
| **GCC** | 0.695 | 0.681 | -0.57 | -0.73 | -0.41 | 0.08 | -7.59 | 1.1E-11 |
| **SCC** | 0.762 | 0.753 | -0.39 | -0.55 | -0.23 | 0.08 | -5.27 | 1.1E-06 |
| **ACR.L** | 0.479 | 0.471 | -0.35 | -0.51 | -0.19 | 0.08 | -4.73 | 3.9E-06 |
| **ACR.R** | 0.484 | 0.474 | -0.40 | -0.56 | -0.24 | 0.08 | -5.36 | 6.3E-07 |
| **ALIC.L** | 0.600 | 0.589 | -0.44 | -0.60 | -0.28 | 0.08 | -5.86 | 5.7E-07 |
| **ALIC.R** | 0.607 | 0.601 | -0.21 | -0.37 | -0.06 | 0.08 | -2.87 | .0059 |
| **CGC.L** | 0.648 | 0.631 | -0.46 | -0.62 | -0.30 | 0.08 | -6.20 | 4.6E-08 |
| **CGC.R** | 0.604 | 0.589 | -0.42 | -0.58 | -0.26 | 0.08 | -5.64 | 3.8E-07 |
| **CGH.L** | 0.553 | 0.532 | -0.41 | -0.57 | -0.25 | 0.08 | -5.55 | 2.5E-07 |
| **CGH.R** | 0.565 | 0.549 | -0.30 | -0.46 | -0.14 | 0.08 | -4.06 | 2.1E-04 |
| **CST.L** | 0.624 | 0.622 | -0.02 | -0.18 | 0.14 | 0.08 | -0.30 | 0.729 |
| **CST.R** | 0.614 | 0.614 | 0.00 | -0.16 | 0.16 | 0.08 | 0.00 | 0.935 |
| **EC.L** | 0.484 | 0.470 | -0.55 | -0.71 | -0.39 | 0.08 | -7.43 | 3.7E-12 |
| **EC.R** | 0.483 | 0.473 | -0.42 | -0.58 | -0.26 | 0.08 | -5.62 | 7.7E-07 |
| **FX.ST.L** | 0.580 | 0.568 | -0.36 | -0.52 | -0.20 | 0.08 | -4.88 | 7.3E-06 |
| **FX.ST.R** | 0.579 | 0.567 | -0.37 | -0.53 | -0.21 | 0.08 | -4.96 | 7.5E-06 |
| **PCR.L** | 0.506 | 0.493 | -0.50 | -0.66 | -0.34 | 0.08 | -6.76 | 4.1E-10 |
| **PCR.R** | 0.512 | 0.502 | -0.39 | -0.55 | -0.23 | 0.08 | -5.29 | 3.8E-07 |
| **PLIC.L** | 0.697 | 0.691 | -0.22 | -0.38 | -0.06 | 0.08 | -2.92 | 0.003 |
| **PLIC.R** | 0.704 | 0.702 | -0.07 | -0.23 | 0.08 | 0.08 | -0.99 | 0.195 |
| **PTR.L** | 0.625 | 0.616 | -0.33 | -0.49 | -0.18 | 0.08 | -4.49 | 1.7E-05 |
| **PTR.R** | 0.629 | 0.621 | -0.34 | -0.50 | -0.18 | 0.08 | -4.61 | 8.0E-05 |
| **RLIC.L** | 0.616 | 0.601 | -0.51 | -0.67 | -0.35 | 0.08 | -6.88 | 2.3E-10 |
| **RLIC.R** | 0.600 | 0.590 | -0.36 | -0.51 | -0.20 | 0.08 | -4.77 | 5.8E-06 |
| **SCR.L** | 0.516 | 0.506 | -0.44 | -0.60 | -0.28 | 0.08 | -5.90 | 1.4E-08 |
| **SCR.R** | 0.511 | 0.505 | -0.28 | -0.44 | -0.12 | 0.08 | -3.73 | 3.2E-04 |
| **SFO.L** | 0.549 | 0.537 | -0.35 | -0.51 | -0.20 | 0.08 | -4.76 | 1.8E-05 |
| **SFO.R** | 0.552 | 0.549 | -0.08 | -0.24 | 0.08 | 0.08 | -1.07 | 0.332 |
| **SLF.L** | 0.529 | 0.516 | -0.57 | -0.73 | -0.41 | 0.08 | -7.60 | 5.0E-12 |
| **SLF.R** | 0.527 | 0.514 | -0.51 | -0.67 | -0.35 | 0.08 | -6.83 | 4.8E-11 |
| **SS.L** | 0.578 | 0.566 | -0.41 | -0.57 | -0.25 | 0.08 | -5.51 | 9.9E-07 |
| **SS.R** | 0.579 | 0.569 | -0.37 | -0.53 | -0.21 | 0.08 | -4.93 | 2.1E-05 |
| **TAP.L** | 0.571 | 0.562 | -0.15 | -0.31 | 0.00 | 0.08 | -2.06 | 0.057 |
| **TAP.R** | 0.603 | 0.595 | -0.19 | -0.34 | -0.03 | 0.08 | -2.51 | 0.033 |
| **UNC.L** | 0.543 | 0.532 | -0.21 | -0.37 | -0.05 | 0.08 | -2.86 | 0.008 |
| **UNC.R** | 0.563 | 0.555 | -0.15 | -0.31 | 0.01 | 0.08 | -2.05 | 0.067 |

**Supplementary Table 11.** Effect sizes for FA differences between healthy controls and the 'ExE' syndrome.

| ROI | Mean FA controls | Mean FA patients | Cohen's d | 95% CI lower | 95% CI upper | Standard error | Z score | P value |
|---|---|---|---|---|---|---|---|---|
| **AverageFA** | 0.585 | 0.570 | -0.75 | -0.91 | -0.60 | 0.08 | -10.41 | 1.1E-19 |
| **BCC** | 0.690 | 0.667 | -0.73 | -0.89 | -0.57 | 0.08 | -10.07 | 9.4E-20 |
| **GCC** | 0.695 | 0.677 | -0.64 | -0.80 | -0.49 | 0.08 | -8.89 | 9.5E-16 |
| **SCC** | 0.762 | 0.752 | -0.42 | -0.57 | -0.26 | 0.08 | -5.75 | 4.2E-07 |
| **ACR.L** | 0.479 | 0.464 | -0.55 | -0.71 | -0.40 | 0.08 | -7.64 | 9.8E-13 |
| **ACR.R** | 0.484 | 0.467 | -0.63 | -0.78 | -0.47 | 0.08 | -8.66 | 2.8E-14 |
| **ALIC.L** | 0.600 | 0.584 | -0.57 | -0.73 | -0.42 | 0.08 | -7.89 | 8.6E-12 |
| **ALIC.R** | 0.607 | 0.593 | -0.49 | -0.65 | -0.34 | 0.08 | -6.81 | 2.1E-09 |
| **CGC.L** | 0.648 | 0.628 | -0.49 | -0.65 | -0.34 | 0.08 | -6.82 | 8.2E-10 |
| **CGC.R** | 0.604 | 0.585 | -0.48 | -0.64 | -0.32 | 0.08 | -6.62 | 4.1E-09 |
| **CGH.L** | 0.553 | 0.532 | -0.41 | -0.56 | -0.25 | 0.08 | -5.60 | 3.7E-07 |
| **CGH.R** | 0.565 | 0.546 | -0.36 | -0.52 | -0.21 | 0.08 | -4.99 | 6.6E-06 |
| **CST.L** | 0.624 | 0.619 | -0.11 | -0.27 | 0.04 | 0.08 | -1.58 | 0.140 |
| **CST.R** | 0.614 | 0.608 | -0.15 | -0.30 | 0.01 | 0.08 | -2.05 | 0.072 |
| **EC.L** | 0.484 | 0.468 | -0.60 | -0.76 | -0.44 | 0.08 | -8.30 | 5.4E-14 |
| **EC.R** | 0.483 | 0.467 | -0.58 | -0.74 | -0.43 | 0.08 | -8.04 | 2.0E-13 |
| **FX.ST.L** | 0.580 | 0.568 | -0.32 | -0.48 | -0.17 | 0.08 | -4.46 | 3.1E-05 |
| **FX.ST.R** | 0.579 | 0.567 | -0.33 | -0.49 | -0.18 | 0.08 | -4.58 | 4.4E-05 |
| **PCR.L** | 0.506 | 0.497 | -0.35 | -0.50 | -0.19 | 0.08 | -4.78 | 1.5E-05 |
| **PCR.R** | 0.512 | 0.502 | -0.39 | -0.54 | -0.23 | 0.08 | -5.34 | 2.6E-07 |
| **PLIC.L** | 0.697 | 0.692 | -0.21 | -0.37 | -0.06 | 0.08 | -2.96 | 0.009 |
| **PLIC.R** | 0.704 | 0.699 | -0.18 | -0.33 | -0.03 | 0.08 | -2.48 | 0.009 |
| **PTR.L** | 0.625 | 0.613 | -0.41 | -0.56 | -0.25 | 0.08 | -5.64 | 3.8E-07 |
| **PTR.R** | 0.629 | 0.617 | -0.42 | -0.58 | -0.27 | 0.08 | -5.84 | 4.1E-07 |
| **RLIC.L** | 0.616 | 0.608 | -0.28 | -0.44 | -0.13 | 0.08 | -3.92 | 1.6E-04 |
| **RLIC.R** | 0.600 | 0.593 | -0.26 | -0.41 | -0.10 | 0.08 | -3.53 | 7.3E-04 |
| **SCR.L** | 0.516 | 0.506 | -0.47 | -0.63 | -0.31 | 0.08 | -6.48 | 4.2E-09 |
| **SCR.R** | 0.511 | 0.502 | -0.39 | -0.55 | -0.24 | 0.08 | -5.44 | 2.6E-07 |
| **SFO.L** | 0.549 | 0.532 | -0.42 | -0.58 | -0.26 | 0.08 | -5.79 | 4.5E-08 |
| **SFO.R** | 0.552 | 0.538 | -0.36 | -0.51 | -0.20 | 0.08 | -4.91 | 9.6E-06 |
| **SLF.L** | 0.529 | 0.518 | -0.43 | -0.59 | -0.28 | 0.08 | -5.95 | 6.5E-08 |
| **SLF.R** | 0.527 | 0.514 | -0.49 | -0.65 | -0.33 | 0.08 | -6.76 | 4.1E-10 |
| **SS.L** | 0.578 | 0.565 | -0.43 | -0.58 | -0.27 | 0.08 | -5.90 | 1.4E-07 |
| **SS.R** | 0.579 | 0.563 | -0.51 | -0.66 | -0.35 | 0.08 | -7.02 | 5.8E-10 |
| **TAP.L** | 0.571 | 0.554 | -0.32 | -0.48 | -0.17 | 0.08 | -4.47 | 9.8E-05 |
| **TAP.R** | 0.603 | 0.590 | -0.24 | -0.40 | -0.09 | 0.08 | -3.33 | 0.002 |
| **UNC.L** | 0.543 | 0.523 | -0.38 | -0.54 | -0.23 | 0.08 | -5.26 | 1.6E-06 |
| **UNC.R** | 0.563 | 0.543 | -0.37 | -0.52 | -0.21 | 0.08 | -5.05 | 3.6E-06 |

**Supplementary Table 12.** Effect sizes for MD differences between healthy controls and the 'All Epilepsies' syndrome.

| ROI | Mean MD controls | Mean MD patients | Cohen's d | 95% CI lower | 95% CI upper | Standard error | Z score | P value |
|---|---|---|---|---|---|---|---|---|
| **AverageMD** | 0.000801 | 0.000819 | 0.37 | 0.28 | 0.46 | 0.05 | 11.92 | 3.3E-16 |
| **BCC** | 0.000881 | 0.000903 | 0.23 | 0.14 | 0.32 | 0.05 | 7.44 | 3.2E-07 |
| **GCC** | 0.000816 | 0.000834 | 0.14 | 0.05 | 0.23 | 0.04 | 4.57 | 0.002 |
| **SCC** | 0.000791 | 0.000795 | 0.06 | -0.03 | 0.14 | 0.04 | 1.81 | 0.210 |
| **ACR.L** | 0.000745 | 0.000763 | 0.32 | 0.23 | 0.41 | 0.05 | 10.20 | 2.5E-12 |
| **ACR.R** | 0.000739 | 0.000757 | 0.34 | 0.26 | 0.43 | 0.05 | 11.11 | 2.7E-14 |
| **ALIC.L** | 0.000695 | 0.000702 | 0.14 | 0.05 | 0.23 | 0.04 | 4.41 | 0.002 |
| **ALIC.R** | 0.000695 | 0.000702 | 0.15 | 0.06 | 0.24 | 0.05 | 4.87 | 8.1E-04 |
| **CGC.L** | 0.000738 | 0.000748 | 0.17 | 0.09 | 0.26 | 0.05 | 5.62 | 1.1E-04 |
| **CGC.R** | 0.000729 | 0.000740 | 0.18 | 0.09 | 0.27 | 0.05 | 5.90 | 4.6E-05 |
| **CGH.L** | 0.000876 | 0.000896 | 0.17 | 0.08 | 0.26 | 0.05 | 5.47 | 1.7E-04 |
| **CGH.R** | 0.000859 | 0.000877 | 0.15 | 0.07 | 0.24 | 0.05 | 4.96 | 6.4E-04 |
| **CST.L** | 0.000740 | 0.000741 | 0.02 | -0.07 | 0.11 | 0.04 | 0.62 | 0.670 |
| **CST.R** | 0.000743 | 0.000743 | 0.01 | -0.08 | 0.10 | 0.04 | 0.40 | 0.780 |
| **EC.L** | 0.000741 | 0.000755 | 0.34 | 0.25 | 0.43 | 0.05 | 11.05 | 4.2E-14 |
| **EC.R** | 0.000737 | 0.000752 | 0.38 | 0.29 | 0.47 | 0.05 | 12.15 | 1.1E-16 |
| **FX.ST.L** | 0.000787 | 0.000797 | 0.14 | 0.05 | 0.22 | 0.04 | 4.39 | 0.002 |
| **FX.ST.R** | 0.000781 | 0.000794 | 0.19 | 0.10 | 0.28 | 0.05 | 6.03 | 3.3E-05 |
| **PCR.L** | 0.000789 | 0.000805 | 0.28 | 0.20 | 0.37 | 0.05 | 9.17 | 2.9E-10 |
| **PCR.R** | 0.000802 | 0.000819 | 0.27 | 0.18 | 0.36 | 0.05 | 8.69 | 2.6E-09 |
| **PLIC.L** | 0.000683 | 0.000686 | 0.12 | 0.03 | 0.20 | 0.04 | 3.73 | 0.010 |
| **PLIC.R** | 0.000677 | 0.000683 | 0.15 | 0.06 | 0.24 | 0.05 | 4.91 | 7.2E-04 |
| **PTR.L** | 0.000820 | 0.000833 | 0.17 | 0.08 | 0.26 | 0.05 | 5.41 | 2.1E-04 |
| **PTR.R** | 0.000811 | 0.000825 | 0.17 | 0.08 | 0.26 | 0.05 | 5.43 | 1.8E-04 |
| **RLIC.L** | 0.000778 | 0.000786 | 0.16 | 0.07 | 0.25 | 0.05 | 5.25 | 3.0E-04 |
| **RLIC.R** | 0.000776 | 0.000784 | 0.13 | 0.04 | 0.22 | 0.04 | 4.20 | 0.004 |
| **SCR.L** | 0.000697 | 0.000708 | 0.27 | 0.18 | 0.36 | 0.05 | 8.82 | 1.5E-09 |
| **SCR.R** | 0.000696 | 0.000706 | 0.23 | 0.14 | 0.32 | 0.05 | 7.37 | 3.7E-07 |
| **SFO.L** | 0.000679 | 0.000690 | 0.11 | 0.02 | 0.20 | 0.04 | 3.60 | 0.013 |
| **SFO.R** | 0.000669 | 0.000684 | 0.18 | 0.09 | 0.27 | 0.05 | 5.77 | 7.3E-05 |
| **SLF.L** | 0.000709 | 0.000719 | 0.29 | 0.20 | 0.38 | 0.05 | 9.46 | 9.8E-11 |
| **SLF.R** | 0.000705 | 0.000717 | 0.36 | 0.27 | 0.45 | 0.05 | 11.66 | 1.7E-15 |
| **SS.L** | 0.000787 | 0.000806 | 0.31 | 0.22 | 0.40 | 0.05 | 10.12 | 4.4E-12 |
| **SS.R** | 0.000773 | 0.000791 | 0.32 | 0.23 | 0.41 | 0.05 | 10.30 | 1.5E-12 |
| **TAP.L** | 0.000936 | 0.000960 | 0.14 | 0.05 | 0.23 | 0.04 | 4.54 | 0.002 |
| **TAP.R** | 0.000922 | 0.000943 | 0.13 | 0.04 | 0.22 | 0.04 | 4.27 | 0.003 |
| **UNC.L** | 0.000761 | 0.000781 | 0.22 | 0.13 | 0.31 | 0.05 | 7.16 | 8.0E-07 |
| **UNC.R** | 0.000740 | 0.000765 | 0.29 | 0.20 | 0.38 | 0.05 | 9.36 | 1.3E-10 |

**Supplementary Table 13.** Effect sizes for MD differences between healthy controls and the 'L TLE-HS' syndrome.

| ROI | Mean MD controls | Mean MD patients | Cohen's d | 95% CI lower | 95% CI upper | Standard error | Z score | P value |
|---|---|---|---|---|---|---|---|---|
| **AverageMD** | 0.000801 | 0.000828 | 0.55 | 0.41 | 0.69 | 0.07 | 9.15 | 3.4E-15 |
| **BCC** | 0.000881 | 0.000918 | 0.37 | 0.23 | 0.50 | 0.07 | 6.09 | 1.2E-07 |
| **GCC** | 0.000816 | 0.000856 | 0.28 | 0.15 | 0.42 | 0.07 | 4.73 | 4.0E-05 |
| **SCC** | 0.000791 | 0.000799 | 0.11 | -0.03 | 0.24 | 0.07 | 1.77 | 0.123 |
| **ACR.L** | 0.000745 | 0.000773 | 0.49 | 0.36 | 0.63 | 0.07 | 8.21 | 1.3E-12 |
| **ACR.R** | 0.000739 | 0.000760 | 0.39 | 0.26 | 0.53 | 0.07 | 6.54 | 1.4E-08 |
| **ALIC.L** | 0.000695 | 0.000712 | 0.33 | 0.20 | 0.47 | 0.07 | 5.56 | 1.4E-06 |
| **ALIC.R** | 0.000695 | 0.000703 | 0.21 | 0.08 | 0.35 | 0.07 | 3.57 | 0.002 |
| **CGC.L** | 0.000738 | 0.000761 | 0.39 | 0.25 | 0.53 | 0.07 | 6.49 | 2.0E-08 |
| **CGC.R** | 0.000729 | 0.000746 | 0.27 | 0.14 | 0.41 | 0.07 | 4.51 | 8.7E-05 |
| **CGH.L** | 0.000876 | 0.000928 | 0.48 | 0.34 | 0.61 | 0.07 | 7.95 | 6.1E-12 |
| **CGH.R** | 0.000859 | 0.000876 | 0.20 | 0.07 | 0.33 | 0.07 | 3.32 | 0.004 |
| **CST.L** | 0.000740 | 0.000737 | 0.01 | -0.13 | 0.14 | 0.07 | 0.09 | 0.936 |
| **CST.R** | 0.000743 | 0.000743 | 0.05 | -0.08 | 0.19 | 0.07 | 0.86 | 0.454 |
| **EC.L** | 0.000741 | 0.000769 | 0.69 | 0.55 | 0.83 | 0.07 | 11.45 | 1.3E-22 |
| **EC.R** | 0.000737 | 0.000755 | 0.48 | 0.34 | 0.61 | 0.07 | 7.94 | 6.3E-12 |
| **FX.ST.L** | 0.000787 | 0.000817 | 0.42 | 0.28 | 0.55 | 0.07 | 6.96 | 1.8E-09 |
| **FX.ST.R** | 0.000781 | 0.000796 | 0.25 | 0.11 | 0.38 | 0.07 | 4.11 | 3.5E-04 |
| **PCR.L** | 0.000789 | 0.000814 | 0.44 | 0.30 | 0.57 | 0.07 | 7.30 | 2.7E-10 |
| **PCR.R** | 0.000802 | 0.000819 | 0.27 | 0.14 | 0.41 | 0.07 | 4.54 | 8.2E-05 |
| **PLIC.L** | 0.000683 | 0.000690 | 0.27 | 0.13 | 0.40 | 0.07 | 4.43 | 1.2E-04 |
| **PLIC.R** | 0.000677 | 0.000684 | 0.26 | 0.13 | 0.40 | 0.07 | 4.35 | 1.5E-04 |
| **PTR.L** | 0.000820 | 0.000843 | 0.26 | 0.13 | 0.39 | 0.07 | 4.32 | 1.7E-04 |
| **PTR.R** | 0.000811 | 0.000829 | 0.20 | 0.06 | 0.33 | 0.07 | 3.27 | 0.004 |
| **RLIC.L** | 0.000778 | 0.000797 | 0.37 | 0.24 | 0.51 | 0.07 | 6.20 | 7.1E-08 |
| **RLIC.R** | 0.000776 | 0.000788 | 0.21 | 0.07 | 0.34 | 0.07 | 3.48 | 0.002 |
| **SCR.L** | 0.000697 | 0.000714 | 0.42 | 0.28 | 0.55 | 0.07 | 6.96 | 1.5E-09 |
| **SCR.R** | 0.000696 | 0.000708 | 0.27 | 0.14 | 0.41 | 0.07 | 4.55 | 7.4E-05 |
| **SFO.L** | 0.000679 | 0.000701 | 0.22 | 0.08 | 0.35 | 0.07 | 3.61 | 0.002 |
| **SFO.R** | 0.000669 | 0.000689 | 0.24 | 0.11 | 0.38 | 0.07 | 4.04 | 4.5E-04 |
| **SLF.L** | 0.000709 | 0.000724 | 0.44 | 0.30 | 0.57 | 0.07 | 7.27 | 3.3E-10 |
| **SLF.R** | 0.000705 | 0.000719 | 0.40 | 0.26 | 0.53 | 0.07 | 6.61 | 1.1E-08 |
| **SS.L** | 0.000787 | 0.000827 | 0.66 | 0.52 | 0.80 | 0.07 | 10.97 | 5.1E-21 |
| **SS.R** | 0.000773 | 0.000789 | 0.31 | 0.17 | 0.44 | 0.07 | 5.09 | 1.0E-05 |
| **TAP.L** | 0.000936 | 0.000976 | 0.21 | 0.07 | 0.34 | 0.07 | 3.42 | 0.003 |
| **TAP.R** | 0.000922 | 0.000943 | 0.11 | -0.03 | 0.24 | 0.07 | 1.79 | 0.119 |
| **UNC.L** | 0.000761 | 0.000801 | 0.47 | 0.34 | 0.61 | 0.07 | 7.86 | 1.1E-11 |
| **UNC.R** | 0.000740 | 0.000763 | 0.30 | 0.17 | 0.44 | 0.07 | 5.01 | 1.3E-05 |

**Supplementary Table 14.** Effect sizes for MD differences between healthy controls and the 'R TLE-HS' syndrome.

| ROI | Mean MD controls | Mean MD patients | Cohen's d | 95% CI lower | 95% CI upper | Standard error | Z score | P value |
|---|---|---|---|---|---|---|---|---|
| **AverageMD** | 0.000801 | 0.000831 | 0.60 | 0.45 | 0.75 | 0.08 | 8.93 | 3.8E-15 |
| **BCC** | 0.000881 | 0.000927 | 0.46 | 0.31 | 0.61 | 0.08 | 6.79 | 1.9E-09 |
| **GCC** | 0.000816 | 0.000866 | 0.32 | 0.17 | 0.47 | 0.08 | 4.73 | 2.7E-05 |
| **SCC** | 0.000791 | 0.000808 | 0.20 | 0.06 | 0.35 | 0.07 | 3.00 | 0.007 |
| **ACR.L** | 0.000745 | 0.000771 | 0.44 | 0.29 | 0.59 | 0.08 | 6.56 | 6.7E-09 |
| **ACR.R** | 0.000739 | 0.000771 | 0.59 | 0.44 | 0.74 | 0.08 | 8.74 | 1.5E-14 |
| **ALIC.L** | 0.000695 | 0.000701 | 0.17 | 0.02 | 0.32 | 0.07 | 2.53 | 0.024 |
| **ALIC.R** | 0.000695 | 0.000707 | 0.30 | 0.15 | 0.45 | 0.08 | 4.47 | 7.5E-05 |
| **CGC.L** | 0.000738 | 0.000747 | 0.18 | 0.03 | 0.33 | 0.07 | 2.70 | 0.017 |
| **CGC.R** | 0.000729 | 0.000751 | 0.34 | 0.19 | 0.49 | 0.08 | 5.08 | 6.6E-06 |
| **CGH.L** | 0.000876 | 0.000880 | 0.13 | -0.02 | 0.28 | 0.07 | 1.92 | 0.087 |
| **CGH.R** | 0.000859 | 0.000895 | 0.37 | 0.22 | 0.51 | 0.08 | 5.44 | 1.4E-06 |
| **CST.L** | 0.000740 | 0.000739 | 0.03 | -0.12 | 0.17 | 0.07 | 0.38 | 0.737 |
| **CST.R** | 0.000743 | 0.000738 | 0.01 | -0.14 | 0.15 | 0.07 | 0.09 | 0.937 |
| **EC.L** | 0.000741 | 0.000756 | 0.41 | 0.26 | 0.55 | 0.08 | 6.02 | 9.0E-08 |
| **EC.R** | 0.000737 | 0.000761 | 0.62 | 0.47 | 0.77 | 0.08 | 9.20 | 6.3E-16 |
| **FX.ST.L** | 0.000787 | 0.000785 | 0.02 | -0.12 | 0.17 | 0.07 | 0.36 | 0.748 |
| **FX.ST.R** | 0.000781 | 0.000810 | 0.43 | 0.28 | 0.58 | 0.08 | 6.36 | 1.8E-08 |
| **PCR.L** | 0.000789 | 0.000805 | 0.30 | 0.15 | 0.44 | 0.08 | 4.39 | 9.1E-05 |
| **PCR.R** | 0.000802 | 0.000830 | 0.44 | 0.29 | 0.58 | 0.08 | 6.48 | 1.0E-08 |
| **PLIC.L** | 0.000683 | 0.000686 | 0.19 | 0.04 | 0.33 | 0.07 | 2.76 | 0.014 |
| **PLIC.R** | 0.000677 | 0.000684 | 0.27 | 0.13 | 0.42 | 0.08 | 4.08 | 3.0E-04 |
| **PTR.L** | 0.000820 | 0.000831 | 0.12 | -0.02 | 0.27 | 0.07 | 1.81 | 0.107 |
| **PTR.R** | 0.000811 | 0.000834 | 0.24 | 0.09 | 0.39 | 0.08 | 3.55 | 0.002 |
| **RLIC.L** | 0.000778 | 0.000780 | 0.08 | -0.07 | 0.22 | 0.07 | 1.16 | 0.302 |
| **RLIC.R** | 0.000776 | 0.000789 | 0.22 | 0.07 | 0.37 | 0.07 | 3.25 | 0.004 |
| **SCR.L** | 0.000697 | 0.000709 | 0.32 | 0.17 | 0.46 | 0.08 | 4.71 | 3.1E-05 |
| **SCR.R** | 0.000696 | 0.000711 | 0.34 | 0.20 | 0.49 | 0.08 | 5.11 | 6.1E-06 |
| **SFO.L** | 0.000679 | 0.000694 | 0.16 | 0.01 | 0.31 | 0.07 | 2.35 | 0.036 |
| **SFO.R** | 0.000669 | 0.000688 | 0.23 | 0.08 | 0.37 | 0.07 | 3.36 | 0.003 |
| **SLF.L** | 0.000709 | 0.000718 | 0.29 | 0.14 | 0.44 | 0.08 | 4.29 | 1.5E-04 |
| **SLF.R** | 0.000705 | 0.000721 | 0.46 | 0.31 | 0.60 | 0.08 | 6.79 | 1.9E-09 |
| **SS.L** | 0.000787 | 0.000802 | 0.27 | 0.12 | 0.42 | 0.08 | 4.03 | 3.5E-04 |
| **SS.R** | 0.000773 | 0.000811 | 0.67 | 0.52 | 0.82 | 0.08 | 9.93 | 3.1E-18 |
| **TAP.L** | 0.000936 | 0.000984 | 0.25 | 0.11 | 0.40 | 0.08 | 3.78 | 7.9E-04 |
| **TAP.R** | 0.000922 | 0.000977 | 0.30 | 0.15 | 0.44 | 0.08 | 4.42 | 8.7E-05 |
| **UNC.L** | 0.000761 | 0.000767 | 0.10 | -0.04 | 0.25 | 0.07 | 1.54 | 0.170 |
| **UNC.R** | 0.000740 | 0.000788 | 0.62 | 0.47 | 0.76 | 0.08 | 9.14 | 8.8E-16 |

**Supplementary Table 15.** Effect sizes for MD differences between healthy controls and the 'L TLE-NL' syndrome.

| ROI | Mean MD controls | Mean MD patients | Cohen's d | 95% CI lower | 95% CI upper | Standard error | Z score | P value |
|---|---|---|---|---|---|---|---|---|
| **AverageMD** | 0.000801 | 0.000810 | 0.20 | 0.04 | 0.37 | 0.08 | 2.60 | 0.020 |
| **BCC** | 0.000881 | 0.000879 | 0.00 | -0.17 | 0.16 | 0.08 | -0.04 | 1.000 |
| **GCC** | 0.000816 | 0.000795 | -0.14 | -0.31 | 0.02 | 0.08 | -1.83 | 0.101 |
| **SCC** | 0.000791 | 0.000777 | -0.15 | -0.31 | 0.02 | 0.08 | -1.90 | 0.088 |
| **ACR.L** | 0.000745 | 0.000750 | 0.10 | -0.06 | 0.27 | 0.08 | 1.35 | 0.227 |
| **ACR.R** | 0.000739 | 0.000747 | 0.17 | 0.01 | 0.34 | 0.08 | 2.24 | 0.044 |
| **ALIC.L** | 0.000695 | 0.000697 | 0.05 | -0.11 | 0.22 | 0.08 | 0.70 | 0.530 |
| **ALIC.R** | 0.000695 | 0.000698 | 0.08 | -0.09 | 0.25 | 0.08 | 1.04 | 0.351 |
| **CGC.L** | 0.000738 | 0.000741 | 0.06 | -0.10 | 0.23 | 0.08 | 0.82 | 0.463 |
| **CGC.R** | 0.000729 | 0.000731 | 0.05 | -0.12 | 0.21 | 0.08 | 0.62 | 0.578 |
| **CGH.L** | 0.000876 | 0.000874 | 0.00 | -0.17 | 0.16 | 0.08 | -0.06 | 0.958 |
| **CGH.R** | 0.000859 | 0.000856 | -0.02 | -0.19 | 0.15 | 0.08 | -0.26 | 0.816 |
| **CST.L** | 0.000740 | 0.000752 | 0.11 | -0.06 | 0.27 | 0.08 | 1.37 | 0.220 |
| **CST.R** | 0.000743 | 0.000752 | 0.09 | -0.07 | 0.26 | 0.08 | 1.19 | 0.287 |
| **EC.L** | 0.000741 | 0.000750 | 0.24 | 0.08 | 0.41 | 0.08 | 3.13 | 0.005 |
| **EC.R** | 0.000737 | 0.000747 | 0.26 | 0.10 | 0.43 | 0.08 | 3.38 | 0.002 |
| **FX.ST.L** | 0.000787 | 0.000787 | 0.02 | -0.14 | 0.19 | 0.08 | 0.31 | 0.780 |
| **FX.ST.R** | 0.000781 | 0.000776 | -0.05 | -0.21 | 0.12 | 0.08 | -0.61 | 0.586 |
| **PCR.L** | 0.000789 | 0.000798 | 0.18 | 0.01 | 0.34 | 0.08 | 2.25 | 0.044 |
| **PCR.R** | 0.000802 | 0.000813 | 0.20 | 0.04 | 0.37 | 0.08 | 2.61 | 0.020 |
| **PLIC.L** | 0.000683 | 0.000686 | 0.10 | -0.07 | 0.27 | 0.08 | 1.29 | 0.246 |
| **PLIC.R** | 0.000677 | 0.000683 | 0.18 | 0.01 | 0.34 | 0.08 | 2.26 | 0.043 |
| **PTR.L** | 0.000820 | 0.000832 | 0.16 | -0.01 | 0.33 | 0.08 | 2.06 | 0.065 |
| **PTR.R** | 0.000811 | 0.000820 | 0.11 | -0.05 | 0.28 | 0.08 | 1.45 | 0.192 |
| **RLIC.L** | 0.000778 | 0.000785 | 0.14 | -0.03 | 0.30 | 0.08 | 1.75 | 0.117 |
| **RLIC.R** | 0.000776 | 0.000780 | 0.08 | -0.08 | 0.25 | 0.08 | 1.04 | 0.349 |
| **SCR.L** | 0.000697 | 0.000705 | 0.22 | 0.05 | 0.38 | 0.08 | 2.77 | 0.013 |
| **SCR.R** | 0.000696 | 0.000704 | 0.21 | 0.04 | 0.37 | 0.08 | 2.63 | 0.018 |
| **SFO.L** | 0.000679 | 0.000676 | -0.01 | -0.17 | 0.16 | 0.08 | -0.09 | 0.938 |
| **SFO.R** | 0.000669 | 0.000678 | 0.11 | -0.05 | 0.28 | 0.08 | 1.47 | 0.188 |
| **SLF.L** | 0.000709 | 0.000717 | 0.25 | 0.08 | 0.42 | 0.08 | 3.21 | 0.004 |
| **SLF.R** | 0.000705 | 0.000716 | 0.35 | 0.18 | 0.51 | 0.09 | 4.46 | 7.1E-05 |
| **SS.L** | 0.000787 | 0.000799 | 0.22 | 0.05 | 0.39 | 0.08 | 2.83 | 0.011 |
| **SS.R** | 0.000773 | 0.000785 | 0.24 | 0.08 | 0.41 | 0.08 | 3.10 | 0.005 |
| **TAP.L** | 0.000936 | 0.000938 | 0.02 | -0.15 | 0.19 | 0.08 | 0.25 | 0.823 |
| **TAP.R** | 0.000922 | 0.000927 | 0.04 | -0.13 | 0.20 | 0.08 | 0.48 | 0.666 |
| **UNC.L** | 0.000761 | 0.000779 | 0.21 | 0.04 | 0.37 | 0.08 | 2.67 | 0.017 |
| **UNC.R** | 0.000740 | 0.000752 | 0.14 | -0.03 | 0.31 | 0.08 | 1.80 | 0.107 |

**Supplementary Table 16.** Effect sizes for MD differences between healthy controls and the 'R TLE-NL' syndrome.

| ROI | Mean MD controls | Mean MD patients | Cohen's d | 95% CI lower | 95% CI upper | Standard error | Z score | P value |
|---|---|---|---|---|---|---|---|---|
| **AverageMD** | 0.000801 | 0.000810 | 0.21 | 0.01 | 0.42 | 0.10 | 2.19 | 0.039 |
| **BCC** | 0.000881 | 0.000890 | 0.13 | -0.07 | 0.33 | 0.10 | 1.32 | 0.212 |
| **GCC** | 0.000816 | 0.000817 | 0.05 | -0.15 | 0.26 | 0.10 | 0.55 | 0.604 |
| **SCC** | 0.000791 | 0.000783 | -0.09 | -0.29 | 0.12 | 0.10 | -0.89 | 0.398 |
| **ACR.L** | 0.000745 | 0.000751 | 0.12 | -0.08 | 0.33 | 0.10 | 1.27 | 0.228 |
| **ACR.R** | 0.000739 | 0.000749 | 0.21 | 0.01 | 0.41 | 0.10 | 2.14 | 0.042 |
| **ALIC.L** | 0.000695 | 0.000700 | 0.07 | -0.13 | 0.28 | 0.10 | 0.76 | 0.472 |
| **ALIC.R** | 0.000695 | 0.000707 | 0.21 | 0.01 | 0.41 | 0.10 | 2.13 | 0.044 |
| **CGC.L** | 0.000738 | 0.000741 | 0.06 | -0.14 | 0.27 | 0.10 | 0.66 | 0.533 |
| **CGC.R** | 0.000729 | 0.000733 | 0.08 | -0.13 | 0.28 | 0.10 | 0.77 | 0.464 |
| **CGH.L** | 0.000876 | 0.000878 | -0.01 | -0.22 | 0.19 | 0.10 | -0.14 | 0.896 |
| **CGH.R** | 0.000859 | 0.000888 | 0.22 | 0.01 | 0.42 | 0.10 | 2.22 | 0.036 |
| **CST.L** | 0.000740 | 0.000747 | 0.05 | -0.16 | 0.25 | 0.10 | 0.46 | 0.663 |
| **CST.R** | 0.000743 | 0.000747 | 0.02 | -0.18 | 0.23 | 0.10 | 0.24 | 0.819 |
| **EC.L** | 0.000741 | 0.000752 | 0.27 | 0.07 | 0.48 | 0.10 | 2.79 | 0.008 |
| **EC.R** | 0.000737 | 0.000755 | 0.46 | 0.26 | 0.66 | 0.10 | 4.70 | 1.0E-05 |
| **FX.ST.L** | 0.000787 | 0.000795 | 0.10 | -0.10 | 0.30 | 0.10 | 1.04 | 0.325 |
| **FX.ST.R** | 0.000781 | 0.000789 | 0.11 | -0.09 | 0.32 | 0.10 | 1.15 | 0.274 |
| **PCR.L** | 0.000789 | 0.000800 | 0.21 | 0.01 | 0.41 | 0.10 | 2.15 | 0.042 |
| **PCR.R** | 0.000802 | 0.000810 | 0.14 | -0.06 | 0.34 | 0.10 | 1.42 | 0.179 |
| **PLIC.L** | 0.000683 | 0.000686 | 0.08 | -0.12 | 0.28 | 0.10 | 0.81 | 0.442 |
| **PLIC.R** | 0.000677 | 0.000683 | 0.12 | -0.08 | 0.33 | 0.10 | 1.27 | 0.231 |
| **PTR.L** | 0.000820 | 0.000830 | 0.15 | -0.05 | 0.35 | 0.10 | 1.51 | 0.152 |
| **PTR.R** | 0.000811 | 0.000828 | 0.22 | 0.02 | 0.42 | 0.10 | 2.25 | 0.034 |
| **RLIC.L** | 0.000778 | 0.000783 | 0.09 | -0.11 | 0.29 | 0.10 | 0.92 | 0.384 |
| **RLIC.R** | 0.000776 | 0.000780 | 0.07 | -0.13 | 0.27 | 0.10 | 0.73 | 0.492 |
| **SCR.L** | 0.000697 | 0.000703 | 0.16 | -0.04 | 0.37 | 0.10 | 1.67 | 0.114 |
| **SCR.R** | 0.000696 | 0.000700 | 0.10 | -0.10 | 0.31 | 0.10 | 1.07 | 0.311 |
| **SFO.L** | 0.000679 | 0.000678 | 0.00 | -0.20 | 0.20 | 0.10 | 0.02 | 1.000 |
| **SFO.R** | 0.000669 | 0.000678 | 0.10 | -0.10 | 0.31 | 0.10 | 1.07 | 0.311 |
| **SLF.L** | 0.000709 | 0.000715 | 0.19 | -0.01 | 0.39 | 0.10 | 1.96 | 0.063 |
| **SLF.R** | 0.000705 | 0.000715 | 0.30 | 0.09 | 0.50 | 0.10 | 3.02 | 0.004 |
| **SS.L** | 0.000787 | 0.000798 | 0.19 | -0.01 | 0.40 | 0.10 | 1.97 | 0.063 |
| **SS.R** | 0.000773 | 0.000796 | 0.42 | 0.22 | 0.63 | 0.10 | 4.31 | 4.8E-05 |
| **TAP.L** | 0.000936 | 0.000940 | 0.05 | -0.16 | 0.25 | 0.10 | 0.48 | 0.652 |
| **TAP.R** | 0.000922 | 0.000926 | 0.05 | -0.15 | 0.26 | 0.10 | 0.56 | 0.597 |
| **UNC.L** | 0.000761 | 0.000772 | 0.11 | -0.09 | 0.32 | 0.10 | 1.16 | 0.272 |
| **UNC.R** | 0.000740 | 0.000777 | 0.42 | 0.22 | 0.63 | 0.10 | 4.33 | 4.4E-05 |

**Supplementary Table 17.** Effect sizes for MD differences between healthy controls and the ‘GGE’ syndrome.

| ROI | Mean MD controls | Mean MD patients | Cohen’s d | 95% CI lower | 95% CI upper | Standard error | Z score | P value |
|---|---|---|---|---|---|---|---|---|
| **AverageMD** | 0.000801 | 0.000799 | -0.02 | -0.22 | 0.17 | 0.10 | -0.25 | 0.813 |
| **BCC** | 0.000881 | 0.000866 | -0.13 | -0.32 | 0.07 | 0.10 | -1.36 | 0.200 |
| **GCC** | 0.000816 | 0.000788 | -0.14 | -0.34 | 0.05 | 0.10 | -1.51 | 0.157 |
| **SCC** | 0.000791 | 0.000779 | -0.14 | -0.34 | 0.05 | 0.10 | -1.52 | 0.154 |
| **ACR.L** | 0.000745 | 0.000753 | 0.16 | -0.04 | 0.35 | 0.10 | 1.65 | 0.121 |
| **ACR.R** | 0.000739 | 0.000749 | 0.21 | 0.02 | 0.41 | 0.10 | 2.28 | 0.033 |
| **ALIC.L** | 0.000695 | 0.000688 | -0.15 | -0.35 | 0.04 | 0.10 | -1.62 | 0.127 |
| **ALIC.R** | 0.000695 | 0.000694 | -0.06 | -0.26 | 0.13 | 0.10 | -0.66 | 0.535 |
| **CGC.L** | 0.000738 | 0.000731 | -0.12 | -0.32 | 0.08 | 0.10 | -1.27 | 0.233 |
| **CGC.R** | 0.000729 | 0.000721 | -0.13 | -0.33 | 0.07 | 0.10 | -1.37 | 0.197 |
| **CGH.L** | 0.000876 | 0.000877 | -0.09 | -0.28 | 0.11 | 0.10 | -0.93 | 0.384 |
| **CGH.R** | 0.000859 | 0.000857 | -0.10 | -0.30 | 0.09 | 0.10 | -1.10 | 0.302 |
| **CST.L** | 0.000740 | 0.000726 | -0.18 | -0.38 | 0.01 | 0.10 | -1.93 | 0.070 |
| **CST.R** | 0.000743 | 0.000722 | -0.26 | -0.46 | -0.07 | 0.10 | -2.79 | 0.009 |
| **EC.L** | 0.000741 | 0.000740 | -0.06 | -0.25 | 0.14 | 0.10 | -0.61 | 0.566 |
| **EC.R** | 0.000737 | 0.000739 | 0.03 | -0.17 | 0.22 | 0.10 | 0.31 | 0.772 |
| **FX.ST.L** | 0.000787 | 0.000779 | -0.14 | -0.33 | 0.06 | 0.10 | -1.47 | 0.168 |
| **FX.ST.R** | 0.000781 | 0.000781 | -0.03 | -0.22 | 0.17 | 0.10 | -0.28 | 0.795 |
| **PCR.L** | 0.000789 | 0.000801 | 0.21 | 0.01 | 0.40 | 0.10 | 2.18 | 0.040 |
| **PCR.R** | 0.000802 | 0.000815 | 0.23 | 0.04 | 0.43 | 0.10 | 2.48 | 0.020 |
| **PLIC.L** | 0.000683 | 0.000679 | -0.20 | -0.40 | -0.01 | 0.10 | -2.13 | 0.045 |
| **PLIC.R** | 0.000677 | 0.000674 | -0.16 | -0.35 | 0.04 | 0.10 | -1.65 | 0.121 |
| **PTR.L** | 0.000820 | 0.000822 | 0.07 | -0.13 | 0.26 | 0.10 | 0.71 | 0.502 |
| **PTR.R** | 0.000811 | 0.000818 | 0.12 | -0.07 | 0.32 | 0.10 | 1.30 | 0.223 |
| **RLIC.L** | 0.000778 | 0.000777 | -0.03 | -0.22 | 0.17 | 0.10 | -0.28 | 0.790 |
| **RLIC.R** | 0.000776 | 0.000770 | -0.11 | -0.30 | 0.09 | 0.10 | -1.14 | 0.283 |
| **SCR.L** | 0.000697 | 0.000699 | 0.05 | -0.14 | 0.25 | 0.10 | 0.58 | 0.585 |
| **SCR.R** | 0.000696 | 0.000698 | 0.04 | -0.15 | 0.24 | 0.10 | 0.46 | 0.666 |
| **SFO.L** | 0.000679 | 0.000675 | -0.04 | -0.24 | 0.15 | 0.10 | -0.45 | 0.676 |
| **SFO.R** | 0.000669 | 0.000677 | 0.08 | -0.12 | 0.27 | 0.10 | 0.84 | 0.431 |
| **SLF.L** | 0.000709 | 0.000714 | 0.13 | -0.06 | 0.33 | 0.10 | 1.40 | 0.190 |
| **SLF.R** | 0.000705 | 0.000711 | 0.18 | -0.02 | 0.37 | 0.10 | 1.90 | 0.075 |
| **SS.L** | 0.000787 | 0.000786 | -0.03 | -0.22 | 0.17 | 0.10 | -0.28 | 0.791 |
| **SS.R** | 0.000773 | 0.000774 | 0.02 | -0.17 | 0.22 | 0.10 | 0.23 | 0.830 |
| **TAP.L** | 0.000936 | 0.000944 | 0.09 | -0.11 | 0.28 | 0.10 | 0.93 | 0.381 |
| **TAP.R** | 0.000922 | 0.000930 | 0.11 | -0.09 | 0.30 | 0.10 | 1.12 | 0.293 |
| **UNC.L** | 0.000761 | 0.000764 | 0.00 | -0.20 | 0.19 | 0.10 | -0.05 | 0.961 |
| **UNC.R** | 0.000740 | 0.000748 | 0.04 | -0.16 | 0.23 | 0.10 | 0.39 | 0.713 |

**Supplementary Table 19.** Effect sizes for MD differences between healthy controls and the 'ExE' syndrome.

| ROI | Mean MD controls | Mean MD patients | Cohen's d | 95% CI lower | 95% CI upper | Standard error | Z score | P value |
|---|---|---|---|---|---|---|---|---|
| **AverageMD** | 0.000801 | 0.000818 | 0.34 | 0.18 | 0.50 | 0.08 | 4.47 | 4.9E-05 |
| **BCC** | 0.000881 | 0.000902 | 0.24 | 0.08 | 0.40 | 0.08 | 3.21 | 0.004 |
| **GCC** | 0.000816 | 0.000830 | 0.17 | 0.01 | 0.33 | 0.08 | 2.25 | 0.041 |
| **SCC** | 0.000791 | 0.000806 | 0.15 | -0.02 | 0.31 | 0.08 | 1.93 | 0.079 |
| **ACR.L** | 0.000745 | 0.000763 | 0.33 | 0.17 | 0.49 | 0.08 | 4.34 | 8.4E-05 |
| **ACR.R** | 0.000739 | 0.000754 | 0.29 | 0.13 | 0.46 | 0.08 | 3.89 | 4.0E-04 |
| **ALIC.L** | 0.000695 | 0.000704 | 0.09 | -0.08 | 0.25 | 0.08 | 1.14 | 0.299 |
| **ALIC.R** | 0.000695 | 0.000701 | 0.06 | -0.11 | 0.22 | 0.08 | 0.73 | 0.504 |
| **CGC.L** | 0.000738 | 0.000749 | 0.18 | 0.01 | 0.34 | 0.08 | 2.32 | 0.035 |
| **CGC.R** | 0.000729 | 0.000743 | 0.24 | 0.08 | 0.40 | 0.08 | 3.16 | 0.004 |
| **CGH.L** | 0.000876 | 0.000905 | 0.15 | -0.01 | 0.31 | 0.08 | 1.94 | 0.078 |
| **CGH.R** | 0.000859 | 0.000877 | 0.08 | -0.08 | 0.24 | 0.08 | 1.04 | 0.345 |
| **CST.L** | 0.000740 | 0.000750 | 0.05 | -0.11 | 0.21 | 0.08 | 0.67 | 0.540 |
| **CST.R** | 0.000743 | 0.000754 | 0.07 | -0.10 | 0.23 | 0.08 | 0.87 | 0.427 |
| **EC.L** | 0.000741 | 0.000749 | 0.17 | 0.01 | 0.33 | 0.08 | 2.26 | 0.040 |
| **EC.R** | 0.000737 | 0.000748 | 0.25 | 0.09 | 0.41 | 0.08 | 3.31 | 0.003 |
| **FX.ST.L** | 0.000787 | 0.000799 | 0.12 | -0.05 | 0.28 | 0.08 | 1.54 | 0.161 |
| **FX.ST.R** | 0.000781 | 0.000797 | 0.18 | 0.01 | 0.34 | 0.08 | 2.32 | 0.035 |
| **PCR.L** | 0.000789 | 0.000804 | 0.25 | 0.09 | 0.41 | 0.08 | 3.30 | 0.003 |
| **PCR.R** | 0.000802 | 0.000817 | 0.25 | 0.08 | 0.41 | 0.08 | 3.26 | 0.003 |
| **PLIC.L** | 0.000683 | 0.000687 | 0.04 | -0.12 | 0.20 | 0.08 | 0.54 | 0.625 |
| **PLIC.R** | 0.000677 | 0.000682 | 0.05 | -0.11 | 0.21 | 0.08 | 0.66 | 0.544 |
| **PTR.L** | 0.000820 | 0.000830 | 0.14 | -0.02 | 0.30 | 0.08 | 1.85 | 0.093 |
| **PTR.R** | 0.000811 | 0.000816 | 0.07 | -0.09 | 0.23 | 0.08 | 0.94 | 0.392 |
| **RLIC.L** | 0.000778 | 0.000787 | 0.13 | -0.03 | 0.30 | 0.08 | 1.78 | 0.105 |
| **RLIC.R** | 0.000776 | 0.000783 | 0.09 | -0.07 | 0.25 | 0.08 | 1.16 | 0.291 |
| **SCR.L** | 0.000697 | 0.000709 | 0.27 | 0.11 | 0.44 | 0.08 | 3.64 | 9.6E-04 |
| **SCR.R** | 0.000696 | 0.000709 | 0.26 | 0.10 | 0.42 | 0.08 | 3.43 | 0.002 |
| **SFO.L** | 0.000679 | 0.000699 | 0.15 | -0.01 | 0.31 | 0.08 | 1.97 | 0.073 |
| **SFO.R** | 0.000669 | 0.000687 | 0.18 | 0.02 | 0.34 | 0.08 | 2.35 | 0.033 |
| **SLF.L** | 0.000709 | 0.000720 | 0.30 | 0.14 | 0.46 | 0.08 | 4.01 | 2.6E-04 |
| **SLF.R** | 0.000705 | 0.000718 | 0.37 | 0.21 | 0.53 | 0.08 | 4.90 | 8.8E-06 |
| **SS.L** | 0.000787 | 0.000802 | 0.23 | 0.06 | 0.39 | 0.08 | 2.98 | 0.007 |
| **SS.R** | 0.000773 | 0.000783 | 0.16 | -0.01 | 0.32 | 0.08 | 2.07 | 0.060 |
| **TAP.L** | 0.000936 | 0.000945 | 0.08 | -0.08 | 0.24 | 0.08 | 1.05 | 0.338 |
| **TAP.R** | 0.000922 | 0.000932 | 0.10 | -0.06 | 0.26 | 0.08 | 1.30 | 0.235 |
| **UNC.L** | 0.000761 | 0.000784 | 0.21 | 0.04 | 0.37 | 0.08 | 2.72 | 0.013 |
| **UNC.R** | 0.000740 | 0.000753 | 0.12 | -0.04 | 0.29 | 0.08 | 1.65 | 0.130 |

**Supplementary Table 20.** Effect sizes for RD differences between healthy controls and the 'All Epilepsies' syndrome.

| ROI | Mean RD controls | Mean RD patients | Cohen's d | 95% CI lower | 95% CI upper | Standard error | Z score | P value |
|---|---|---|---|---|---|---|---|---|
| **AverageRD** | 0.000556 | 0.000581 | 0.47 | 0.38 | 0.56 | 0.05 | 15.05 | 1.2E-24 |
| **BCC** | 0.000550 | 0.000582 | 0.31 | 0.22 | 0.40 | 0.05 | 10.08 | 4.6E-12 |
| **GCC** | 0.000505 | 0.000533 | 0.21 | 0.13 | 0.30 | 0.04 | 6.90 | 2.1E-06 |
| **SCC** | 0.000452 | 0.000462 | 0.10 | 0.01 | 0.19 | 0.04 | 3.27 | 0.024 |
| **ACR.L** | 0.000560 | 0.000582 | 0.38 | 0.29 | 0.47 | 0.05 | 12.24 | 5.6E-17 |
| **ACR.R** | 0.000550 | 0.000574 | 0.45 | 0.36 | 0.54 | 0.05 | 14.56 | 2.9E-23 |
| **ALIC.L** | 0.000460 | 0.000473 | 0.23 | 0.14 | 0.31 | 0.04 | 7.31 | 4.9E-07 |
| **ALIC.R** | 0.000458 | 0.000471 | 0.24 | 0.15 | 0.33 | 0.05 | 7.80 | 8.3E-08 |
| **CGC.L** | 0.000462 | 0.000488 | 0.39 | 0.30 | 0.48 | 0.05 | 12.62 | 5.7E-18 |
| **CGC.R** | 0.000492 | 0.000515 | 0.33 | 0.25 | 0.42 | 0.05 | 10.82 | 1.2E-13 |
| **CGH.L** | 0.000663 | 0.000695 | 0.26 | 0.17 | 0.35 | 0.05 | 8.33 | 9.8E-09 |
| **CGH.R** | 0.000641 | 0.000670 | 0.23 | 0.15 | 0.32 | 0.05 | 7.55 | 2.0E-07 |
| **CST.L** | 0.000514 | 0.000520 | 0.06 | -0.03 | 0.15 | 0.04 | 1.91 | 0.188 |
| **CST.R** | 0.000523 | 0.000528 | 0.05 | -0.03 | 0.14 | 0.04 | 1.73 | 0.232 |
| **EC.L** | 0.000552 | 0.000575 | 0.48 | 0.39 | 0.57 | 0.05 | 15.61 | 2.4E-26 |
| **EC.R** | 0.000551 | 0.000573 | 0.52 | 0.43 | 0.61 | 0.05 | 16.93 | 1.2E-30 |
| **FX.ST.L** | 0.000551 | 0.000570 | 0.24 | 0.15 | 0.33 | 0.05 | 7.70 | 1.1E-07 |
| **FX.ST.R** | 0.000549 | 0.000569 | 0.26 | 0.17 | 0.34 | 0.05 | 8.28 | 1.3E-08 |
| **PCR.L** | 0.000584 | 0.000601 | 0.31 | 0.22 | 0.40 | 0.05 | 10.10 | 3.8E-12 |
| **PCR.R** | 0.000593 | 0.000614 | 0.33 | 0.24 | 0.42 | 0.05 | 10.64 | 3.0E-13 |
| **PLIC.L** | 0.000390 | 0.000396 | 0.18 | 0.09 | 0.27 | 0.04 | 5.73 | 7.7E-05 |
| **PLIC.R** | 0.000384 | 0.000393 | 0.21 | 0.12 | 0.30 | 0.04 | 6.72 | 3.6E-06 |
| **PTR.L** | 0.000539 | 0.000558 | 0.24 | 0.15 | 0.33 | 0.05 | 7.70 | 1.2E-07 |
| **PTR.R** | 0.000532 | 0.000553 | 0.25 | 0.16 | 0.34 | 0.05 | 8.09 | 2.5E-08 |
| **RLIC.L** | 0.000505 | 0.000521 | 0.28 | 0.19 | 0.36 | 0.05 | 8.90 | 9.3E-10 |
| **RLIC.R** | 0.000521 | 0.000536 | 0.24 | 0.15 | 0.32 | 0.05 | 7.63 | 1.6E-07 |
| **SCR.L** | 0.000501 | 0.000515 | 0.32 | 0.23 | 0.41 | 0.05 | 10.47 | 7.2E-13 |
| **SCR.R** | 0.000501 | 0.000514 | 0.31 | 0.22 | 0.40 | 0.05 | 10.05 | 5.6E-12 |
| **SFO.L** | 0.000472 | 0.000490 | 0.17 | 0.08 | 0.26 | 0.04 | 5.56 | 1.2E-04 |
| **SFO.R** | 0.000461 | 0.000479 | 0.21 | 0.13 | 0.30 | 0.04 | 6.90 | 2.1E-06 |
| **SLF.L** | 0.000513 | 0.000528 | 0.40 | 0.31 | 0.49 | 0.05 | 13.04 | 5.0E-19 |
| **SLF.R** | 0.000509 | 0.000527 | 0.48 | 0.39 | 0.57 | 0.05 | 15.51 | 4.7E-26 |
| **SS.L** | 0.000539 | 0.000564 | 0.41 | 0.32 | 0.50 | 0.05 | 13.25 | 1.4E-19 |
| **SS.R** | 0.000532 | 0.000558 | 0.43 | 0.34 | 0.52 | 0.05 | 13.85 | 3.7E-21 |
| **TAP.L** | 0.000670 | 0.000692 | 0.14 | 0.05 | 0.23 | 0.04 | 4.46 | 0.002 |
| **TAP.R** | 0.000637 | 0.000661 | 0.15 | 0.06 | 0.24 | 0.04 | 4.81 | 9.0E-04 |
| **UNC.L** | 0.000544 | 0.000574 | 0.29 | 0.20 | 0.38 | 0.05 | 9.36 | 1.2E-10 |
| **UNC.R** | 0.000525 | 0.000557 | 0.33 | 0.24 | 0.41 | 0.05 | 10.52 | 4.8E-13 |

**Supplementary Table 21.** Effect sizes for RD differences between healthy controls and the 'L TLE-HS' syndrome.

| ROI | Mean RD controls | Mean RD patients | Cohen's d | 95% CI lower | 95% CI upper | Standard error | Z score | P value |
|---|---|---|---|---|---|---|---|---|
| **AverageRD** | 0.000556 | 0.000591 | 0.66 | 0.52 | 0.79 | 0.07 | 10.96 | 5.1E-21 |
| **BCC** | 0.000550 | 0.000597 | 0.44 | 0.30 | 0.57 | 0.07 | 7.34 | 2.1E-10 |
| **GCC** | 0.000505 | 0.000556 | 0.35 | 0.22 | 0.49 | 0.07 | 5.88 | 3.4E-07 |
| **SCC** | 0.000452 | 0.000465 | 0.13 | -0.01 | 0.26 | 0.07 | 2.12 | 0.064 |
| **ACR.L** | 0.000560 | 0.000595 | 0.57 | 0.44 | 0.71 | 0.07 | 9.56 | 1.9E-16 |
| **ACR.R** | 0.000550 | 0.000579 | 0.52 | 0.38 | 0.65 | 0.07 | 8.66 | 8.0E-14 |
| **ALIC.L** | 0.000460 | 0.000481 | 0.39 | 0.25 | 0.52 | 0.07 | 6.51 | 1.6E-08 |
| **ALIC.R** | 0.000458 | 0.000470 | 0.24 | 0.11 | 0.38 | 0.07 | 4.10 | 3.7E-04 |
| **CGC.L** | 0.000462 | 0.000507 | 0.69 | 0.55 | 0.82 | 0.07 | 11.46 | 1.1E-22 |
| **CGC.R** | 0.000492 | 0.000523 | 0.46 | 0.33 | 0.60 | 0.07 | 7.76 | 2.0E-11 |
| **CGH.L** | 0.000663 | 0.000738 | 0.67 | 0.53 | 0.81 | 0.07 | 11.21 | 7.4E-22 |
| **CGH.R** | 0.000641 | 0.000664 | 0.24 | 0.10 | 0.37 | 0.07 | 3.97 | 5.7E-04 |
| **CST.L** | 0.000514 | 0.000516 | 0.04 | -0.09 | 0.17 | 0.07 | 0.67 | 0.556 |
| **CST.R** | 0.000523 | 0.000528 | 0.08 | -0.05 | 0.22 | 0.07 | 1.40 | 0.221 |
| **EC.L** | 0.000552 | 0.000590 | 0.83 | 0.69 | 0.97 | 0.07 | 13.84 | 6.5E-32 |
| **EC.R** | 0.000551 | 0.000576 | 0.61 | 0.47 | 0.74 | 0.07 | 10.16 | 2.5E-18 |
| **FX.ST.L** | 0.000551 | 0.000598 | 0.59 | 0.46 | 0.73 | 0.07 | 9.91 | 1.5E-17 |
| **FX.ST.R** | 0.000549 | 0.000572 | 0.32 | 0.18 | 0.45 | 0.07 | 5.27 | 4.6E-06 |
| **PCR.L** | 0.000584 | 0.000610 | 0.45 | 0.31 | 0.58 | 0.07 | 7.49 | 9.0E-11 |
| **PCR.R** | 0.000593 | 0.000617 | 0.36 | 0.23 | 0.49 | 0.07 | 6.03 | 1.7E-07 |
| **PLIC.L** | 0.000390 | 0.000399 | 0.27 | 0.13 | 0.40 | 0.07 | 4.48 | 9.7E-05 |
| **PLIC.R** | 0.000384 | 0.000394 | 0.25 | 0.12 | 0.38 | 0.07 | 4.18 | 2.8E-04 |
| **PTR.L** | 0.000539 | 0.000572 | 0.36 | 0.22 | 0.49 | 0.07 | 6.01 | 1.8E-07 |
| **PTR.R** | 0.000532 | 0.000559 | 0.28 | 0.14 | 0.41 | 0.07 | 4.62 | 5.8E-05 |
| **RLIC.L** | 0.000505 | 0.000534 | 0.51 | 0.37 | 0.64 | 0.07 | 8.45 | 2.8E-13 |
| **RLIC.R** | 0.000521 | 0.000541 | 0.31 | 0.18 | 0.44 | 0.07 | 5.18 | 6.5E-06 |
| **SCR.L** | 0.000501 | 0.000520 | 0.40 | 0.27 | 0.54 | 0.07 | 6.74 | 5.3E-09 |
| **SCR.R** | 0.000501 | 0.000517 | 0.37 | 0.23 | 0.50 | 0.07 | 6.13 | 1.1E-07 |
| **SFO.L** | 0.000472 | 0.000499 | 0.25 | 0.12 | 0.39 | 0.07 | 4.23 | 2.4E-04 |
| **SFO.R** | 0.000461 | 0.000486 | 0.30 | 0.16 | 0.43 | 0.07 | 4.99 | 1.4E-05 |
| **SLF.L** | 0.000513 | 0.000535 | 0.56 | 0.43 | 0.70 | 0.07 | 9.44 | 5.1E-16 |
| **SLF.R** | 0.000509 | 0.000529 | 0.53 | 0.40 | 0.67 | 0.07 | 8.89 | 1.7E-14 |
| **SS.L** | 0.000539 | 0.000588 | 0.79 | 0.65 | 0.92 | 0.07 | 13.15 | 5.1E-29 |
| **SS.R** | 0.000532 | 0.000559 | 0.44 | 0.30 | 0.57 | 0.07 | 7.33 | 2.2E-10 |
| **TAP.L** | 0.000670 | 0.000708 | 0.22 | 0.08 | 0.35 | 0.07 | 3.61 | 0.002 |
| **TAP.R** | 0.000637 | 0.000664 | 0.14 | 0.01 | 0.27 | 0.07 | 2.32 | 0.043 |
| **UNC.L** | 0.000544 | 0.000593 | 0.47 | 0.34 | 0.61 | 0.07 | 7.88 | 9.7E-12 |
| **UNC.R** | 0.000525 | 0.000556 | 0.33 | 0.20 | 0.47 | 0.07 | 5.59 | 1.1E-06 |

**Supplementary Table 22.** Effect sizes for RD differences between healthy controls and the 'R TLE-HS' syndrome.

| ROI | Mean RD controls | Mean RD patients | Cohen's d | 95% CI lower | 95% CI upper | Standard error | Z score | P value |
|---|---|---|---|---|---|---|---|---|
| **AverageRD** | 0.000556 | 0.000593 | 0.70 | 0.55 | 0.85 | 0.08 | 10.39 | 7.3E-20 |
| **BCC** | 0.000550 | 0.000610 | 0.56 | 0.41 | 0.71 | 0.08 | 8.29 | 2.4E-13 |
| **GCC** | 0.000505 | 0.000565 | 0.39 | 0.24 | 0.54 | 0.08 | 5.79 | 2.5E-07 |
| **SCC** | 0.000452 | 0.000478 | 0.26 | 0.12 | 0.41 | 0.08 | 3.88 | 5.4E-04 |
| **ACR.L** | 0.000560 | 0.000591 | 0.51 | 0.36 | 0.66 | 0.08 | 7.57 | 2.1E-11 |
| **ACR.R** | 0.000550 | 0.000589 | 0.69 | 0.54 | 0.84 | 0.08 | 10.23 | 2.5E-19 |
| **ALIC.L** | 0.000460 | 0.000472 | 0.25 | 0.10 | 0.40 | 0.08 | 3.66 | 0.001 |
| **ALIC.R** | 0.000458 | 0.000481 | 0.43 | 0.28 | 0.58 | 0.08 | 6.36 | 1.6E-08 |
| **CGC.L** | 0.000462 | 0.000492 | 0.48 | 0.33 | 0.63 | 0.08 | 7.08 | 3.3E-10 |
| **CGC.R** | 0.000492 | 0.000530 | 0.57 | 0.42 | 0.72 | 0.08 | 8.44 | 9.2E-14 |
| **CGH.L** | 0.000663 | 0.000679 | 0.21 | 0.06 | 0.36 | 0.08 | 3.09 | 0.006 |
| **CGH.R** | 0.000641 | 0.000709 | 0.61 | 0.46 | 0.76 | 0.08 | 9.05 | 1.5E-15 |
| **CST.L** | 0.000514 | 0.000515 | 0.02 | -0.13 | 0.17 | 0.08 | 0.29 | 0.798 |
| **CST.R** | 0.000523 | 0.000526 | 0.06 | -0.08 | 0.21 | 0.08 | 0.92 | 0.409 |
| **EC.L** | 0.000552 | 0.000580 | 0.63 | 0.48 | 0.78 | 0.08 | 9.34 | 2.1E-16 |
| **EC.R** | 0.000551 | 0.000586 | 0.84 | 0.69 | 0.99 | 0.08 | 12.47 | 1.8E-27 |
| **FX.ST.L** | 0.000551 | 0.000558 | 0.12 | -0.03 | 0.27 | 0.08 | 1.77 | 0.113 |
| **FX.ST.R** | 0.000549 | 0.000589 | 0.52 | 0.37 | 0.67 | 0.08 | 7.74 | 7.0E-12 |
| **PCR.L** | 0.000584 | 0.000602 | 0.31 | 0.16 | 0.46 | 0.08 | 4.62 | 3.7E-05 |
| **PCR.R** | 0.000593 | 0.000621 | 0.42 | 0.28 | 0.57 | 0.08 | 6.27 | 2.7E-08 |
| **PLIC.L** | 0.000390 | 0.000396 | 0.19 | 0.04 | 0.34 | 0.08 | 2.84 | 0.011 |
| **PLIC.R** | 0.000384 | 0.000396 | 0.30 | 0.15 | 0.45 | 0.08 | 4.44 | 8.2E-05 |
| **PTR.L** | 0.000539 | 0.000557 | 0.19 | 0.04 | 0.33 | 0.08 | 2.76 | 0.014 |
| **PTR.R** | 0.000532 | 0.000562 | 0.31 | 0.16 | 0.45 | 0.08 | 4.54 | 5.3E-05 |
| **RLIC.L** | 0.000505 | 0.000517 | 0.21 | 0.06 | 0.36 | 0.08 | 3.09 | 0.006 |
| **RLIC.R** | 0.000521 | 0.000542 | 0.33 | 0.18 | 0.47 | 0.08 | 4.83 | 1.8E-05 |
| **SCR.L** | 0.000501 | 0.000518 | 0.38 | 0.23 | 0.53 | 0.08 | 5.61 | 6.0E-07 |
| **SCR.R** | 0.000501 | 0.000519 | 0.40 | 0.25 | 0.55 | 0.08 | 5.89 | 1.8E-07 |
| **SFO.L** | 0.000472 | 0.000499 | 0.27 | 0.12 | 0.42 | 0.08 | 3.96 | 4.0E-04 |
| **SFO.R** | 0.000461 | 0.000487 | 0.30 | 0.15 | 0.45 | 0.08 | 4.41 | 8.4E-05 |
| **SLF.L** | 0.000513 | 0.000530 | 0.45 | 0.30 | 0.60 | 0.08 | 6.61 | 4.5E-09 |
| **SLF.R** | 0.000509 | 0.000533 | 0.62 | 0.47 | 0.77 | 0.08 | 9.22 | 4.1E-16 |
| **SS.L** | 0.000539 | 0.000562 | 0.39 | 0.24 | 0.54 | 0.08 | 5.73 | 3.5E-07 |
| **SS.R** | 0.000532 | 0.000579 | 0.76 | 0.61 | 0.91 | 0.08 | 11.25 | 6.8E-23 |
| **TAP.L** | 0.000670 | 0.000717 | 0.27 | 0.12 | 0.42 | 0.08 | 3.98 | 4.0E-04 |
| **TAP.R** | 0.000637 | 0.000687 | 0.27 | 0.12 | 0.42 | 0.08 | 4.00 | 3.7E-04 |
| **UNC.L** | 0.000544 | 0.000569 | 0.27 | 0.12 | 0.42 | 0.08 | 4.02 | 3.4E-04 |
| **UNC.R** | 0.000525 | 0.000591 | 0.72 | 0.57 | 0.87 | 0.08 | 10.69 | 6.4E-21 |

**Supplementary Table 23.** Effect sizes for RD differences between healthy controls and the 'L TLE-NL' syndrome.

| ROI | Mean RD controls | Mean RD patients | Cohen's d | 95% CI lower | 95% CI upper | Standard error | Z score | P value |
|---|---|---|---|---|---|---|---|---|
| **AverageRD** | 0.000556 | 0.000568 | 0.25 | 0.08 | 0.42 | 0.09 | -24.62 | 0.004 |
| **BCC** | 0.000550 | 0.000554 | 0.05 | -0.12 | 0.22 | 0.09 | -20.69 | 0.545 |
| **GCC** | 0.000505 | 0.000492 | -0.09 | -0.26 | 0.08 | 0.09 | -20.66 | 0.280 |
| **SCC** | 0.000452 | 0.000446 | -0.07 | -0.24 | 0.10 | 0.09 | -12.40 | 0.395 |
| **ACR.L** | 0.000560 | 0.000567 | 0.14 | -0.03 | 0.31 | 0.09 | -17.31 | 0.104 |
| **ACR.R** | 0.000550 | 0.000563 | 0.27 | 0.10 | 0.44 | 0.09 | -17.99 | 0.002 |
| **ALIC.L** | 0.000460 | 0.000466 | 0.13 | -0.04 | 0.30 | 0.09 | -16.61 | 0.138 |
| **ALIC.R** | 0.000458 | 0.000464 | 0.14 | -0.03 | 0.31 | 0.09 | -11.76 | 0.116 |
| **CGC.L** | 0.000462 | 0.000473 | 0.19 | 0.02 | 0.36 | 0.09 | -19.73 | 0.031 |
| **CGC.R** | 0.000492 | 0.000501 | 0.14 | -0.03 | 0.31 | 0.09 | -17.62 | 0.114 |
| **CGH.L** | 0.000663 | 0.000664 | 0.01 | -0.16 | 0.18 | 0.09 | -18.10 | 0.877 |
| **CGH.R** | 0.000641 | 0.000637 | -0.03 | -0.20 | 0.14 | 0.09 | -16.93 | 0.699 |
| **CST.L** | 0.000514 | 0.000526 | 0.11 | -0.06 | 0.28 | 0.09 | -3.22 | 0.222 |
| **CST.R** | 0.000523 | 0.000527 | 0.04 | -0.14 | 0.21 | 0.09 | -3.35 | 0.687 |
| **EC.L** | 0.000552 | 0.000566 | 0.33 | 0.15 | 0.50 | 0.09 | -22.43 | 1.8E-04 |
| **EC.R** | 0.000551 | 0.000567 | 0.40 | 0.23 | 0.57 | 0.09 | -22.01 | 4.1E-06 |
| **FX.ST.L** | 0.000551 | 0.000558 | 0.11 | -0.06 | 0.28 | 0.09 | -14.33 | 0.227 |
| **FX.ST.R** | 0.000549 | 0.000552 | 0.05 | -0.12 | 0.22 | 0.09 | -13.93 | 0.593 |
| **PCR.L** | 0.000584 | 0.000595 | 0.22 | 0.05 | 0.39 | 0.09 | -13.25 | 0.011 |
| **PCR.R** | 0.000593 | 0.000608 | 0.26 | 0.09 | 0.43 | 0.09 | -14.43 | 0.003 |
| **PLIC.L** | 0.000390 | 0.000395 | 0.17 | 0.00 | 0.34 | 0.09 | -6.07 | 0.057 |
| **PLIC.R** | 0.000384 | 0.000393 | 0.24 | 0.07 | 0.41 | 0.09 | -6.23 | 0.005 |
| **PTR.L** | 0.000539 | 0.000552 | 0.18 | 0.01 | 0.35 | 0.09 | -14.00 | 0.036 |
| **PTR.R** | 0.000532 | 0.000546 | 0.17 | 0.00 | 0.34 | 0.09 | -14.35 | 0.047 |
| **RLIC.L** | 0.000505 | 0.000515 | 0.19 | 0.02 | 0.36 | 0.09 | -14.08 | 0.028 |
| **RLIC.R** | 0.000521 | 0.000531 | 0.17 | 0.00 | 0.34 | 0.09 | -13.03 | 0.052 |
| **SCR.L** | 0.000501 | 0.000511 | 0.24 | 0.07 | 0.41 | 0.09 | -14.60 | 0.006 |
| **SCR.R** | 0.000501 | 0.000511 | 0.26 | 0.09 | 0.43 | 0.09 | -13.45 | 0.003 |
| **SFO.L** | 0.000472 | 0.000474 | 0.03 | -0.14 | 0.20 | 0.09 | -11.88 | 0.696 |
| **SFO.R** | 0.000461 | 0.000470 | 0.12 | -0.05 | 0.29 | 0.09 | -9.02 | 0.171 |
| **SLF.L** | 0.000513 | 0.000523 | 0.30 | 0.13 | 0.47 | 0.09 | -16.24 | 0.001 |
| **SLF.R** | 0.000509 | 0.000525 | 0.43 | 0.26 | 0.60 | 0.09 | -16.84 | 7.7E-07 |
| **SS.L** | 0.000539 | 0.000554 | 0.28 | 0.11 | 0.45 | 0.09 | -19.04 | 0.001 |
| **SS.R** | 0.000532 | 0.000550 | 0.32 | 0.15 | 0.49 | 0.09 | -18.13 | 2.6E-04 |
| **TAP.L** | 0.000670 | 0.000668 | -0.01 | -0.18 | 0.16 | 0.09 | -12.06 | 0.924 |
| **TAP.R** | 0.000637 | 0.000642 | 0.04 | -0.13 | 0.21 | 0.09 | -10.09 | 0.673 |
| **UNC.L** | 0.000544 | 0.000567 | 0.24 | 0.07 | 0.41 | 0.09 | -11.64 | 0.006 |
| **UNC.R** | 0.000525 | 0.000534 | 0.09 | -0.08 | 0.26 | 0.09 | -13.80 | 0.282 |

**Supplementary Table 24.** Effect sizes for RD differences between healthy controls and the 'R TLE-NL' syndrome.

| ROI | Mean RD controls | Mean RD patients | Cohen's d | 95% CI lower | 95% CI upper | Standard error | Z score | P value |
|---|---|---|---|---|---|---|---|---|
| **AverageRD** | 0.000556 | 0.000567 | 0.23 | 0.07 | 0.39 | 0.08 | 3.19 | 0.019 |
| **BCC** | 0.000550 | 0.000558 | 0.09 | -0.07 | 0.26 | 0.08 | 1.31 | 0.333 |
| **GCC** | 0.000505 | 0.000512 | 0.09 | -0.07 | 0.25 | 0.08 | 1.22 | 0.368 |
| **SCC** | 0.000452 | 0.000450 | -0.03 | -0.19 | 0.13 | 0.08 | -0.37 | 0.786 |
| **ACR.L** | 0.000560 | 0.000566 | 0.13 | -0.04 | 0.29 | 0.08 | 1.74 | 0.200 |
| **ACR.R** | 0.000550 | 0.000562 | 0.24 | 0.08 | 0.40 | 0.08 | 3.31 | 0.015 |
| **ALIC.L** | 0.000460 | 0.000465 | 0.09 | -0.07 | 0.25 | 0.08 | 1.23 | 0.364 |
| **ALIC.R** | 0.000458 | 0.000468 | 0.18 | 0.01 | 0.34 | 0.08 | 2.43 | 0.072 |
| **CGC.L** | 0.000462 | 0.000477 | 0.21 | 0.05 | 0.38 | 0.08 | 2.98 | 0.029 |
| **CGC.R** | 0.000492 | 0.000507 | 0.21 | 0.05 | 0.37 | 0.08 | 2.90 | 0.033 |
| **CGH.L** | 0.000663 | 0.000672 | 0.04 | -0.12 | 0.20 | 0.08 | 0.61 | 0.655 |
| **CGH.R** | 0.000641 | 0.000673 | 0.24 | 0.07 | 0.40 | 0.08 | 3.28 | 0.016 |
| **CST.L** | 0.000514 | 0.000525 | 0.08 | -0.08 | 0.24 | 0.08 | 1.16 | 0.394 |
| **CST.R** | 0.000523 | 0.000527 | 0.02 | -0.14 | 0.18 | 0.08 | 0.29 | 0.831 |
| **EC.L** | 0.000552 | 0.000567 | 0.33 | 0.17 | 0.49 | 0.08 | 4.54 | 8.0E-04 |
| **EC.R** | 0.000551 | 0.000572 | 0.50 | 0.33 | 0.66 | 0.08 | 6.91 | 4.0E-07 |
| **FX.ST.L** | 0.000551 | 0.000562 | 0.13 | -0.03 | 0.30 | 0.08 | 1.87 | 0.168 |
| **FX.ST.R** | 0.000549 | 0.000563 | 0.17 | 0.01 | 0.34 | 0.08 | 2.42 | 0.074 |
| **PCR.L** | 0.000584 | 0.000594 | 0.19 | 0.03 | 0.35 | 0.08 | 2.61 | 0.053 |
| **PCR.R** | 0.000593 | 0.000603 | 0.17 | 0.01 | 0.33 | 0.08 | 2.35 | 0.083 |
| **PLIC.L** | 0.000390 | 0.000394 | 0.12 | -0.04 | 0.28 | 0.08 | 1.66 | 0.220 |
| **PLIC.R** | 0.000384 | 0.000391 | 0.15 | -0.01 | 0.31 | 0.08 | 2.04 | 0.133 |
| **PTR.L** | 0.000539 | 0.000545 | 0.10 | -0.06 | 0.26 | 0.08 | 1.43 | 0.292 |
| **PTR.R** | 0.000532 | 0.000544 | 0.16 | 0.00 | 0.32 | 0.08 | 2.20 | 0.104 |
| **RLIC.L** | 0.000505 | 0.000513 | 0.15 | -0.02 | 0.31 | 0.08 | 2.03 | 0.135 |
| **RLIC.R** | 0.000521 | 0.000530 | 0.15 | -0.01 | 0.31 | 0.08 | 2.06 | 0.129 |
| **SCR.L** | 0.000501 | 0.000509 | 0.19 | 0.02 | 0.35 | 0.08 | 2.59 | 0.057 |
| **SCR.R** | 0.000501 | 0.000507 | 0.15 | -0.01 | 0.31 | 0.08 | 2.13 | 0.117 |
| **SFO.L** | 0.000472 | 0.000463 | -0.08 | -0.24 | 0.08 | 0.08 | -1.13 | 0.405 |
| **SFO.R** | 0.000461 | 0.000459 | -0.02 | -0.18 | 0.14 | 0.08 | -0.32 | 0.813 |
| **SLF.L** | 0.000513 | 0.000520 | 0.20 | 0.04 | 0.36 | 0.08 | 2.76 | 0.042 |
| **SLF.R** | 0.000509 | 0.000520 | 0.30 | 0.14 | 0.46 | 0.08 | 4.16 | 0.002 |
| **SS.L** | 0.000539 | 0.000550 | 0.21 | 0.04 | 0.37 | 0.08 | 2.85 | 0.035 |
| **SS.R** | 0.000532 | 0.000558 | 0.44 | 0.27 | 0.60 | 0.08 | 6.04 | 9.1E-06 |
| **TAP.L** | 0.000670 | 0.000654 | -0.08 | -0.24 | 0.08 | 0.08 | -1.15 | 0.398 |
| **TAP.R** | 0.000637 | 0.000637 | 0.02 | -0.15 | 0.18 | 0.08 | 0.22 | 0.869 |
| **UNC.L** | 0.000544 | 0.000565 | 0.20 | 0.04 | 0.36 | 0.08 | 2.74 | 0.044 |
| **UNC.R** | 0.000525 | 0.000560 | 0.35 | 0.18 | 0.51 | 0.08 | 4.79 | 4.0E-04 |

**Supplementary Table 25.** Effect sizes for RD differences between healthy controls and the 'GGE' syndrome.

| ROI | Mean RD controls | Mean RD patients | Cohen's d | 95% CI lower | 95% CI upper | Standard error | Z score | P value |
|---|---|---|---|---|---|---|---|---|
| **AverageRD** | 0.000556 | 0.000563 | 0.17 | -0.02 | 0.37 | 0.10 | 1.86 | 0.081 |
| **BCC** | 0.000550 | 0.000550 | 0.04 | -0.15 | 0.24 | 0.10 | 0.46 | 0.664 |
| **GCC** | 0.000505 | 0.000498 | 0.04 | -0.16 | 0.23 | 0.10 | 0.39 | 0.713 |
| **SCC** | 0.000452 | 0.000439 | -0.14 | -0.33 | 0.05 | 0.10 | -1.49 | 0.163 |
| **ACR.L** | 0.000560 | 0.000572 | 0.26 | 0.07 | 0.45 | 0.10 | 2.79 | 0.009 |
| **ACR.R** | 0.000550 | 0.000566 | 0.36 | 0.16 | 0.55 | 0.10 | 3.83 | 3.4E-04 |
| **ALIC.L** | 0.000460 | 0.000464 | 0.06 | -0.14 | 0.25 | 0.10 | 0.62 | 0.563 |
| **ALIC.R** | 0.000458 | 0.000465 | 0.12 | -0.07 | 0.31 | 0.10 | 1.30 | 0.223 |
| **CGC.L** | 0.000462 | 0.000468 | 0.09 | -0.10 | 0.28 | 0.10 | 0.97 | 0.361 |
| **CGC.R** | 0.000492 | 0.000494 | 0.03 | -0.17 | 0.22 | 0.10 | 0.29 | 0.782 |
| **CGH.L** | 0.000663 | 0.000671 | -0.02 | -0.21 | 0.18 | 0.10 | -0.18 | 0.869 |
| **CGH.R** | 0.000641 | 0.000652 | 0.02 | -0.17 | 0.21 | 0.10 | 0.23 | 0.830 |
| **CST.L** | 0.000514 | 0.000510 | -0.07 | -0.26 | 0.13 | 0.10 | -0.72 | 0.496 |
| **CST.R** | 0.000523 | 0.000515 | -0.12 | -0.32 | 0.07 | 0.10 | -1.33 | 0.210 |
| **EC.L** | 0.000552 | 0.000558 | 0.12 | -0.07 | 0.32 | 0.10 | 1.34 | 0.208 |
| **EC.R** | 0.000551 | 0.000560 | 0.21 | 0.02 | 0.41 | 0.10 | 2.28 | 0.032 |
| **FX.ST.L** | 0.000551 | 0.000548 | -0.05 | -0.24 | 0.14 | 0.10 | -0.54 | 0.611 |
| **FX.ST.R** | 0.000549 | 0.000555 | 0.08 | -0.11 | 0.27 | 0.10 | 0.84 | 0.428 |
| **PCR.L** | 0.000584 | 0.000599 | 0.31 | 0.12 | 0.51 | 0.10 | 3.37 | 0.002 |
| **PCR.R** | 0.000593 | 0.000614 | 0.39 | 0.19 | 0.58 | 0.10 | 4.14 | 1.1E-04 |
| **PLIC.L** | 0.000390 | 0.000389 | -0.03 | -0.22 | 0.17 | 0.10 | -0.27 | 0.800 |
| **PLIC.R** | 0.000384 | 0.000384 | -0.01 | -0.20 | 0.18 | 0.10 | -0.11 | 0.916 |
| **PTR.L** | 0.000539 | 0.000549 | 0.19 | 0.00 | 0.39 | 0.10 | 2.07 | 0.052 |
| **PTR.R** | 0.000532 | 0.000551 | 0.29 | 0.10 | 0.49 | 0.10 | 3.15 | 0.003 |
| **RLIC.L** | 0.000505 | 0.000515 | 0.18 | -0.01 | 0.38 | 0.10 | 1.95 | 0.066 |
| **RLIC.R** | 0.000521 | 0.000526 | 0.11 | -0.09 | 0.30 | 0.10 | 1.16 | 0.275 |
| **SCR.L** | 0.000501 | 0.000508 | 0.21 | 0.02 | 0.40 | 0.10 | 2.26 | 0.034 |
| **SCR.R** | 0.000501 | 0.000508 | 0.19 | 0.00 | 0.39 | 0.10 | 2.06 | 0.053 |
| **SFO.L** | 0.000472 | 0.000477 | 0.06 | -0.14 | 0.25 | 0.10 | 0.61 | 0.564 |
| **SFO.R** | 0.000461 | 0.000471 | 0.11 | -0.08 | 0.31 | 0.10 | 1.20 | 0.258 |
| **SLF.L** | 0.000513 | 0.000524 | 0.32 | 0.13 | 0.52 | 0.10 | 3.48 | 0.001 |
| **SLF.R** | 0.000509 | 0.000522 | 0.37 | 0.18 | 0.57 | 0.10 | 3.99 | 1.9E-04 |
| **SS.L** | 0.000539 | 0.000544 | 0.11 | -0.08 | 0.31 | 0.10 | 1.23 | 0.249 |
| **SS.R** | 0.000532 | 0.000542 | 0.19 | 0.00 | 0.38 | 0.10 | 2.03 | 0.056 |
| **TAP.L** | 0.000670 | 0.000680 | 0.09 | -0.11 | 0.28 | 0.10 | 0.93 | 0.380 |
| **TAP.R** | 0.000637 | 0.000658 | 0.18 | -0.01 | 0.37 | 0.10 | 1.93 | 0.070 |
| **UNC.L** | 0.000544 | 0.000551 | 0.06 | -0.14 | 0.25 | 0.10 | 0.60 | 0.574 |
| **UNC.R** | 0.000525 | 0.000528 | -0.01 | -0.20 | 0.19 | 0.10 | -0.09 | 0.934 |

**Supplementary Table 26.** Effect sizes for RD differences between healthy controls and the 'ExE' syndrome.

| ROI | Mean RD controls | Mean RD patients | Cohen's d | 95% CI lower | 95% CI upper | Standard error | Z score | P value |
|---|---|---|---|---|---|---|---|---|
| **AverageRD** | 0.000556 | 0.000579 | 0.48 | 0.32 | 0.64 | 0.08 | 6.39 | 7.6E-09 |
| **BCC** | 0.000550 | 0.000585 | 0.39 | 0.22 | 0.55 | 0.08 | 5.11 | 3.5E-06 |
| **GCC** | 0.000505 | 0.000530 | 0.29 | 0.13 | 0.45 | 0.08 | 3.87 | 4.4E-04 |
| **SCC** | 0.000452 | 0.000474 | 0.23 | 0.07 | 0.40 | 0.08 | 3.10 | 0.005 |
| **ACR.L** | 0.000560 | 0.000581 | 0.42 | 0.26 | 0.59 | 0.08 | 5.62 | 3.6E-07 |
| **ACR.R** | 0.000550 | 0.000572 | 0.46 | 0.30 | 0.62 | 0.08 | 6.11 | 3.3E-08 |
| **ALIC.L** | 0.000460 | 0.000477 | 0.27 | 0.10 | 0.43 | 0.08 | 3.53 | 0.001 |
| **ALIC.R** | 0.000458 | 0.000473 | 0.26 | 0.10 | 0.42 | 0.08 | 3.45 | 0.002 |
| **CGC.L** | 0.000462 | 0.000485 | 0.37 | 0.21 | 0.53 | 0.08 | 4.91 | 8.5E-06 |
| **CGC.R** | 0.000492 | 0.000516 | 0.36 | 0.20 | 0.53 | 0.08 | 4.82 | 1.2E-05 |
| **CGH.L** | 0.000663 | 0.000701 | 0.24 | 0.08 | 0.40 | 0.08 | 3.21 | 0.003 |
| **CGH.R** | 0.000641 | 0.000667 | 0.16 | 0.00 | 0.32 | 0.08 | 2.10 | 0.056 |
| **CST.L** | 0.000514 | 0.000532 | 0.14 | -0.02 | 0.30 | 0.08 | 1.89 | 0.086 |
| **CST.R** | 0.000523 | 0.000541 | 0.15 | -0.01 | 0.32 | 0.08 | 2.04 | 0.063 |
| **EC.L** | 0.000552 | 0.000570 | 0.38 | 0.22 | 0.55 | 0.08 | 5.09 | 3.9E-06 |
| **EC.R** | 0.000551 | 0.000569 | 0.43 | 0.27 | 0.59 | 0.08 | 5.71 | 2.2E-07 |
| **FX.ST.L** | 0.000551 | 0.000569 | 0.21 | 0.05 | 0.37 | 0.08 | 2.83 | 0.010 |
| **FX.ST.R** | 0.000549 | 0.000566 | 0.22 | 0.05 | 0.38 | 0.08 | 2.85 | 0.009 |
| **PCR.L** | 0.000584 | 0.000599 | 0.29 | 0.13 | 0.45 | 0.08 | 3.82 | 5.1E-04 |
| **PCR.R** | 0.000593 | 0.000611 | 0.32 | 0.16 | 0.48 | 0.08 | 4.27 | 1.1E-04 |
| **PLIC.L** | 0.000390 | 0.000398 | 0.22 | 0.05 | 0.38 | 0.08 | 2.86 | 0.009 |
| **PLIC.R** | 0.000384 | 0.000393 | 0.21 | 0.05 | 0.37 | 0.08 | 2.78 | 0.011 |
| **PTR.L** | 0.000539 | 0.000555 | 0.26 | 0.10 | 0.42 | 0.08 | 3.42 | 0.002 |
| **PTR.R** | 0.000532 | 0.000545 | 0.21 | 0.05 | 0.37 | 0.08 | 2.82 | 0.010 |
| **RLIC.L** | 0.000505 | 0.000517 | 0.23 | 0.07 | 0.39 | 0.08 | 3.04 | 0.006 |
| **RLIC.R** | 0.000521 | 0.000532 | 0.19 | 0.03 | 0.35 | 0.08 | 2.48 | 0.024 |
| **SCR.L** | 0.000501 | 0.000515 | 0.38 | 0.22 | 0.54 | 0.08 | 5.05 | 4.4E-06 |
| **SCR.R** | 0.000501 | 0.000516 | 0.38 | 0.22 | 0.54 | 0.08 | 5.00 | 5.7E-06 |
| **SFO.L** | 0.000472 | 0.000501 | 0.29 | 0.13 | 0.45 | 0.08 | 3.81 | 5.5E-04 |
| **SFO.R** | 0.000461 | 0.000486 | 0.27 | 0.11 | 0.44 | 0.08 | 3.64 | 9.5E-04 |
| **SLF.L** | 0.000513 | 0.000528 | 0.43 | 0.27 | 0.59 | 0.08 | 5.70 | 2.6E-07 |
| **SLF.R** | 0.000509 | 0.000526 | 0.49 | 0.33 | 0.65 | 0.08 | 6.48 | 4.4E-09 |
| **SS.L** | 0.000539 | 0.000556 | 0.31 | 0.15 | 0.47 | 0.08 | 4.11 | 1.9E-04 |
| **SS.R** | 0.000532 | 0.000550 | 0.33 | 0.16 | 0.49 | 0.08 | 4.32 | 9.0E-05 |
| **TAP.L** | 0.000670 | 0.000688 | 0.13 | -0.03 | 0.29 | 0.08 | 1.72 | 0.118 |
| **TAP.R** | 0.000637 | 0.000657 | 0.16 | 0.00 | 0.32 | 0.08 | 2.15 | 0.051 |
| **UNC.L** | 0.000544 | 0.000578 | 0.32 | 0.15 | 0.48 | 0.08 | 4.18 | 1.5E-04 |
| **UNC.R** | 0.000525 | 0.000553 | 0.28 | 0.12 | 0.45 | 0.08 | 3.76 | 6.3E-04 |

**Supplementary Table 27.** Effect sizes for AD differences between healthy controls and the 'All Epilepsies' syndrome.

| ROI | Mean AD controls | Mean AD patients | Cohen's d | 95% CI lower | 95% CI upper | Standard error | Z score | P value |
|---|---|---|---|---|---|---|---|---|
| **AverageAD** | 0.00132 | 0.00132 | 0.01 | -0.08 | 0.10 | 0.04 | 0.34 | 0.816 |
| **BCC** | 0.00156 | 0.00156 | 0.00 | -0.09 | 0.08 | 0.04 | -0.13 | 0.930 |
| **GCC** | 0.00149 | 0.00150 | 0.06 | -0.02 | 0.15 | 0.04 | 2.07 | 0.157 |
| **SCC** | 0.00153 | 0.00154 | 0.06 | -0.03 | 0.15 | 0.04 | 1.91 | 0.190 |
| **ACR.L** | 0.00114 | 0.00115 | 0.16 | 0.07 | 0.24 | 0.04 | 5.16 | 4.4E-04 |
| **ACR.R** | 0.00114 | 0.00115 | 0.17 | 0.08 | 0.26 | 0.04 | 5.55 | 1.5E-04 |
| **ALIC.L** | 0.00120 | 0.00119 | -0.04 | -0.13 | 0.04 | 0.04 | -1.48 | 0.312 |
| **ALIC.R** | 0.00120 | 0.00120 | -0.03 | -0.11 | 0.06 | 0.04 | -0.90 | 0.536 |
| **CGC.L** | 0.00131 | 0.00130 | -0.17 | -0.26 | -0.08 | 0.04 | -5.51 | 1.7E-04 |
| **CGC.R** | 0.00124 | 0.00123 | -0.11 | -0.20 | -0.02 | 0.04 | -3.62 | 0.013 |
| **CGH.L** | 0.00132 | 0.00132 | -0.01 | -0.09 | 0.08 | 0.04 | -0.21 | 0.884 |
| **CGH.R** | 0.00131 | 0.00131 | 0.01 | -0.08 | 0.09 | 0.04 | 0.25 | 0.862 |
| **CST.L** | 0.00122 | 0.00123 | 0.04 | -0.04 | 0.13 | 0.04 | 1.39 | 0.340 |
| **CST.R** | 0.00121 | 0.00121 | 0.04 | -0.05 | 0.13 | 0.04 | 1.30 | 0.373 |
| **EC.L** | 0.00114 | 0.00114 | 0.06 | -0.03 | 0.15 | 0.04 | 2.02 | 0.167 |
| **EC.R** | 0.00113 | 0.00114 | 0.08 | -0.01 | 0.16 | 0.04 | 2.51 | 0.086 |
| **FX.ST.L** | 0.00130 | 0.00129 | -0.03 | -0.11 | 0.06 | 0.04 | -0.89 | 0.543 |
| **FX.ST.R** | 0.00129 | 0.00129 | 0.03 | -0.05 | 0.12 | 0.04 | 1.12 | 0.445 |
| **PCR.L** | 0.00122 | 0.00123 | 0.15 | 0.07 | 0.24 | 0.04 | 5.08 | 0.001 |
| **PCR.R** | 0.00123 | 0.00125 | 0.19 | 0.10 | 0.27 | 0.04 | 6.15 | 2.8E-05 |
| **PLIC.L** | 0.00131 | 0.00130 | -0.01 | -0.10 | 0.07 | 0.04 | -0.47 | 0.750 |
| **PLIC.R** | 0.00130 | 0.00130 | 0.05 | -0.04 | 0.13 | 0.04 | 1.53 | 0.296 |
| **PTR.L** | 0.00141 | 0.00141 | 0.03 | -0.05 | 0.12 | 0.04 | 1.06 | 0.470 |
| **PTR.R** | 0.00140 | 0.00140 | 0.01 | -0.08 | 0.10 | 0.04 | 0.36 | 0.803 |
| **RLIC.L** | 0.00135 | 0.00135 | 0.01 | -0.08 | 0.09 | 0.04 | 0.24 | 0.872 |
| **RLIC.R** | 0.00131 | 0.00131 | 0.08 | 0.00 | 0.17 | 0.04 | 2.78 | 0.057 |
| **SCR.L** | 0.00112 | 0.00112 | 0.11 | 0.02 | 0.20 | 0.04 | 3.67 | 0.012 |
| **SCR.R** | 0.00110 | 0.00111 | 0.14 | 0.05 | 0.23 | 0.04 | 4.61 | 0.002 |
| **SFO.L** | 0.00112 | 0.00112 | 0.02 | -0.06 | 0.11 | 0.04 | 0.81 | 0.579 |
| **SFO.R** | 0.00112 | 0.00112 | 0.05 | -0.04 | 0.14 | 0.04 | 1.66 | 0.257 |
| **SLF.L** | 0.00113 | 0.00113 | 0.02 | -0.07 | 0.10 | 0.04 | 0.53 | 0.718 |
| **SLF.R** | 0.00113 | 0.00113 | 0.07 | -0.02 | 0.16 | 0.04 | 2.35 | 0.107 |
| **SS.L** | 0.00132 | 0.00133 | 0.09 | 0.00 | 0.18 | 0.04 | 3.02 | 0.039 |
| **SS.R** | 0.00129 | 0.00129 | 0.12 | 0.04 | 0.21 | 0.04 | 4.07 | 0.005 |
| **TAP.L** | 0.00149 | 0.00150 | 0.08 | -0.01 | 0.16 | 0.04 | 2.47 | 0.092 |
| **TAP.R** | 0.00150 | 0.00151 | 0.06 | -0.03 | 0.14 | 0.04 | 1.89 | 0.195 |
| **UNC.L** | 0.00123 | 0.00124 | 0.03 | -0.06 | 0.12 | 0.04 | 0.97 | 0.509 |
| **UNC.R** | 0.00123 | 0.00124 | 0.13 | 0.05 | 0.22 | 0.04 | 4.36 | 0.003 |

**Supplementary Table 28.** Effect sizes for AD differences between healthy controls and the 'L TLE-HS' syndrome.

| ROI | Mean AD controls | Mean AD patients | Cohen's d | 95% CI lower | 95% CI upper | Standard error | Z score | P value |
|---|---|---|---|---|---|---|---|---|
| **AverageAD** | 0.00132 | 0.00133 | 0.17 | 0.04 | 0.30 | 0.07 | 2.87 | 0.013 |
| **BCC** | 0.00156 | 0.00157 | 0.10 | -0.03 | 0.23 | 0.07 | 1.71 | 0.135 |
| **GCC** | 0.00149 | 0.00151 | 0.18 | 0.05 | 0.32 | 0.07 | 3.11 | 0.007 |
| **SCC** | 0.00153 | 0.00154 | 0.14 | 0.01 | 0.27 | 0.07 | 2.38 | 0.038 |
| **ACR.L** | 0.00114 | 0.00116 | 0.29 | 0.16 | 0.42 | 0.07 | 4.94 | 1.9E-05 |
| **ACR.R** | 0.00114 | 0.00115 | 0.22 | 0.09 | 0.35 | 0.07 | 3.75 | 0.001 |
| **ALIC.L** | 0.00120 | 0.00120 | 0.06 | -0.07 | 0.20 | 0.07 | 1.10 | 0.338 |
| **ALIC.R** | 0.00120 | 0.00120 | 0.05 | -0.08 | 0.18 | 0.07 | 0.81 | 0.480 |
| **CGC.L** | 0.00131 | 0.00130 | -0.15 | -0.29 | -0.02 | 0.07 | -2.62 | 0.023 |
| **CGC.R** | 0.00124 | 0.00123 | -0.11 | -0.25 | 0.02 | 0.07 | -1.94 | 0.091 |
| **CGH.L** | 0.00132 | 0.00133 | 0.13 | -0.01 | 0.26 | 0.07 | 2.12 | 0.065 |
| **CGH.R** | 0.00131 | 0.00131 | 0.09 | -0.04 | 0.22 | 0.07 | 1.51 | 0.187 |
| **CST.L** | 0.00122 | 0.00123 | 0.09 | -0.04 | 0.22 | 0.07 | 1.49 | 0.196 |
| **CST.R** | 0.00121 | 0.00121 | 0.08 | -0.05 | 0.21 | 0.07 | 1.39 | 0.225 |
| **EC.L** | 0.00114 | 0.00115 | 0.23 | 0.10 | 0.37 | 0.07 | 3.96 | 0.001 |
| **EC.R** | 0.00113 | 0.00114 | 0.13 | -0.01 | 0.26 | 0.07 | 2.14 | 0.063 |
| **FX.ST.L** | 0.00130 | 0.00130 | 0.10 | -0.04 | 0.23 | 0.07 | 1.61 | 0.162 |
| **FX.ST.R** | 0.00129 | 0.00129 | 0.07 | -0.06 | 0.20 | 0.07 | 1.21 | 0.295 |
| **PCR.L** | 0.00122 | 0.00123 | 0.24 | 0.11 | 0.38 | 0.07 | 4.11 | 3.7E-04 |
| **PCR.R** | 0.00123 | 0.00124 | 0.15 | 0.02 | 0.29 | 0.07 | 2.60 | 0.023 |
| **PLIC.L** | 0.00131 | 0.00131 | 0.07 | -0.06 | 0.21 | 0.07 | 1.26 | 0.274 |
| **PLIC.R** | 0.00130 | 0.00130 | 0.13 | 0.00 | 0.27 | 0.07 | 2.26 | 0.050 |
| **PTR.L** | 0.00141 | 0.00142 | 0.12 | -0.02 | 0.25 | 0.07 | 1.97 | 0.085 |
| **PTR.R** | 0.00140 | 0.00140 | 0.02 | -0.11 | 0.15 | 0.07 | 0.32 | 0.780 |
| **RLIC.L** | 0.00135 | 0.00135 | 0.11 | -0.02 | 0.24 | 0.07 | 1.83 | 0.111 |
| **RLIC.R** | 0.00131 | 0.00131 | 0.11 | -0.02 | 0.24 | 0.07 | 1.87 | 0.104 |
| **SCR.L** | 0.00112 | 0.00113 | 0.28 | 0.15 | 0.42 | 0.07 | 4.81 | 3.2E-05 |
| **SCR.R** | 0.00110 | 0.00111 | 0.17 | 0.04 | 0.30 | 0.07 | 2.86 | 0.013 |
| **SFO.L** | 0.00112 | 0.00112 | 0.09 | -0.04 | 0.22 | 0.07 | 1.54 | 0.181 |
| **SFO.R** | 0.00112 | 0.00112 | 0.10 | -0.03 | 0.24 | 0.07 | 1.75 | 0.128 |
| **SLF.L** | 0.00113 | 0.00113 | 0.07 | -0.06 | 0.21 | 0.07 | 1.26 | 0.274 |
| **SLF.R** | 0.00113 | 0.00113 | 0.10 | -0.03 | 0.23 | 0.07 | 1.68 | 0.144 |
| **SS.L** | 0.00132 | 0.00134 | 0.34 | 0.21 | 0.47 | 0.07 | 5.73 | 6.6E-07 |
| **SS.R** | 0.00129 | 0.00129 | 0.04 | -0.09 | 0.17 | 0.07 | 0.70 | 0.542 |
| **TAP.L** | 0.00149 | 0.00150 | 0.09 | -0.04 | 0.22 | 0.07 | 1.56 | 0.174 |
| **TAP.R** | 0.00150 | 0.00150 | 0.00 | -0.13 | 0.13 | 0.07 | 0.00 | 1.000 |
| **UNC.L** | 0.00123 | 0.00125 | 0.15 | 0.01 | 0.28 | 0.07 | 2.48 | 0.031 |
| **UNC.R** | 0.00123 | 0.00124 | 0.18 | 0.05 | 0.32 | 0.07 | 3.11 | 0.007 |

**Supplementary Table 29.** Effect sizes for AD differences between healthy controls and the 'R TLE-HS' syndrome.

| ROI | Mean AD controls | Mean AD patients | Cohen's d | 95% CI lower | 95% CI upper | Standard error | Z score | P value |
|---|---|---|---|---|---|---|---|---|
| **AverageAD** | 0.00132 | 0.00133 | 0.16 | 0.01 | 0.30 | 0.07 | 2.40 | 0.034 |
| **BCC** | 0.00156 | 0.00157 | 0.09 | -0.06 | 0.23 | 0.07 | 1.30 | 0.247 |
| **GCC** | 0.00149 | 0.00151 | 0.17 | 0.02 | 0.31 | 0.07 | 2.55 | 0.023 |
| **SCC** | 0.00153 | 0.00154 | 0.10 | -0.05 | 0.24 | 0.07 | 1.47 | 0.191 |
| **ACR.L** | 0.00114 | 0.00116 | 0.24 | 0.09 | 0.38 | 0.07 | 3.61 | 0.001 |
| **ACR.R** | 0.00114 | 0.00116 | 0.35 | 0.21 | 0.50 | 0.07 | 5.38 | 2.0E-06 |
| **ALIC.L** | 0.00120 | 0.00119 | 0.01 | -0.13 | 0.16 | 0.07 | 0.19 | 0.869 |
| **ALIC.R** | 0.00120 | 0.00120 | 0.06 | -0.08 | 0.21 | 0.07 | 0.93 | 0.407 |
| **CGC.L** | 0.00131 | 0.00129 | -0.31 | -0.46 | -0.17 | 0.07 | -4.75 | 2.7E-05 |
| **CGC.R** | 0.00124 | 0.00122 | -0.18 | -0.33 | -0.04 | 0.07 | -2.78 | 0.013 |
| **CGH.L** | 0.00132 | 0.00130 | -0.04 | -0.19 | 0.10 | 0.07 | -0.64 | 0.570 |
| **CGH.R** | 0.00131 | 0.00130 | -0.02 | -0.16 | 0.13 | 0.07 | -0.27 | 0.807 |
| **CST.L** | 0.00122 | 0.00122 | 0.03 | -0.11 | 0.18 | 0.07 | 0.53 | 0.636 |
| **CST.R** | 0.00121 | 0.00121 | 0.03 | -0.12 | 0.17 | 0.07 | 0.40 | 0.722 |
| **EC.L** | 0.00114 | 0.00114 | 0.02 | -0.12 | 0.16 | 0.07 | 0.29 | 0.795 |
| **EC.R** | 0.00113 | 0.00113 | 0.10 | -0.05 | 0.24 | 0.07 | 1.49 | 0.186 |
| **FX.ST.L** | 0.00130 | 0.00128 | -0.12 | -0.26 | 0.03 | 0.07 | -1.79 | 0.110 |
| **FX.ST.R** | 0.00129 | 0.00130 | 0.15 | 0.01 | 0.29 | 0.07 | 2.27 | 0.044 |
| **PCR.L** | 0.00122 | 0.00123 | 0.21 | 0.06 | 0.35 | 0.07 | 3.18 | 0.005 |
| **PCR.R** | 0.00123 | 0.00126 | 0.41 | 0.26 | 0.55 | 0.07 | 6.18 | 4.9E-08 |
| **PLIC.L** | 0.00131 | 0.00130 | 0.00 | -0.14 | 0.15 | 0.07 | 0.05 | 1.000 |
| **PLIC.R** | 0.00130 | 0.00130 | 0.10 | -0.04 | 0.25 | 0.07 | 1.59 | 0.159 |
| **PTR.L** | 0.00141 | 0.00141 | 0.04 | -0.11 | 0.18 | 0.07 | 0.55 | 0.626 |
| **PTR.R** | 0.00140 | 0.00140 | 0.08 | -0.06 | 0.22 | 0.07 | 1.23 | 0.274 |
| **RLIC.L** | 0.00135 | 0.00134 | -0.01 | -0.16 | 0.13 | 0.07 | -0.21 | 0.853 |
| **RLIC.R** | 0.00131 | 0.00131 | 0.18 | 0.04 | 0.32 | 0.07 | 2.75 | 0.015 |
| **SCR.L** | 0.00112 | 0.00111 | 0.08 | -0.07 | 0.22 | 0.07 | 1.17 | 0.298 |
| **SCR.R** | 0.00110 | 0.00112 | 0.29 | 0.14 | 0.43 | 0.07 | 4.34 | 1.2E-04 |
| **SFO.L** | 0.00112 | 0.00111 | 0.03 | -0.11 | 0.17 | 0.07 | 0.45 | 0.691 |
| **SFO.R** | 0.00112 | 0.00112 | 0.06 | -0.09 | 0.20 | 0.07 | 0.85 | 0.451 |
| **SLF.L** | 0.00113 | 0.00113 | -0.04 | -0.19 | 0.10 | 0.07 | -0.67 | 0.550 |
| **SLF.R** | 0.00113 | 0.00113 | 0.06 | -0.09 | 0.20 | 0.07 | 0.89 | 0.430 |
| **SS.L** | 0.00132 | 0.00132 | 0.01 | -0.14 | 0.15 | 0.07 | 0.10 | 0.932 |
| **SS.R** | 0.00129 | 0.00131 | 0.44 | 0.29 | 0.58 | 0.07 | 6.65 | 4.3E-09 |
| **TAP.L** | 0.00149 | 0.00150 | 0.09 | -0.06 | 0.23 | 0.07 | 1.32 | 0.243 |
| **TAP.R** | 0.00150 | 0.00153 | 0.16 | 0.02 | 0.30 | 0.07 | 2.43 | 0.031 |
| **UNC.L** | 0.00123 | 0.00122 | -0.09 | -0.23 | 0.06 | 0.07 | -1.30 | 0.250 |
| **UNC.R** | 0.00123 | 0.00124 | 0.15 | 0.01 | 0.30 | 0.07 | 2.35 | 0.037 |

**Supplementary Table 30.** Effect sizes for AD differences between healthy controls and the 'L TLE-NL' syndrome.

| ROI | Mean AD controls | Mean AD patients | Cohen's d | 95% CI lower | 95% CI upper | Standard error | Z score | P value |
|---|---|---|---|---|---|---|---|---|
| **AverageAD** | 0.00132 | 0.00132 | -0.06 | -0.23 | 0.11 | 0.09 | -0.74 | 0.491 |
| **BCC** | 0.00156 | 0.00156 | -0.04 | -0.21 | 0.13 | 0.09 | -0.54 | 0.615 |
| **GCC** | 0.00149 | 0.00148 | -0.03 | -0.20 | 0.14 | 0.09 | -0.41 | 0.705 |
| **SCC** | 0.00153 | 0.00152 | -0.08 | -0.25 | 0.09 | 0.09 | -0.97 | 0.372 |
| **ACR.L** | 0.00114 | 0.00115 | 0.09 | -0.07 | 0.26 | 0.09 | 1.18 | 0.275 |
| **ACR.R** | 0.00114 | 0.00115 | 0.11 | -0.06 | 0.28 | 0.09 | 1.41 | 0.193 |
| **ALIC.L** | 0.00120 | 0.00120 | 0.01 | -0.16 | 0.18 | 0.09 | 0.10 | 0.928 |
| **ALIC.R** | 0.00120 | 0.00120 | 0.03 | -0.14 | 0.20 | 0.09 | 0.37 | 0.729 |
| **CGC.L** | 0.00131 | 0.00131 | 0.03 | -0.14 | 0.20 | 0.09 | 0.39 | 0.718 |
| **CGC.R** | 0.00124 | 0.00124 | 0.02 | -0.15 | 0.18 | 0.09 | 0.20 | 0.853 |
| **CGH.L** | 0.00132 | 0.00131 | -0.05 | -0.22 | 0.12 | 0.09 | -0.64 | 0.555 |
| **CGH.R** | 0.00131 | 0.00130 | -0.01 | -0.17 | 0.16 | 0.09 | -0.08 | 0.943 |
| **CST.L** | 0.00122 | 0.00124 | 0.11 | -0.06 | 0.28 | 0.09 | 1.34 | 0.214 |
| **CST.R** | 0.00121 | 0.00123 | 0.17 | 0.00 | 0.34 | 0.09 | 2.10 | 0.053 |
| **EC.L** | 0.00114 | 0.00114 | 0.07 | -0.10 | 0.24 | 0.09 | 0.89 | 0.409 |
| **EC.R** | 0.00113 | 0.00114 | 0.10 | -0.07 | 0.27 | 0.09 | 1.25 | 0.249 |
| **FX.ST.L** | 0.00130 | 0.00129 | -0.08 | -0.25 | 0.08 | 0.09 | -1.05 | 0.331 |
| **FX.ST.R** | 0.00129 | 0.00128 | -0.07 | -0.24 | 0.10 | 0.09 | -0.91 | 0.399 |
| **PCR.L** | 0.00122 | 0.00122 | 0.10 | -0.07 | 0.27 | 0.09 | 1.21 | 0.262 |
| **PCR.R** | 0.00123 | 0.00124 | 0.14 | -0.03 | 0.31 | 0.09 | 1.77 | 0.101 |
| **PLIC.L** | 0.00131 | 0.00131 | 0.07 | -0.10 | 0.24 | 0.09 | 0.89 | 0.409 |
| **PLIC.R** | 0.00130 | 0.00131 | 0.18 | 0.01 | 0.35 | 0.09 | 2.27 | 0.035 |
| **PTR.L** | 0.00141 | 0.00141 | -0.01 | -0.18 | 0.15 | 0.09 | -0.18 | 0.865 |
| **PTR.R** | 0.00140 | 0.00140 | 0.02 | -0.15 | 0.19 | 0.09 | 0.29 | 0.791 |
| **RLIC.L** | 0.00135 | 0.00135 | 0.03 | -0.14 | 0.20 | 0.09 | 0.35 | 0.744 |
| **RLIC.R** | 0.00131 | 0.00131 | 0.12 | -0.05 | 0.29 | 0.09 | 1.52 | 0.159 |
| **SCR.L** | 0.00112 | 0.00112 | 0.15 | -0.02 | 0.32 | 0.09 | 1.87 | 0.084 |
| **SCR.R** | 0.00110 | 0.00111 | 0.18 | 0.01 | 0.34 | 0.09 | 2.20 | 0.042 |
| **SFO.L** | 0.00112 | 0.00111 | 0.00 | -0.17 | 0.17 | 0.09 | -0.02 | 1.000 |
| **SFO.R** | 0.00112 | 0.00112 | 0.05 | -0.12 | 0.22 | 0.09 | 0.66 | 0.539 |
| **SLF.L** | 0.00113 | 0.00113 | 0.09 | -0.08 | 0.26 | 0.09 | 1.14 | 0.290 |
| **SLF.R** | 0.00113 | 0.00113 | 0.15 | -0.02 | 0.31 | 0.09 | 1.83 | 0.091 |
| **SS.L** | 0.00132 | 0.00132 | 0.07 | -0.10 | 0.24 | 0.09 | 0.86 | 0.427 |
| **SS.R** | 0.00129 | 0.00129 | 0.09 | -0.08 | 0.26 | 0.09 | 1.16 | 0.283 |
| **TAP.L** | 0.00149 | 0.00149 | 0.02 | -0.15 | 0.19 | 0.09 | 0.26 | 0.809 |
| **TAP.R** | 0.00150 | 0.00150 | 0.04 | -0.13 | 0.21 | 0.09 | 0.52 | 0.630 |
| **UNC.L** | 0.00123 | 0.00124 | 0.08 | -0.09 | 0.25 | 0.09 | 0.97 | 0.372 |
| **UNC.R** | 0.00123 | 0.00124 | 0.19 | 0.02 | 0.36 | 0.09 | 2.41 | 0.025 |

**Supplementary Table 31.** Effect sizes for AD differences between healthy controls and the 'R TLE-NL' syndrome.

| ROI | Mean AD controls | Mean AD patients | Cohen's d | 95% CI lower | 95% CI upper | Standard error | Z score | P value |
|---|---|---|---|---|---|---|---|---|
| **AverageAD** | 0.00132 | 0.00131 | -0.13 | -0.33 | 0.07 | 0.10 | -1.31 | 0.216 |
| **BCC** | 0.00156 | 0.00156 | -0.03 | -0.23 | 0.17 | 0.10 | -0.33 | 0.756 |
| **GCC** | 0.00149 | 0.00149 | -0.02 | -0.22 | 0.18 | 0.10 | -0.22 | 0.834 |
| **SCC** | 0.00153 | 0.00152 | -0.10 | -0.30 | 0.10 | 0.10 | -0.99 | 0.347 |
| **ACR.L** | 0.00114 | 0.00115 | 0.03 | -0.17 | 0.23 | 0.10 | 0.33 | 0.754 |
| **ACR.R** | 0.00114 | 0.00115 | 0.09 | -0.11 | 0.29 | 0.10 | 0.92 | 0.385 |
| **ALIC.L** | 0.00120 | 0.00120 | 0.00 | -0.20 | 0.20 | 0.10 | 0.01 | 1.000 |
| **ALIC.R** | 0.00120 | 0.00121 | 0.03 | -0.17 | 0.23 | 0.10 | 0.34 | 0.745 |
| **CGC.L** | 0.00131 | 0.00130 | -0.13 | -0.33 | 0.07 | 0.10 | -1.33 | 0.207 |
| **CGC.R** | 0.00124 | 0.00123 | -0.13 | -0.33 | 0.07 | 0.10 | -1.35 | 0.202 |
| **CGH.L** | 0.00132 | 0.00131 | -0.07 | -0.27 | 0.13 | 0.10 | -0.68 | 0.519 |
| **CGH.R** | 0.00131 | 0.00132 | 0.06 | -0.14 | 0.26 | 0.10 | 0.66 | 0.530 |
| **CST.L** | 0.00122 | 0.00123 | 0.06 | -0.14 | 0.26 | 0.10 | 0.67 | 0.526 |
| **CST.R** | 0.00121 | 0.00121 | -0.04 | -0.24 | 0.16 | 0.10 | -0.43 | 0.685 |
| **EC.L** | 0.00114 | 0.00115 | 0.17 | -0.03 | 0.37 | 0.10 | 1.79 | 0.091 |
| **EC.R** | 0.00113 | 0.00115 | 0.27 | 0.07 | 0.47 | 0.10 | 2.83 | 0.008 |
| **FX.ST.L** | 0.00130 | 0.00130 | 0.05 | -0.15 | 0.25 | 0.10 | 0.51 | 0.630 |
| **FX.ST.R** | 0.00129 | 0.00128 | -0.05 | -0.25 | 0.15 | 0.10 | -0.54 | 0.611 |
| **PCR.L** | 0.00122 | 0.00123 | 0.08 | -0.12 | 0.28 | 0.10 | 0.88 | 0.406 |
| **PCR.R** | 0.00123 | 0.00124 | 0.11 | -0.09 | 0.31 | 0.10 | 1.12 | 0.290 |
| **PLIC.L** | 0.00131 | 0.00131 | 0.01 | -0.19 | 0.21 | 0.10 | 0.14 | 0.895 |
| **PLIC.R** | 0.00130 | 0.00130 | 0.02 | -0.18 | 0.22 | 0.10 | 0.22 | 0.833 |
| **PTR.L** | 0.00141 | 0.00142 | 0.07 | -0.13 | 0.27 | 0.10 | 0.77 | 0.468 |
| **PTR.R** | 0.00140 | 0.00141 | 0.09 | -0.11 | 0.29 | 0.10 | 0.96 | 0.363 |
| **RLIC.L** | 0.00135 | 0.00135 | -0.03 | -0.23 | 0.17 | 0.10 | -0.27 | 0.800 |
| **RLIC.R** | 0.00131 | 0.00131 | 0.00 | -0.20 | 0.20 | 0.10 | -0.03 | 1.000 |
| **SCR.L** | 0.00112 | 0.00112 | 0.07 | -0.13 | 0.27 | 0.10 | 0.75 | 0.478 |
| **SCR.R** | 0.00110 | 0.00110 | -0.03 | -0.23 | 0.17 | 0.10 | -0.30 | 0.780 |
| **SFO.L** | 0.00112 | 0.00112 | 0.02 | -0.18 | 0.22 | 0.10 | 0.20 | 0.847 |
| **SFO.R** | 0.00112 | 0.00113 | 0.15 | -0.05 | 0.35 | 0.10 | 1.50 | 0.155 |
| **SLF.L** | 0.00113 | 0.00113 | -0.02 | -0.22 | 0.18 | 0.10 | -0.16 | 0.877 |
| **SLF.R** | 0.00113 | 0.00113 | 0.06 | -0.14 | 0.26 | 0.10 | 0.61 | 0.563 |
| **SS.L** | 0.00132 | 0.00132 | -0.03 | -0.23 | 0.17 | 0.10 | -0.32 | 0.765 |
| **SS.R** | 0.00129 | 0.00130 | 0.22 | 0.02 | 0.42 | 0.10 | 2.23 | 0.035 |
| **TAP.L** | 0.00149 | 0.00151 | 0.17 | -0.03 | 0.37 | 0.10 | 1.80 | 0.089 |
| **TAP.R** | 0.00150 | 0.00151 | 0.08 | -0.12 | 0.28 | 0.10 | 0.83 | 0.429 |
| **UNC.L** | 0.00123 | 0.00124 | 0.02 | -0.18 | 0.22 | 0.10 | 0.25 | 0.813 |
| **UNC.R** | 0.00123 | 0.00125 | 0.21 | 0.01 | 0.41 | 0.10 | 2.20 | 0.038 |

**Supplementary Table 32.** Effect sizes for AD differences between healthy controls and the 'GGE' syndrome.

| ROI | Mean AD controls | Mean AD patients | Cohen's d | 95% CI lower | 95% CI upper | Standard error | Z score | P value |
|---|---|---|---|---|---|---|---|---|
| **AverageAD** | 0.00132 | 0.00131 | -0.19 | -0.39 | 0.00 | 0.10 | -2.11 | 0.048 |
| **BCC** | 0.00156 | 0.00154 | -0.20 | -0.40 | -0.01 | 0.10 | -2.22 | 0.038 |
| **GCC** | 0.00149 | 0.00148 | -0.09 | -0.28 | 0.10 | 0.10 | -0.96 | 0.369 |
| **SCC** | 0.00153 | 0.00154 | 0.06 | -0.13 | 0.26 | 0.10 | 0.70 | 0.512 |
| **ACR.L** | 0.00114 | 0.00115 | 0.04 | -0.15 | 0.23 | 0.10 | 0.41 | 0.699 |
| **ACR.R** | 0.00114 | 0.00115 | 0.07 | -0.12 | 0.26 | 0.10 | 0.76 | 0.474 |
| **ALIC.L** | 0.00120 | 0.00118 | -0.29 | -0.48 | -0.10 | 0.10 | -3.15 | 0.003 |
| **ALIC.R** | 0.00120 | 0.00119 | -0.23 | -0.42 | -0.03 | 0.10 | -2.45 | 0.022 |
| **CGC.L** | 0.00131 | 0.00129 | -0.22 | -0.41 | -0.03 | 0.10 | -2.40 | 0.025 |
| **CGC.R** | 0.00124 | 0.00122 | -0.21 | -0.40 | -0.02 | 0.10 | -2.30 | 0.031 |
| **CGH.L** | 0.00132 | 0.00131 | -0.14 | -0.34 | 0.05 | 0.10 | -1.56 | 0.142 |
| **CGH.R** | 0.00131 | 0.00129 | -0.16 | -0.35 | 0.03 | 0.10 | -1.73 | 0.106 |
| **CST.L** | 0.00122 | 0.00122 | -0.08 | -0.28 | 0.11 | 0.10 | -0.91 | 0.393 |
| **CST.R** | 0.00121 | 0.00120 | -0.08 | -0.27 | 0.11 | 0.10 | -0.87 | 0.412 |
| **EC.L** | 0.00114 | 0.00114 | -0.17 | -0.36 | 0.03 | 0.10 | -1.80 | 0.091 |
| **EC.R** | 0.00113 | 0.00113 | -0.14 | -0.34 | 0.05 | 0.10 | -1.58 | 0.139 |
| **FX.ST.L** | 0.00130 | 0.00129 | -0.16 | -0.35 | 0.03 | 0.10 | -1.77 | 0.098 |
| **FX.ST.R** | 0.00129 | 0.00128 | -0.12 | -0.32 | 0.07 | 0.10 | -1.36 | 0.203 |
| **PCR.L** | 0.00122 | 0.00122 | 0.02 | -0.18 | 0.21 | 0.10 | 0.17 | 0.870 |
| **PCR.R** | 0.00123 | 0.00124 | 0.04 | -0.16 | 0.23 | 0.10 | 0.39 | 0.714 |
| **PLIC.L** | 0.00131 | 0.00130 | -0.29 | -0.48 | -0.09 | 0.10 | -3.10 | 0.004 |
| **PLIC.R** | 0.00130 | 0.00129 | -0.27 | -0.46 | -0.08 | 0.10 | -2.92 | 0.006 |
| **PTR.L** | 0.00141 | 0.00140 | -0.12 | -0.31 | 0.07 | 0.10 | -1.31 | 0.219 |
| **PTR.R** | 0.00140 | 0.00139 | -0.13 | -0.32 | 0.06 | 0.10 | -1.40 | 0.190 |
| **RLIC.L** | 0.00135 | 0.00133 | -0.21 | -0.40 | -0.02 | 0.10 | -2.25 | 0.035 |
| **RLIC.R** | 0.00131 | 0.00130 | -0.14 | -0.33 | 0.05 | 0.10 | -1.54 | 0.149 |
| **SCR.L** | 0.00112 | 0.00111 | -0.09 | -0.29 | 0.10 | 0.10 | -1.03 | 0.335 |
| **SCR.R** | 0.00110 | 0.00111 | 0.02 | -0.17 | 0.21 | 0.10 | 0.19 | 0.861 |
| **SFO.L** | 0.00112 | 0.00111 | -0.08 | -0.27 | 0.12 | 0.10 | -0.82 | 0.442 |
| **SFO.R** | 0.00112 | 0.00111 | -0.03 | -0.22 | 0.16 | 0.10 | -0.29 | 0.786 |
| **SLF.L** | 0.00113 | 0.00113 | -0.13 | -0.32 | 0.07 | 0.10 | -1.37 | 0.198 |
| **SLF.R** | 0.00113 | 0.00112 | -0.08 | -0.27 | 0.11 | 0.10 | -0.84 | 0.428 |
| **SS.L** | 0.00132 | 0.00131 | -0.14 | -0.33 | 0.05 | 0.10 | -1.55 | 0.145 |
| **SS.R** | 0.00129 | 0.00128 | -0.09 | -0.28 | 0.10 | 0.10 | -1.00 | 0.346 |
| **TAP.L** | 0.00149 | 0.00149 | 0.05 | -0.14 | 0.24 | 0.10 | 0.53 | 0.618 |
| **TAP.R** | 0.00150 | 0.00149 | 0.02 | -0.17 | 0.21 | 0.10 | 0.19 | 0.860 |
| **UNC.L** | 0.00123 | 0.00124 | 0.01 | -0.18 | 0.20 | 0.10 | 0.11 | 0.916 |
| **UNC.R** | 0.00123 | 0.00124 | 0.08 | -0.11 | 0.27 | 0.10 | 0.88 | 0.408 |

**Supplementary Table 33.** Effect sizes for AD differences between healthy controls and the 'ExE' syndrome.

| ROI | Mean AD controls | Mean AD patients | Cohen's d | 95% CI lower | 95% CI upper | Standard error | Z score | P value |
|---|---|---|---|---|---|---|---|---|
| **AverageAD** | 0.00132 | 0.00132 | -0.16 | -0.32 | 0.00 | 0.08 | -2.19 | 0.047 |
| **BCC** | 0.00156 | 0.00155 | -0.12 | -0.28 | 0.04 | 0.08 | -1.61 | 0.142 |
| **GCC** | 0.00149 | 0.00149 | -0.05 | -0.21 | 0.11 | 0.08 | -0.63 | 0.567 |
| **SCC** | 0.00153 | 0.00154 | 0.06 | -0.10 | 0.22 | 0.08 | 0.85 | 0.440 |
| **ACR.L** | 0.00114 | 0.00115 | 0.03 | -0.13 | 0.19 | 0.08 | 0.44 | 0.686 |
| **ACR.R** | 0.00114 | 0.00115 | 0.00 | -0.16 | 0.16 | 0.08 | -0.05 | 1.000 |
| **ALIC.L** | 0.00120 | 0.00119 | -0.23 | -0.39 | -0.07 | 0.08 | -3.13 | 0.004 |
| **ALIC.R** | 0.00120 | 0.00119 | -0.23 | -0.39 | -0.07 | 0.08 | -3.14 | 0.004 |
| **CGC.L** | 0.00131 | 0.00130 | -0.16 | -0.32 | 0.00 | 0.08 | -2.18 | 0.047 |
| **CGC.R** | 0.00124 | 0.00123 | -0.04 | -0.20 | 0.12 | 0.08 | -0.56 | 0.611 |
| **CGH.L** | 0.00132 | 0.00133 | -0.03 | -0.19 | 0.13 | 0.08 | -0.35 | 0.751 |
| **CGH.R** | 0.00131 | 0.00131 | -0.03 | -0.19 | 0.13 | 0.08 | -0.42 | 0.703 |
| **CST.L** | 0.00122 | 0.00123 | 0.00 | -0.16 | 0.16 | 0.08 | -0.01 | 1.000 |
| **CST.R** | 0.00121 | 0.00122 | 0.03 | -0.12 | 0.19 | 0.08 | 0.47 | 0.671 |
| **EC.L** | 0.00114 | 0.00114 | -0.11 | -0.27 | 0.05 | 0.08 | -1.45 | 0.186 |
| **EC.R** | 0.00113 | 0.00113 | -0.04 | -0.20 | 0.12 | 0.08 | -0.48 | 0.664 |
| **FX.ST.L** | 0.00130 | 0.00130 | -0.03 | -0.19 | 0.13 | 0.08 | -0.44 | 0.689 |
| **FX.ST.R** | 0.00129 | 0.00130 | 0.05 | -0.11 | 0.21 | 0.08 | 0.73 | 0.505 |
| **PCR.L** | 0.00122 | 0.00123 | 0.14 | -0.02 | 0.30 | 0.08 | 1.86 | 0.090 |
| **PCR.R** | 0.00123 | 0.00125 | 0.14 | -0.02 | 0.30 | 0.08 | 1.89 | 0.086 |
| **PLIC.L** | 0.00131 | 0.00131 | -0.10 | -0.26 | 0.06 | 0.08 | -1.29 | 0.239 |
| **PLIC.R** | 0.00130 | 0.00131 | -0.06 | -0.22 | 0.10 | 0.08 | -0.77 | 0.484 |
| **PTR.L** | 0.00141 | 0.00141 | 0.00 | -0.16 | 0.16 | 0.08 | -0.02 | 1.000 |
| **PTR.R** | 0.00140 | 0.00140 | -0.08 | -0.24 | 0.08 | 0.08 | -1.05 | 0.338 |
| **RLIC.L** | 0.00135 | 0.00135 | 0.01 | -0.15 | 0.17 | 0.08 | 0.11 | 0.918 |
| **RLIC.R** | 0.00131 | 0.00132 | 0.06 | -0.10 | 0.22 | 0.08 | 0.76 | 0.487 |
| **SCR.L** | 0.00112 | 0.00112 | 0.03 | -0.13 | 0.19 | 0.08 | 0.36 | 0.742 |
| **SCR.R** | 0.00110 | 0.00111 | 0.09 | -0.07 | 0.25 | 0.08 | 1.20 | 0.274 |
| **SFO.L** | 0.00112 | 0.00112 | -0.02 | -0.18 | 0.14 | 0.08 | -0.24 | 0.825 |
| **SFO.R** | 0.00112 | 0.00111 | -0.07 | -0.23 | 0.09 | 0.08 | -0.93 | 0.398 |
| **SLF.L** | 0.00113 | 0.00114 | 0.05 | -0.11 | 0.21 | 0.08 | 0.68 | 0.535 |
| **SLF.R** | 0.00113 | 0.00113 | 0.10 | -0.06 | 0.26 | 0.08 | 1.34 | 0.223 |
| **SS.L** | 0.00132 | 0.00133 | 0.06 | -0.10 | 0.22 | 0.08 | 0.83 | 0.448 |
| **SS.R** | 0.00129 | 0.00129 | -0.01 | -0.17 | 0.15 | 0.08 | -0.16 | 0.885 |
| **TAP.L** | 0.00149 | 0.00149 | 0.04 | -0.12 | 0.20 | 0.08 | 0.53 | 0.632 |
| **TAP.R** | 0.00150 | 0.00150 | 0.02 | -0.14 | 0.18 | 0.08 | 0.28 | 0.800 |
| **UNC.L** | 0.00123 | 0.00123 | -0.07 | -0.23 | 0.09 | 0.08 | -0.99 | 0.365 |
| **UNC.R** | 0.00123 | 0.00122 | -0.04 | -0.20 | 0.12 | 0.08 | -0.57 | 0.602 |

**Supplementary Table 34. Relationship between FA and age of disease onset by syndrome**. Partial correlations between FA in each ROI and age of disease onset controlling for age, age$^2$ and sex. Results are split by syndrome: all syndromes (All), temporal lobe epilepsy with hippocampal sclerosis in the left and right hemisphere (TLE-HS-l and TLE-HS-r respectively), non-lesional temporal lobe epilepsy in the left and right hemisphere (TLE-NL-l and TLE-NL-r respectively), genetic generalized epilepsy (GGE) and extra temporal (ExE). ROIs are separated by left (.L) and right (.R) hemisphere where indicated. Significance was set at *** $p$ =< 0.001 (controlling for multiple comparisons, one sided). * $p$ < 0.05 ** $p$ < 0.01 *** $p$ =< 0.001.

| ROI | All | TLE-HS-l | TLE-HS-r | TLE-NL-l | TLE-NL-r | GGE | ExE |
|---|---|---|---|---|---|---|---|
| **AverageFA** | .19*** | .20*** | .24*** | .08 | .14 | .08 | .02 |
| **BCC** | .15*** | .19*** | .24*** | .08 | .00 | -.05 | .03 |
| **GCC** | .16*** | .20*** | .19** | .06 | .09 | -.06 | .05 |
| **SCC** | .13*** | .11* | .28*** | .02 | -.09 | .00 | .11 |
| **ACR.L** | .13*** | .18** | .16** | .03 | .07 | -.03 | .02 |
| **ACR.R** | .11*** | .12* | .07 | .13 | .15 | -.02 | .00 |
| **ALIC.L** | .09** | .08 | .12* | .02 | .16 | -.04 | -.03 |
| **ALIC.R** | .11*** | .15** | .10 | .12 | .11 | .02 | -.01 |
| **CGC.L** | .16*** | .17** | .15* | .17* | .08 | .01 | .01 |
| **CGC.R** | .16*** | .17** | .19** | .14* | .11 | .09 | -.01 |
| **CGH.L** | .11*** | .08 | .22*** | -.03 | -.12 | .11 | .06 |
| **CGH.R** | .12*** | -.03 | .27*** | .09 | .12 | .11 | -.01 |
| **CST.L** | .06* | .07 | .07 | .07 | .04 | .11 | -.01 |
| **CST.R** | .03 | .03 | -.01 | .01 | .09 | .03 | -.01 |
| **EC.L** | .16*** | .20*** | .19** | .08 | .13 | .08 | -.05 |
| **EC.R** | .17*** | .17** | .25*** | .07 | .15 | .16* | .00 |
| **FX.ST.L** | .11*** | .12* | .13* | .09 | .02 | .00 | .01 |
| **FX.ST.R** | .13*** | .14** | .19** | .09 | .06 | .07 | -.03 |
| **PCR.L** | .09** | .10* | .15* | .05 | -.11 | .07 | .02 |
| **PCR.R** | .13*** | .11* | .22*** | .06 | -.08 | .05 | .07 |
| **PLIC.L** | .03 | .03 | .04 | -.01 | -.03 | -.05 | .06 |
| **PLIC.R** | .05 | .05 | .06 | -.01 | -.07 | .07 | .05 |
| **PTR.L** | .11*** | .16** | .13* | .11 | -.02 | .03 | .02 |
| **PTR.R** | .10*** | .12* | .14* | .11 | -.04 | .01 | .00 |
| **RLIC.L** | .10*** | .14* | .09 | .05 | -.02 | .04 | .08 |
| **RLIC.R** | .07* | .05 | .10 | .00 | .03 | .05 | -.03 |
| **SCR.L** | .07** | .11* | .08 | -.02 | .02 | -.03 | .07 |
| **SCR.R** | .09*** | .14* | .08 | -.01 | -.03 | .02 | .10 |
| **SFO.L** | .09** | .12* | .03 | .138* | .08 | -.01 | -.03 |
| **SFO.R** | .11*** | .15** | .08 | .08 | .08 | .08 | .02 |
| **SLF.L** | .13*** | .13* | .16** | .155* | -.01 | .01 | .05 |
| **SLF.R** | .14*** | .07 | .23*** | .16* | .04 | .11 | .05 |
| **SS.L** | .12*** | .16** | .15* | .06 | .00 | .02 | .08 |
| **SS.R** | .12*** | .12* | .17** | .13 | -.01 | .07 | .02 |
| **TAP.L** | .13*** | .11* | .18** | .09 | .07 | -.06 | .08 |
| **TAP.R** | .13*** | .09 | .19** | .02 | .10 | .00 | .06 |
| **UNC.L** | .08** | .13* | .09 | -.07 | .09 | .11 | -.06 |
| **UNC.R** | .15*** | .05 | .27*** | .05 | .30*** | .09 | .00 |

**Supplementary Table 35. Relationship between FA and disease duration by syndrome**. Partial correlations between FA in each ROI and disease duration controlling for age, age$^2$ and sex. Results are split by syndrome: all syndromes (All), temporal lobe epilepsy with hippocampal sclerosis in the left and right hemisphere (TLE-HS-l and TLE-HS-r respectively), non-lesional temporal lobe epilepsy in the left and right hemisphere (TLE-NL-l and TLE-NL-r respectively), genetic generalized epilepsy (GGE) and extra temporal (Extra). ROIs are separated by left (.L) and right (.R) hemisphere where indicated. Significance was set at *** $p =< 0.001$ (controlling for multiple comparisons, one sided). * $p < 0.05$ ** $p < 0.01$ *** $p =< 0.001$.

| ROI | All | TLE-HS-l | TLE-HS-r | TLE-NL-l | TLE-NL-r | GGE | ExE |
|---|---|---|---|---|---|---|---|
| **AverageFA** | -.17*** | -.17** | -.21*** | -.09 | -.23* | -.09 | -.02 |
| **BCC** | -.16*** | -.21*** | -.20** | -.15* | -.13 | .04 | -.03 |
| **GCC** | -.13*** | -.16** | -.18** | -.06 | -.15 | .02 | -.05 |
| **SCC** | -.10*** | -.06 | -.22*** | -.01 | .01 | -.01 | -.11 |
| **ACR.L** | -.11*** | -.14* | -.13* | -.06 | -.16 | .00 | -.01 |
| **ACR.R** | -.10*** | -.09 | -.05 | -.15* | -.25** | -.01 | .02 |
| **ALIC.L** | -.08** | -.08 | -.09 | -.04 | -.23* | .06 | .04 |
| **ALIC.R** | -.09** | -.10 | -.09 | -.14* | -.19* | -.02 | .06 |
| **CGC.L** | -.17** | -.20*** | -.11 | -.22** | -.16 | .01 | -.02 |
| **CGC.R** | -.17*** | -.20*** | -.14* | -.19* | -.21* | -.06 | .00 |
| **CGH.L** | -.09** | -.03 | -.13* | -.01 | .05 | -.10 | -.03 |
| **CGH.R** | -.10** | .06 | -.28*** | -.07 | -.08 | -.09 | .03 |
| **CST.L** | -.05 | -.02 | -.12* | -.05 | -.13 | -.11 | .04 |
| **CST.R** | -.01 | -.03 | .01 | .04 | -.12 | -.04 | .08 |
| **EC.L** | -.16** | -.22*** | -.13* | -.10 | -.17 | -.07 | .05 |
| **EC.R** | -.16** | -.15** | -.22*** | -.10 | -.19* | -.17* | -.01 |
| **FX.ST.L** | -.09** | -.11* | -.02 | -.12 | -.05 | -.01 | -.03 |
| **FX.ST.R** | -.13** | -.15** | -.12* | -.11 | -.09 | -.07 | -.01 |
| **PCR.L** | -.06* | -.04 | -.11 | .01 | .03 | -.07 | -.02 |
| **PCR.R** | -.10*** | -.03 | -.26*** | .00 | .01 | -.09 | -.07 |
| **PLIC.L** | -.03 | -.02 | -.06 | .01 | -.06 | .07 | -.05 |
| **PLIC.R** | -.04 | -.02 | -.06 | -.03 | .01 | -.07 | -.01 |
| **PTR.L** | -.09** | -.12* | -.14* | -.07 | -.06 | -.07 | -.03 |
| **PTR.R** | -.08** | -.09 | -.14* | -.06 | -.05 | -.07 | -.02 |
| **RLIC.L** | -.08** | -.13* | -.04 | -.10 | -.05 | -.04 | -.04 |
| **RLIC.R** | -.05* | -.03 | -.09 | -.03 | -.10 | -.05 | .07 |
| **SCR.L** | -.06* | -.09 | -.09 | .04 | -.06 | .03 | -.02 |
| **SCR.R** | -.09** | -.10* | -.15* | .00 | .02 | -.03 | -.06 |
| **SFO.L** | -.06* | -.10 | .02 | -.11 | -.08 | .02 | .01 |
| **SFO.R** | -.09** | -.12* | -.09 | -.12 | -.04 | -.09 | .02 |
| **SLF.L** | -.13*** | -.12* | -.12* | -.14 | -.07 | .00 | -.07 |
| **SLF.R** | -.12*** | -.05 | -.20** | -.14* | -.07 | -.10 | -.05 |
| **SS.L** | -.12*** | -.15** | -.09 | -.07 | -.16 | -.03 | -.09 |
| **SS.R** | -.11*** | -.07 | -.18** | -.13 | -.09 | -.09 | -.01 |
| **TAP.L** | -.14*** | -.08 | -.26*** | -.05 | -.16 | .05 | -.10 |
| **TAP.R** | -.13*** | -.03 | -.26*** | -.02 | -.12 | -.03 | -.08 |
| **UNC.L** | -.07** | -.15** | -.02 | .10 | -.07 | -.12 | .03 |
| **UNC.R** | -.14** | -.05 | -.24*** | -.01 | -.37*** | -.09 | -.05 |

**Supplementary Table 36. Relationship between MD and age of disease onset by syndrome**. Partial correlations between MD in each ROI and age of disease onset controlling for age, age$^2$ and sex. Results are split by syndrome: all syndromes (All), temporal lobe epilepsy with hippocampal sclerosis in the left and right hemisphere (TLE-HS-l and TLE-HS-r respectively), non-lesional temporal lobe epilepsy in the left and right hemisphere (TLE-NL-l and TLE-NL-r respectively), genetic generalized epilepsy (GGE) and extra temporal (Extra). ROIs are separated by left (.L) and right (.R) hemisphere where indicated. Significance was set at *** $p$ =< 0.001 (controlling for multiple comparisons, one sided). * $p$ < 0.05 ** $p$ < 0.01 *** $p$ =< 0.001.

| ROI | All | TLE-HS-l | TLE-HS-r | TLE-NL-l | TLE-NL-r | GGE | ExE |
|---|---|---|---|---|---|---|---|
| **AverageMD** | -.16*** | -.11* | -.27*** | -.14* | -.09 | -.02 | .00 |
| **BCC** | -.13*** | -.08 | -.20** | -.06 | -.10 | .03 | -.02 |
| **GCC** | -.10*** | -.12* | -.10 | -.02 | -.05 | .06 | -.03 |
| **SCC** | -.11*** | -.08 | -.21*** | -.02 | -.08 | -.10 | .04 |
| **ACR.L** | -.10*** | -.14** | -.11* | .02 | -.05 | .03 | -.05 |
| **ACR.R** | -.10** | -.05 | -.11 | -.03 | -.06 | .05 | -.09 |
| **ALIC.L** | -.07* | -.08 | -.04 | -.04 | .06 | .04 | -.08 |
| **ALIC.R** | -.08** | -.09 | -.05 | -.04 | .14 | .02 | -.13 |
| **CGC.L** | -.11*** | -.11* | -.07 | -.10 | -.04 | -.10 | -.06 |
| **CGC.R** | -.10*** | .00 | -.11 | -.15* | -.04 | -.10 | -.12 |
| **CGH.L** | -.08** | -.10 | -.15* | -.02 | .14 | .03 | -.05 |
| **CGH.R** | -.10*** | -.01 | -.23*** | -.05 | .02 | -.05 | -.01 |
| **CST.L** | -.03 | -.05 | -.13* | -.01 | .14 | .04 | .02 |
| **CST.R** | -.01 | .01 | -.13* | .08 | .13 | .02 | .03 |
| **EC.L** | -.09*** | -.06 | -.11 | -.04 | .01 | -.06 | -.03 |
| **EC.R** | -.10*** | -.08 | -.14* | -.01 | .06 | -.05 | -.12 |
| **FX.ST.L** | -.08** | -.04 | -.15* | -.13 | .00 | .04 | -.02 |
| **FX.ST.R** | -.11*** | .00 | -.20*** | -.12 | -.01 | .05 | -.08 |
| **PCR.L** | -.08** | -.02 | -.15* | .03 | -.06 | -.05 | -.02 |
| **PCR.R** | -.09** | -.04 | -.17** | .02 | -.07 | -.04 | -.01 |
| **PLIC.L** | -.03 | .02 | -.03 | -.08 | .05 | .12 | -.18* |
| **PLIC.R** | -.05 | .01 | -.06 | -.09 | .13 | .03 | -.12 |
| **PTR.L** | -.08** | -.05 | -.19** | .04 | .02 | .01 | -.06 |
| **PTR.R** | -.07* | -.04 | -.13* | .05 | -.03 | .00 | -.01 |
| **RLIC.L** | -.10*** | -.08 | -.10 | -.06 | -.07 | -.07 | -.11 |
| **RLIC.R** | -.09** | .02 | -.19** | .00 | -.06 | -.02 | -.09 |
| **SCR.L** | -.065* | -.05 | -.13* | .06 | -.02 | .07 | -.08 |
| **SCR.R** | -.064* | -.05 | -.08 | .03 | .09 | .06 | -.12 |
| **SFO.L** | -.04 | .00 | -.06 | -.02 | .05 | .15 | -.07 |
| **SFO.R** | -.06* | .02 | -.10 | -.01 | .00 | .08 | -.14* |
| **SLF.L** | -.07** | -.06 | -.11 | .01 | .07 | -.04 | -.05 |
| **SLF.R** | -.07* | -.05 | -.10 | -.04 | .12 | -.01 | -.06 |
| **SS.L** | -.10*** | -.14** | -.15* | .05 | .10 | -.01 | -.07 |
| **SS.R** | -.10*** | -.02 | -.22*** | .06 | -.02 | -.01 | -.04 |
| **TAP.L** | -.12*** | -.15** | -.22*** | .02 | -.10 | -.03 | .05 |
| **TAP.R** | -.10*** | -.05 | -.16** | -.16* | -.17* | -.01 | .08 |
| **UNC.L** | -.06* | -.11* | -.04 | .06 | .07 | -.23** | .06 |
| **UNC.R** | -.09** | .01 | -.20*** | .07 | -.02 | -.01 | -.09 |

**Supplementary Table 37. Relationship between MD and disease duration by syndrome.** Partial correlations between MD in each ROI and disease duration controlling for age, age$^2$ and sex. Results are split by syndrome: all syndromes (All), temporal lobe epilepsy with hippocampal sclerosis in the left and right hemisphere (TLE-HS-l and TLE-HS-r respectively), non-lesional temporal lobe epilepsy in the left and right hemisphere (TLE-NL-l and TLE-NL-r respectively), genetic generalized epilepsy (GGE) and extra temporal (Extra). ROIs are separated by left (.L) and right (.R) hemisphere where indicated. Significance was set at *** $p$ =< 0.001 (controlling for multiple comparisons, one sided). * $p < 0.05$ ** $p < 0.01$ *** $p$ =< 0.001.

| ROI | All | TLE-HS-l | TLE-HS-r | TLE-NL-l | TLE-NL-r | GGE | ExE |
|---|---|---|---|---|---|---|---|
| **AverageMD** | .16*** | .12* | .26*** | .14* | .05 | .00 | .05 |
| **BCC** | .13*** | .09 | .18** | .11 | .06 | -.04 | .04 |
| **GCC** | .11*** | .16** | .12* | -.04 | .00 | -.07 | .07 |
| **SCC** | .09** | .05 | .21*** | .02 | -.04 | .09 | -.02 |
| **ACR.L** | .11*** | .15** | .17** | -.07 | .02 | -.05 | .06 |
| **ACR.R** | .10*** | .06 | .16* | -.03 | .03 | -.07 | .09 |
| **ALIC.L** | .03 | .03 | -.03 | .02 | -.13 | -.06 | .12 |
| **ALIC.R** | .04 | .05 | -.01 | -.01 | -.18* | -.04 | .16* |
| **CGC.L** | .08** | .06 | .06 | .03 | .06 | .08 | .07 |
| **CGC.R** | .09** | -.01 | .13* | .09 | .03 | .08 | .15* |
| **CGH.L** | .06* | .03 | .14* | -.02 | -.10 | -.03 | .06 |
| **CGH.R** | .09** | -.03 | .26*** | .04 | .06 | .05 | .02 |
| **CST.L** | .00 | -.02 | .14* | .01 | -.18* | -.03 | .01 |
| **CST.R** | -.02 | -.03 | .10 | -.10 | -.21* | -.01 | .00 |
| **EC.L** | .08** | .07 | .08 | -.03 | -.01 | .03 | .08 |
| **EC.R** | .09** | .03 | .15* | -.01 | .00 | .03 | .17* |
| **FX.ST.L** | .05* | .03 | .12* | .06 | -.03 | -.07 | -.02 |
| **FX.ST.R** | .10*** | -.02 | .18** | .05 | .12 | -.07 | .11 |
| **PCR.L** | .08** | .04 | .16** | -.06 | -.03 | .04 | .04 |
| **PCR.R** | .09** | .03 | .17** | .03 | -.04 | .02 | .02 |
| **PLIC.L** | .00 | -.06 | -.03 | .05 | -.06 | -.14 | .19** |
| **PLIC.R** | .04 | -.02 | .01 | .13 | -.11 | -.07 | .14* |
| **PTR.L** | .09** | .07 | .21*** | -.04 | -.10 | -.01 | .07 |
| **PTR.R** | .06* | .04 | .13* | -.05 | -.02 | .00 | -.01 |
| **RLIC.L** | .09** | .06 | .12* | .05 | .07 | .06 | .14* |
| **RLIC.R** | .09** | -.01 | .17** | -.01 | .06 | .01 | .13 |
| **SCR.L** | .06* | .03 | .13* | -.07 | .02 | -.09 | .08 |
| **SCR.R** | .05* | .03 | .09 | -.09 | -.05 | -.07 | .12 |
| **SFO.L** | .03 | .03 | .03 | -.05 | .00 | -.16* | .08 |
| **SFO.R** | .070* | .03 | .11 | -.04 | .00 | -.09 | .13 |
| **SLF.L** | .07** | .03 | .11 | -.02 | .03 | .02 | .09 |
| **SLF.R** | .06* | .02 | .10 | .02 | -.07 | -.01 | .10 |
| **SS.L** | .11*** | .18** | .139* | -.10 | -.03 | .00 | .11 |
| **SS.R** | .11*** | .03 | .24*** | -.12 | .07 | .01 | .07 |
| **TAP.L** | .13*** | .14* | .29*** | -.04 | -.05 | .03 | .02 |
| **TAP.R** | .09** | .05 | .20** | .09 | -.02 | .00 | -.05 |
| **UNC.L** | .05 | .10 | .03 | -.12 | -.05 | .23** | -.06 |
| **UNC.R** | .08** | .03 | .12* | -.12 | .13 | -.01 | .12 |

**Supplementary Table 38. Relationship between RD and age of disease onset by syndrome**. Partial correlations between RD in each ROI and age of disease onset controlling for age, age$^2$ and sex. Results are split by syndrome: all syndromes (All), temporal lobe epilepsy with hippocampal sclerosis in the left and right hemisphere (TLE-HS-l and TLE-HS-r respectively), non-lesional temporal lobe epilepsy in the left and right hemisphere (TLE-NL-l and TLE-NL-r respectively), genetic generalized epilepsy (GGE) and extra temporal (Extra). ROIs are separated by left (.L) and right (.R) hemisphere where indicated. Significance was set at *** $p$ =< 0.001 (controlling for multiple comparisons, one sided). * $p$ < 0.05 ** $p$ < 0.01 *** $p$ =< 0.001.

| ROI | All | TLE-HS-l | TLE-HS-r | TLE-NL-l | TLE-NL-r | GGE | ExE |
|---|---|---|---|---|---|---|---|
| **AverageRD** | -.18*** | -.15** | -.25*** | -.14* | -.18* | -.05 | -.02 |
| **BCC** | -.17*** | -.12* | -.24*** | -.15* | -.19* | .03 | -.03 |
| **GCC** | -.13*** | -.16*** | -.14* | -.05 | -.11 | .01 | -.03 |
| **SCC** | -.14*** | -.08 | -.27*** | -.02 | -.16 | -.06 | -.08 |
| **ACR.L** | -.13*** | -.17*** | -.12* | .00 | -.14 | .00 | -.05 |
| **ACR.R** | -.11*** | -.07 | -.09 | -.06 | -.16* | .01 | -.07 |
| **ALIC.L** | -.09*** | -.09 | -.11* | -.04 | -.03 | .07 | -.06 |
| **ALIC.R** | -.12*** | -.08 | -.10 | -.11 | -.08 | -.03 | -.11 |
| **CGC.L** | -.14*** | -.14** | -.11* | -.10 | -.06 | -.05 | -.04 |
| **CGC.R** | -.13*** | -.04 | -.14* | -.14* | -.11 | -.12 | -.05 |
| **CGH.L** | -.13*** | -.14** | -.18** | -.06 | .09 | -.03 | -.07 |
| **CGH.R** | -.12*** | -.02 | -.24*** | -.13 | -.05 | -.09 | .00 |
| **CST.L** | -.05* | -.12* | -.10 | -.04 | .08 | .01 | -.01 |
| **CST.R** | -.04 | -.05 | -.08 | -.04 | .06 | .03 | -.02 |
| **EC.L** | -.14*** | -.14** | -.15* | -.08 | -.08 | -.09 | -.01 |
| **EC.R** | -.14*** | -.12* | -.20** | -.06 | -.06 | -.10 | -.08 |
| **FX.ST.L** | -.12*** | -.10* | -.11* | -.21** | -.01 | .01 | -.02 |
| **FX.ST.R** | -.13*** | -.07 | -.18** | -.17* | -.08 | -.05 | -.03 |
| **PCR.L** | -.10 | -.02 | -.16** | -.03 | -.08 | -.10 | -.03 |
| **PCR.R** | -.11*** | -.08 | -.19** | .02 | -.07 | -.07 | -.05 |
| **PLIC.L** | -.05 | .03 | -.09 | -.06 | -.02 | .07 | -.13 |
| **PLIC.R** | -.08** | -.01 | -.10 | -.11 | .01 | -.04 | -.07 |
| **PTR.L** | -.12*** | -.10* | -.20** | -.03 | -.11 | -.05 | -.06 |
| **PTR.R** | -.11*** | -.10 | -.15* | .03 | -.14 | -.03 | -.02 |
| **RLIC.L** | -.13*** | -.10* | -.14* | -.10 | -.12 | -.08 | -.14* |
| **RLIC.R** | -.11*** | -.02 | -.17** | -.04 | -.09 | -.04 | -.06 |
| **SCR.L** | -.07* | -.06 | -.10 | .05 | -.09 | .06 | -.10 |
| **SCR.R** | -.08** | -.07 | -.07 | .03 | .05 | .01 | -.15* |
| **SFO.L** | -.09** | -.07 | -.11 | .00 | -.07 | .12 | -.04 |
| **SFO.R** | -.11*** | -.04 | -.15* | -.07 | .02 | .01 | -.10 |
| **SLF.L** | -.11*** | -.08 | -.15* | -.03 | .05 | -.05 | -.05 |
| **SLF.R** | -.10*** | -.03 | -.14* | -.04 | .02 | -.08 | -.07 |
| **SS.L** | -.14*** | -.17** | -.17** | -.01 | -.05 | -.05 | -.10 |
| **SS.R** | -.12*** | -.08 | -.21*** | .05 | -.16 | -.06 | -.02 |
| **TAP.L** | -.14*** | -.11* | -.24*** | .03 | -.16* | -.03 | -.01 |
| **TAP.R** | -.13*** | -.07 | -.19** | -.14* | -.21* | -.04 | .03 |
| **UNC.L** | -.07* | -.09 | -.05 | .04 | .03 | -.23** | .07 |
| **UNC.R** | -.12*** | -.05 | -.21*** | .02 | -.15 | -.02 | -.05 |

**Supplementary Table 39. Relationship between RD and disease duration by syndrome.** Partial correlations between RD in each ROI and disease duration controlling for age, age$^2$ and sex. Results are split by syndrome: all syndromes (All), temporal lobe epilepsy with hippocampal sclerosis in the left and right hemisphere (TLE-HS-l and TLE-HS-r respectively), non-lesional temporal lobe epilepsy in the left and right hemisphere (TLE-NL-l and TLE-NL-r respectively), genetic generalized epilepsy (GGE) and extra temporal (Extra). ROIs are separated by left (.L) and right (.R) hemisphere where indicated. Significance was set at *** $p$ =< 0.001 (controlling for multiple comparisons, one sided). * $p$ < 0.05 ** $p$ < 0.01 *** $p$ =< 0.001.

| ROI | All | TLE-HS-l | TLE-HS-r | TLE-NL-l | TLE-NL-r | GGE | ExE |
|---|---|---|---|---|---|---|---|
| **AverageRD** | .19*** | .17** | .27*** | .15* | .16 | .04 | .05 |
| **BCC** | .18*** | .14* | .24*** | .21** | .15 | -.04 | .04 |
| **GCC** | .15*** | .21** | .18** | .00 | .08 | -.02 | .04 |
| **SCC** | .15*** | .09 | .29*** | .04 | .05 | .05 | .11 |
| **ACR.L** | .14*** | .19** | .18** | -.03 | .11 | -.01 | .06 |
| **ACR.R** | .11*** | .09 | .13* | .01 | .16 | -.03 | .06 |
| **ALIC.L** | .07* | .07 | .05 | -.01 | .06 | -.09 | .09 |
| **ALIC.R** | .09** | .08 | .08 | .04 | .07 | .01 | .11 |
| **CGC.L** | .14*** | .15** | .12* | .04 | .10 | .03 | .06 |
| **CGC.R** | .14*** | .04 | .18** | .10 | .11 | .10 | .07 |
| **CGH.L** | .11*** | .08 | .18** | .02 | -.06 | .03 | .08 |
| **CGH.R** | .12*** | -.02 | .28*** | .10 | .13 | .09 | .02 |
| **CST.L** | .03 | .09 | .13* | .00 | -.14 | -.01 | .01 |
| **CST.R** | .01 | .04 | .08 | .00 | -.16 | -.02 | .01 |
| **EC.L** | .14*** | .14* | .15* | .04 | .11 | .07 | .04 |
| **EC.R** | .14*** | .09 | .21*** | .04 | .10 | .09 | .12 |
| **FX.ST.L** | .10*** | .10 | .11 | .15* | .01 | -.01 | .02 |
| **FX.ST.R** | .14*** | .07 | .17** | .12 | .15 | .03 | .07 |
| **PCR.L** | .10** | .03 | .20** | -.02 | -.01 | .09 | .05 |
| **PCR.R** | .11*** | .07 | .21*** | -.01 | -.03 | .05 | .06 |
| **PLIC.L** | .02 | -.04 | .07 | -.05 | -.01 | -.08 | .15* |
| **PLIC.R** | .06* | .01 | .09 | .09 | -.06 | .02 | .06 |
| **PTR.L** | .13*** | .12* | .24*** | .02 | .02 | .05 | .07 |
| **PTR.R** | .11*** | .11* | .18** | -.06 | .12 | .03 | .01 |
| **RLIC.L** | .10*** | .06 | .16* | .02 | .05 | .07 | .13 |
| **RLIC.R** | .11*** | .04 | .17** | .02 | .08 | .03 | .06 |
| **SCR.L** | .07* | .06 | .13* | -.08 | .08 | -.08 | .07 |
| **SCR.R** | .08** | .07 | .08 | -.08 | -.03 | -.02 | .13 |
| **SFO.L** | .07* | .05 | .07 | -.03 | .06 | -.13 | .05 |
| **SFO.R** | .10*** | .06 | .11 | .06 | .04 | -.02 | .07 |
| **SLF.L** | .12*** | .08 | .17** | .04 | .05 | .04 | .09 |
| **SLF.R** | .10*** | .00 | .16* | .04 | .03 | .07 | .09 |
| **SS.L** | .15*** | .19*** | .17** | -.03 | .13 | .05 | .12 |
| **SS.R** | .14*** | .12* | .22*** | -.06 | .22* | .05 | .04 |
| **TAP.L** | .14*** | .12* | .32*** | -.11 | .03 | .04 | .08 |
| **TAP.R** | .13*** | .07 | .25*** | .07 | .08 | .04 | .01 |
| **UNC.L** | .07* | .12* | .05 | -.11 | -.02 | .24** | -.06 |
| **UNC.R** | .11*** | .00 | .20** | -.06 | .25** | .01 | .09 |

**Supplementary Table 40. Relationship between AD and age of disease onset by syndrome**. Partial correlations between AD in each ROI and age of disease onset controlling for age, age$^2$ and sex. Results are split by syndrome: all syndromes (All), temporal lobe epilepsy with hippocampal sclerosis in the left and right hemisphere (TLE-HS-l and TLE-HS-r respectively), non-lesional temporal lobe epilepsy in the left and right hemisphere (TLE-NL-l and TLE-NL-r respectively), genetic generalized epilepsy (GGE) and extra temporal (Extra). ROIs are separated by left (.L) and right (.R) hemisphere where indicated. Significance was set at *** $p =< 0.001$ (controlling for multiple comparisons, one sided). * $p < 0.05$ ** $p < 0.01$ *** $p =< 0.001$.

| ROI | All | TLE-HS-l | TLE-HS-r | TLE-NL-l | TLE-NL-r | GGE | ExE |
|---|---|---|---|---|---|---|---|
| **AverageAD** | -.05* | -.03 | -.16** | -.01 | .09 | .05 | .03 |
| **BCC** | -.01 | .01 | .00 | -.03 | -.02 | .01 | .02 |
| **GCC** | -.02 | -.05 | .04 | -.03 | .06 | .08 | .00 |
| **SCC** | -.06* | -.08 | -.11 | .02 | -.07 | -.08 | .17* |
| **ACR.L** | -.02 | -.04 | -.01 | .04 | .10 | .06 | -.02 |
| **ACR.R** | -.04 | -.02 | -.08 | .04 | .11 | .09 | -.10 |
| **ALIC.L** | -.02 | -.04 | .01 | .00 | .11 | -.04 | -.07 |
| **ALIC.R** | -.01 | -.04 | .04 | .05 | .22* | .01 | -.13 |
| **CGC.L** | .05 | .03 | .02 | .04 | .16 | -.10 | -.01 |
| **CGC.R** | .05 | .12* | .03 | .00 | .16 | -.04 | -.10 |
| **CGH.L** | .00 | -.02 | -.04 | .04 | .14 | .09 | .00 |
| **CGH.R** | -.02 | -.01 | -.13* | .03 | .19* | .02 | -.02 |
| **CST.L** | .02 | .03 | -.10 | .05 | .15 | .02 | .04 |
| **CST.R** | .04 | .11* | -.09 | .12 | .17* | -.01 | .07 |
| **EC.L** | .01 | .01 | .02 | .02 | .19* | .02 | -.05 |
| **EC.R** | .03 | .05 | .06 | .03 | .26** | .05 | -.14* |
| **FX.ST.L** | -.03 | .00 | -.09 | -.10 | .05 | .05 | -.01 |
| **FX.ST.R** | -.02 | .13* | -.12* | -.02 | .04 | .12 | -.12 |
| **PCR.L** | -.04 | -.04 | -.05 | .04 | -.12 | .02 | .02 |
| **PCR.R** | -.02 | .01 | -.04 | -.04 | -.06 | .01 | .06 |
| **PLIC.L** | .00 | -.03 | .05 | -.08 | .05 | .14 | -.09 |
| **PLIC.R** | -.01 | .00 | .02 | -.07 | .02 | .10 | -.04 |
| **PTR.L** | .00 | .01 | -.12* | .13 | .12 | .09 | -.02 |
| **PTR.R** | .02 | .05 | -.05 | .08 | .08 | .05 | .02 |
| **RLIC.L** | -.03 | -.04 | -.01 | -.06 | -.02 | -.01 | .00 |
| **RLIC.R** | -.04 | .06 | -.11* | -.09 | .02 | .02 | -.11 |
| **SCR.L** | -.01 | -.03 | -.03 | .02 | .09 | .10 | -.01 |
| **SCR.R** | -.02 | .01 | -.04 | -.04 | .07 | .13 | -.03 |
| **SFO.L** | .01 | .03 | -.01 | .01 | .09 | .14 | -.08 |
| **SFO.R** | .01 | .05 | -.02 | .04 | .17* | .13 | -.12 |
| **SLF.L** | .02 | -.01 | .03 | .06 | .16 | .00 | -.02 |
| **SLF.R** | .06* | .00 | .09 | .03 | .24** | .13 | -.02 |
| **SS.L** | -.03 | -.10* | -.05 | .10 | .15 | .04 | .00 |
| **SS.R** | -.05 | .02 | -.17** | -.01 | .08 | .07 | -.06 |
| **TAP.L** | -.04 | -.08 | -.15* | .02 | .02 | -.04 | .16* |
| **TAP.R** | -.01 | .01 | -.05 | -.11 | -.06 | .01 | .14* |
| **UNC.L** | .00 | -.06 | .04 | .08 | .11 | -.09 | .00 |
| **UNC.R** | .00 | .00 | -.05 | .12 | .21* | .09 | -.07 |

**Supplementary Table 41. Relationship between AD and disease duration by syndrome**. Partial correlations between AD in each ROI and disease duration controlling for age, $age^2$ and sex. Results are split by syndrome: all syndromes (All), temporal lobe epilepsy with hippocampal sclerosis in the left and right hemisphere (TLE-HS-l and TLE-HS-r respectively), non-lesional temporal lobe epilepsy in the left and right hemisphere (TLE-NL-l and TLE-NL-r respectively), genetic generalized epilepsy (GGE) and extra temporal (Extra). ROIs are separated by left (.L) and right (.R) hemisphere where indicated. Significance was set at *** $p =< 0.001$ (controlling for multiple comparisons, one sided). * $p < 0.05$ ** $p < 0.01$ *** $p =< 0.001$.

| ROI | All | TLE-HS-l | TLE-HS-r | TLE-NL-l | TLE-NL-r | GGE | ExE |
|---|---|---|---|---|---|---|---|
| **AverageAD** | .03 | -.02 | .12* | -.04 | -.13 | -.06 | .02 |
| **BCC** | -.01 | -.04 | -.04 | -.01 | -.06 | -.01 | .01 |
| **GCC** | .02 | .06 | -.03 | -.06 | -.16 | -.09 | .06 |
| **SCC** | .05 | .03 | .09 | -.03 | .00 | .08 | -.08 |
| **ACR.L** | .03 | .04 | .06 | -.08 | -.08 | -.07 | .03 |
| **ACR.R** | .04 | .02 | .11 | -.09 | -.16 | -.10 | .12 |
| **ALIC.L** | -.01 | .01 | -.10 | -.01 | -.12 | .03 | .11 |
| **ALIC.R** | -.02 | .02 | -.13* | -.08 | -.22* | -.03 | .19* |
| **CGC.L** | -.06* | -.07 | -.05 | -.04 | -.07 | .09 | .02 |
| **CGC.R** | -.04 | -.10 | -.03 | .01 | -.14 | .03 | .14* |
| **CGH.L** | -.03 | -.05 | .02 | -.06 | -.10 | -.10 | .01 |
| **CGH.R** | .00 | -.04 | .11 | -.04 | -.10 | -.03 | .02 |
| **CST.L** | -.02 | -.08 | .14* | -.04 | -.19* | -.01 | .00 |
| **CST.R** | -.04 | -.09 | .07 | -.13 | -.22* | .01 | .02 |
| **EC.L** | -.01 | -.02 | -.05 | -.01 | -.14 | -.04 | .10 |
| **EC.R** | -.03 | -.08 | -.07 | -.04 | -.18* | -.07 | .19** |
| **FX.ST.L** | .01 | -.04 | .09 | .08 | -.07 | -.07 | -.04 |
| **FX.ST.R** | .01 | -.13* | .10 | -.02 | .01 | -.14 | .09 |
| **PCR.L** | .05 | .05 | .10 | -.06 | -.04 | -.03 | .00 |
| **PCR.R** | .03 | .00 | .06 | .08 | -.02 | -.03 | -.05 |
| **PLIC.L** | .00 | .01 | -.09 | .13 | -.04 | -.14 | .10 |
| **PLIC.R** | .01 | .02 | -.07 | .08 | -.01 | -.12 | .07 |
| **PTR.L** | .00 | -.02 | .09 | -.11 | -.20* | -.08 | .04 |
| **PTR.R** | -.03 | -.06 | .04 | -.08 | -.16 | -.05 | -.04 |
| **RLIC.L** | .04 | .01 | .02 | .09 | .04 | .02 | .05 |
| **RLIC.R** | .04 | -.08 | .10 | .11 | -.03 | -.02 | .16* |
| **SCR.L** | .00 | -.01 | .01 | .02 | -.11 | -.10 | .04 |
| **SCR.R** | .01 | -.01 | .02 | .02 | -.03 | -.14 | .06 |
| **SFO.L** | -.04 | -.07 | -.05 | -.07 | -.01 | -.14 | .08 |
| **SFO.R** | -.03 | -.04 | -.05 | -.12 | -.12 | -.13 | .16* |
| **SLF.L** | -.02 | -.01 | -.02 | -.05 | -.06 | -.01 | .04 |
| **SLF.R** | -.06* | -.03 | -.08 | -.05 | -.17 | -.15 | .06 |
| **SS.L** | .03 | .08 | .04 | -.09 | -.08 | -.05 | .05 |
| **SS.R** | .04 | -.02 | .14* | -.09 | -.06 | -.08 | .11 |
| **TAP.L** | .05 | .09 | .19** | -.05 | -.11 | .04 | -.11 |
| **TAP.R** | -.02 | -.03 | .05 | .05 | -.16 | -.02 | -.14* |
| **UNC.L** | -.01 | .03 | -.06 | -.12 | -.05 | .08 | -.01 |
| **UNC.R** | -.02 | -.06 | .02 | -.08 | -.13 | -.10 | .06 |
